# Cells specify fate within an optimal window of positional information determined by morphogenesis

**DOI:** 10.64898/2026.06.23.734083

**Authors:** Chia-Teng Chang, Julian Renaud, Gasper Tkacik, Tony Yu-Chen Tsai

## Abstract

How cells sense position to adopt appropriate fates is a central problem in development. In a classic paradigm, cells read morphogen gradients encoding positional information (PI), with prolonged signal integration improving fate precision. However, how cells adapt this strategy in morphogenetic tissues remains unclear. Here, we reconstructed complete positional and signalling histories for individual cells during zebrafish neurulation, where the Sonic hedgehog (Shh) gradient patterns ventral progenitors as the neural plate folds into a tube. Despite steadily increasing Shh activity, Shh-encoded PI peaked early and then declined. Morphogenesis set this early readout window and imposed a ∼1.2-bit ceiling on Shh-encoded information about final position and fate. Fate mapping, transcriptomic analyses, and timed Shh inhibition showed that fate specification is temporally and functionally aligned with this early readout window. Thus, when morphogenesis decouples signal quality from signal strength, cells specify fate when signalling is most informative, not when signalling is strongest.

## Main

How cells sense their position within a tissue to adopt the right fates in the right places remains a central problem in cell and developmental biology. A classic paradigm posits that morphogen gradients encode positional information that cells decode to specify fate^1^. Landmark studies in the Drosophila blastoderm substantiated this paradigm by quantifying the precision of the Bicoid morphogen gradient and the reproducibility of downstream gene expression patterns^2–6^. In this syncytial and relatively static system, nuclei integrate morphogen levels over time into spatial profiles of transcription factors that encode positional information with near single-nucleus precision^7^. Snapshots of expression patterns of key transcription factors can be optimally decoded to predict the positions of downstream genes in wild type and across patterning mutants^8^. This success partially relied on a syncytial context in which nuclei share a common cytoplasm, a quasi-static geometry with minimal rearrangement, and an effectively one-dimensional axis.

In contrast, applying this paradigm to tissues undergoing morphogenesis faces non-trivial challenges. Morphogen sensing is inherently noisy, requiring temporal integration to improve precision^9–12^. However, as morphogenesis proceeds, cell rearrangements continuously alter relative positions, progressively decoupling accumulated signalling histories from cells’ current positions. As a result, prolonged signalling integration does not necessarily yield more information about where a cell will ultimately reside. How morphogenesis shapes the dynamics of morphogen-encoded positional information, and how cells make reliable fate decisions despite ongoing tissue reorganization, therefore remain key unanswered questions.

Vertebrate neurulation provides a direct testbed for this question because it couples tissue patterning with morphogenesis, two canonical processes often analysed separately. Sonic hedgehog (Shh) secreted from midline tissues (i.e., notochord and floor plate) forms a gradient that patterns neural progenitors^13,14^. At the same time, the neural plate converges toward the midline and folds into a neural tube, converting the original medial–lateral ordering of the plate into the ventral–dorsal ordering of the tube (Fig. 1a)^15,16^. Thus, cells are challenged to interpret the Shh gradient while cell rearrangements alter their positions relative to the Shh source at the midline.

**Fig. 1.**
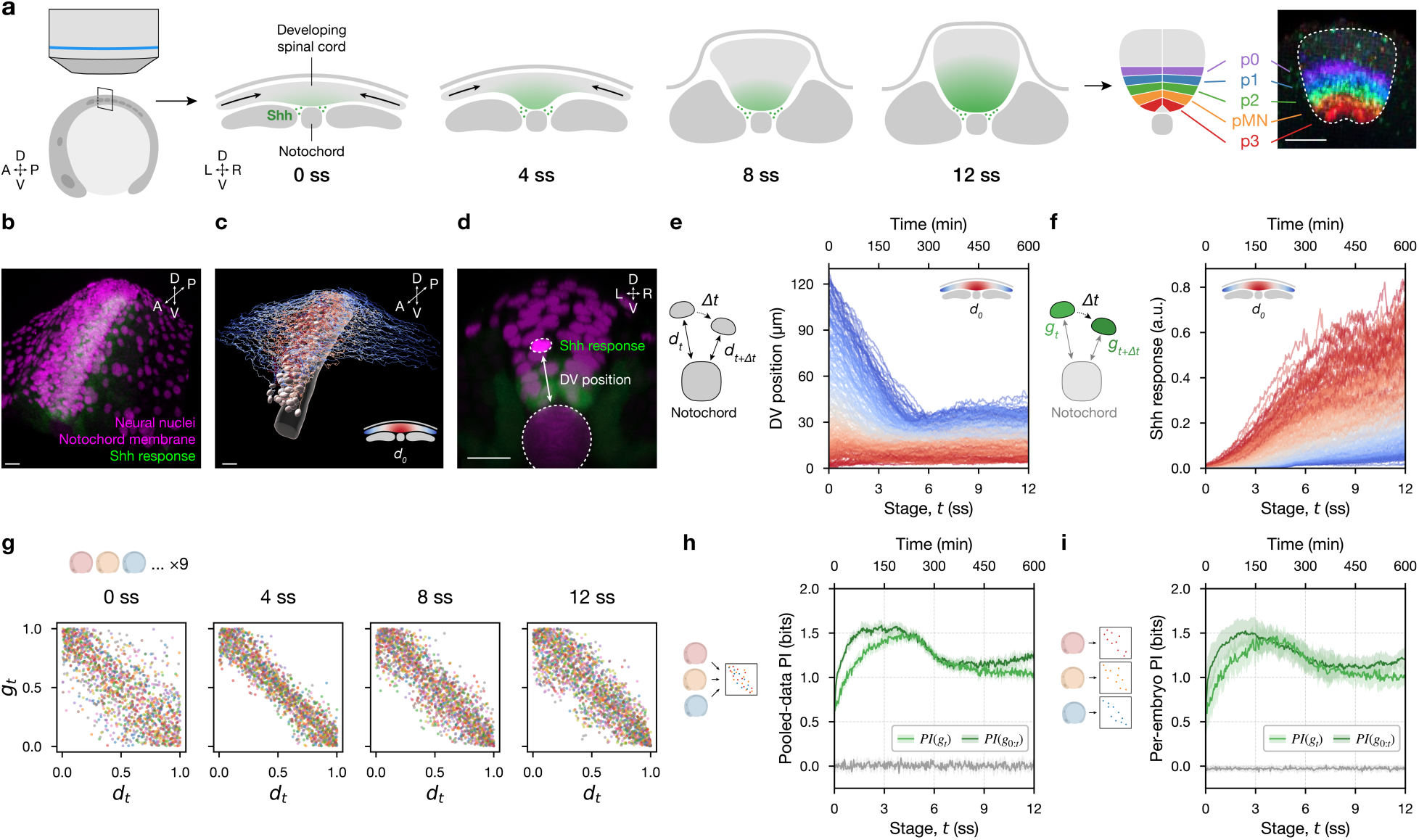
Lineage reconstruction of ventral neural progenitors reveals Shh-encoded positional information dynamics. **(A)** Schematic of zebrafish neurulation across 0–12 somite stages (ss). Shh is produced by midline tissues (notochord) and patterns ventral progenitor domains (p3–p0) by 12 ss as the neural plate folds into a tube. A, anterior; P, posterior; D, dorsal; V, ventral; L, left; R, right. Scale bar, 20 µm. **b,** Representative confocal z-stack at 12 ss (*Tg(sox19a:H2B-mCherry)*; *Tg(shha:mem-mCherry)*; *TgBAC(ptch2:Kaede)*) showing neural nuclei (mCherry, magenta), notochord membrane (mCherry, magenta), and Shh response (Kaede, green). Scale bar, 20 µm. **c,** Lineage reconstruction of ventral spinal cord progenitors from one wild-type embryo, coloured by initial DV position *d*_0_. **D,E,** DV position trajectories (**D**) and Shh response trajectories (**E**) for the lineages in **c** (n=221 lineages), colour-coded by *d*_0_. **F,** Relationship between within-embryo percentile-ranked Shh response *g*_*t*_ and DV position *d*_*t*_ at 0, 4, 8 and 12 ss, colour-coded by individual embryos (N=9 embryos; n=1,561 cells). **G,** Positional information between Shh response and DV position at each stage, computed from cells pooled across embryos after within-embryo rank normalization. Curves show mean with 95% CI (Supplementary Note 2). Grey curve, shuffle control. **H,** Positional information computed per embryo and summarized as mean ± s.d. across embryos (N=9).

Here we quantitatively connect patterning and morphogenesis by tracking ventral spinal cord progenitors through neurulation using *in toto* single-cell lineage tracking, measuring Shh response and position over time, and analysing these data using an information-theoretic framework. This approach allows us to ask, at each stage, how well a cell’s Shh signalling history predicts its past, current, and future position. We uncovered a simple organizing principle for patterning during morphogenesis: cells accumulate positional information by temporally integrating Shh signalling, but concurrent cell rearrangements progressively erode this information. Although Shh signalling strengthens over time, the opposing effect of cell rearrangement creates an early readout window in which positional information is maximized.

Analyses of fate specification dynamics also suggest that cells specify fate not when Shh signalling is strongest, but when it encodes maximal positional information. Together, our study demonstrates that morphogenesis is not merely a backdrop for tissue patterning, but an active determinant of the optimal information window which cells exploit for fate specification.

### Shh-encoded positional information peaks early during neurulation

We set out to obtain the full history of every cell’s Shh response and dorsoventral (DV) position in the ventral spinal cord throughout the morphogenetic transition from neural plate to neural tube, and the concurrent emergence of stereotypic stripe patterns of ventral progenitors (Fig. 1a). Our *in toto* live-imaging captured every cell within the four most ventral progenitor types (p3, pMN, p2, p1) in a 303 × 303 × 145 µm volume spanning somites 2–6, imaged every 2 min for 10 h at 23 °C (0–12 somite stages, ss; approximately 10–15 hpf equivalent at 28.5 °C). We developed deep-learning-based image analysis pipelines, achieving >99% segmentation accuracy and continuous, error-free tracking of individual spinal cord cells and the Shh-secreting notochord (Fig. 1b–f; Extended Data Fig. 1–2; Supplementary Methods 1–2; Supplementary Video 1)^14–16^. This high-fidelity segmentation and tracking pipeline allowed us to reconstruct complete lineages within the ventral spinal cord, yielding 1,561 lineages across 9 zebrafish embryos.

To quantify how well Shh response predicts instantaneous DV position, we measured morphogen-encoded positional information (in bits) as the mutual information between Shh response and DV position (Supplementary Note 1)^7,17^. Because this is the first *in vivo* positional information analysis based on single-cell, lineage-resolved data, we compared multiple mutual information estimators and included shuffle controls and finite-sample debiasing to guard against estimation artefacts (Extended Data Fig. 4; Supplementary Note 2–3)^18–20^. We benchmarked candidate transgenic reporters for Shh response and DV position metrics and selected the combination that maximized estimated positional information, defining Shh response as nuclear *TgBAC(ptch2:Kaede)* fluorescence intensity (a cumulative readout of Shh signalling over time)^21^and DV position as the shortest 3D distance from each nucleus to the notochord surface (Fig. 1d; Supplementary Note 4; Extended Data Fig. 3a–h,n–r).

We quantified positional information *PI*(*g*_*t*_) at each somite stage *t* by pooling cells across embryos after normalizing Shh response *g*_*t*_ and DV position *d*_*t*_ within each embryo (Fig. 1g,h; Supplementary Notes 1–2). Despite a monotonic increase in Shh response—both in cumulative *TgBAC(ptch2:Kaede)* signal and in an instantaneous response inferred from reporter transcripts (Fig. 1f; Extended Data Fig. 3j)—positional information rose to an early maximum at ∼4 ss and then declined to an intermediate level (light green curve in Fig. 1h). Thus, increasing Shh response does not guarantee increasing positional information.

Could this rise-and-fall of positional information be an artefact of reporter dynamics (e.g., memory, delay, smoothing, or bleaching), rather than reflecting the actual information available in Shh response? To address this concern, we asked whether the entire signalling history, rather than the snapshot reporter value alone, could yield additional information about DV position and explain this rise-and-fall. We therefore developed a history-based decoder to estimate trajectory-based positional information, *PI*(*g*_0:*t*_)^22^. Specifically, at each stage we trained a supervised decoder to predict each lineage’s current DV position *d*_*t*_ from its entire Shh response history *g*_0:*t*_, allowing the decoder to maximize positional inference by up-weighting more informative stages and down-weighting less informative ones (Supplementary Note 5; Extended Data Fig. 5a–e). This trajectory-based positional information extended the early high-information window towards earlier times (∼2 ss) but did not increase the peak information nor eliminate the subsequent decline (dark green curve in Fig. 1h).

To account for reporter delay, we placed a conservative upper bound on the effective delay of the *TgBAC(ptch2:Kaede)* readout using the Shh inhibitor cyclopamine. This yielded an estimated total delay of ∼1.6 ss, encompassing drug onset, pathway shutdown, and reporter fluorescence dynamics (Extended Data Fig. 3l,m). Importantly, the decline persisted even when positional information was estimated from Shh response histories extending to later stages (Extended Data Fig. 5i,j), a procedure that would recover additional information if later reporter values were more informative about current DV position. Thus, the decline in positional information is not explained by reporter delay. Because the estimated delay is short relative to the neurulation timescale, and because endogenous Shh targets also require time to be expressed and accumulate, we use the measured Shh response at each stage without an explicit delay correction in subsequent analyses.

Finally, this rise-and-fall pattern was observed consistently when positional information was computed separately within each embryo (Fig. 1i), indicating that the information dynamics are highly stereotypic across embryos. Together, these results establish—in a single-cell, lineage-resolved manner—an early readout window in which Shh-encoded positional information peaks during neurulation despite continuously increasing Shh response.

### Cell rearrangements define an upper bound of positional information

The decline in positional information could not be explained by Shh response kinetics alone (Fig. 1h), suggesting that changes in DV position progressively weakened the relationship between Shh response and DV position. To isolate the effect of changes in DV position, we defined positional persistence, *PP*_*t*’_(*d*_*t*_), as the mutual information (in bits) between DV position at stage *t* and DV position at stage *t*’, and estimated it across stages (Fig. 2a; Supplementary Note 1). Intuitively, high positional persistence indicates that DV position at stage *t* remains strongly predictive of DV position at stage *t*^′^, whereas low positional persistence indicates that DV position is effectively randomized across stages.

**Fig. 2.**
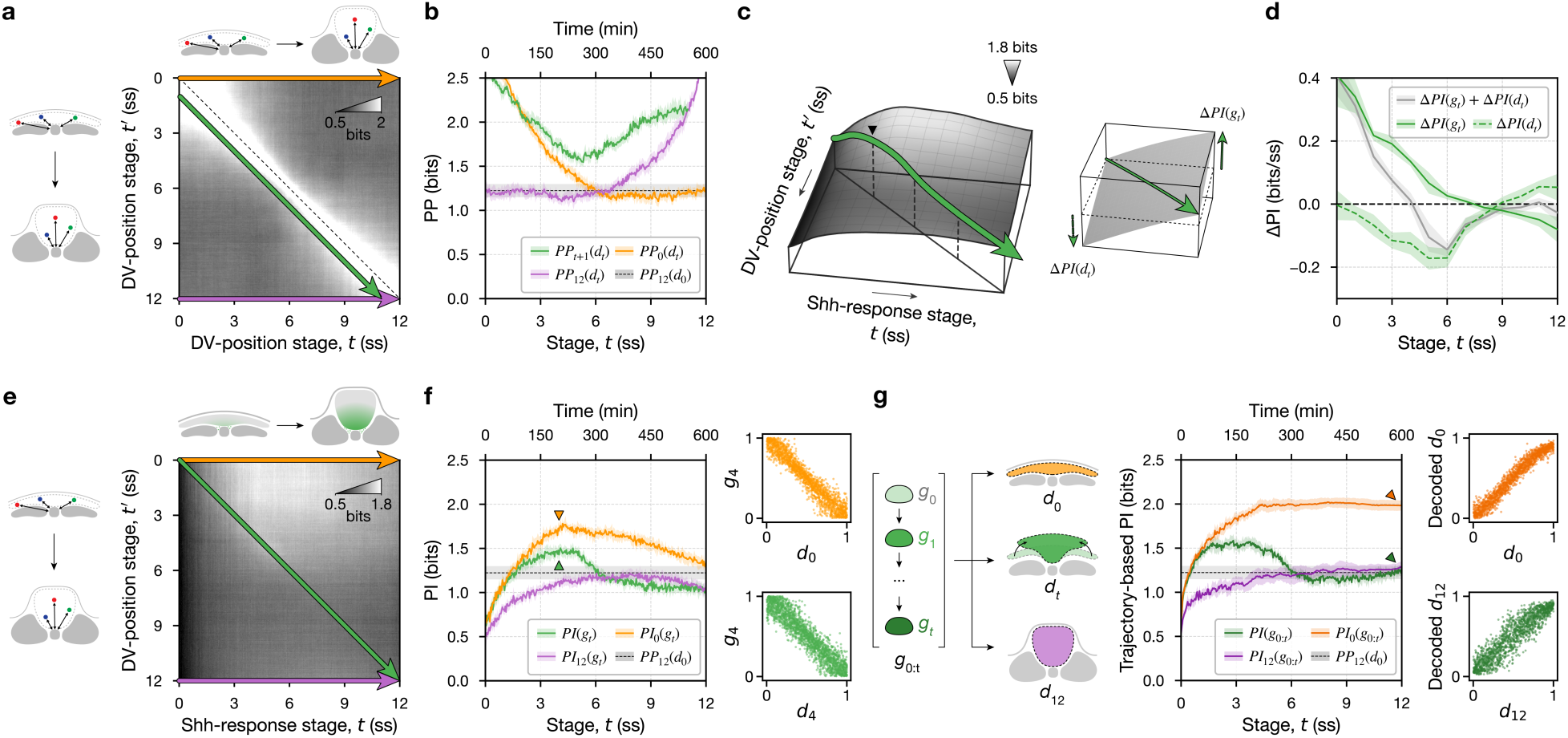
Cross-temporal positional persistence defines a ceiling on Shh-encoded positional information. **a,** Cross-temporal positional persistence map, *PP*_*t*_’(*d*_*t*_), quantifying how well DV position at stage *t* predicts DV position at stage *t*′ (Supplementary Note 1). **b,** Summary slices from **a**: positional persistence over one-somite-stage intervals, *PP*_*t*+1_(*d*_*t*_) (green); relative to initial DV position *PP*_0_(*d*_*t*_) (orange); and relative to final DV position, *PP*_12_(*d*_*t*_) (purple). Dashed line, *PP*_12_(*d*_0_), indicating the ceiling on information about final DV position that can be preserved from the initial DV position through neurulation under cell rearrangements. Curves show mean with 95% CI. **c,** Schematic for decomposing the stage-to-stage change in positional information into a contribution from changes in Shh response, Δ*PI*(*g*_*t*_), and a contribution from changes in DV position, Δ*PI*(*d*_*t*_) (Supplementary Note 6). **d,** Δ*PI*(*g*_*t*_) and Δ*PI*(*d*_*t*_) across stages (Supplementary Note 6). Values are binned by somite stage; curves show mean with 95% CI. **e,** Cross-temporal positional information map, *PI*_*t*′_(*g*_*t*_) quantifying how well Shh response at stage *t* predicts DV position at stage *t*′ (Supplementary Notes 1). **f,** Summary slices from **e**: positional information about DV position at the same stage, *PI*(*g*_*t*_) (green); about initial DV position, *PI*_0_(*g*_*t*_) (orange); and about final DV position *PI*_12_(*g*_*t*_) (purple). Dashed line, *PP*_12_(*d*_0_) from **b**. Insets, Shh response at 4 ss plotted against DV position at 0 ss or 4 ss. Curves show mean with 95% CI. **g,** Trajectory-based positional information computed from Shh response histories, *PI*_*t*′_(*g*_0:*t*_), plotted as in **f** (Supplementary Note 5). Insets, decoded DV position from *g*_0:*t*_ versus measured DV position at 0 ss or 12 ss.

Positional persistence between consecutive somite stages, *PP*_*t*+1_(*d*_*t*_), fell to a minimum around∼5 ss (green curve in Fig. 2b), indicating the most extensive rearrangements of DV position, and then increased at later stages as cell rearrangements slowed down. Next, we decomposed the stepwise gain or loss of positional information at each time point into orthogonal contributions from signal accumulation, *ΔPI*(*g*_*t*_), and cell rearrangement, *ΔPI*(*d*_*t*_) (Fig. 2c,d; Extended Data Fig. 5p–t; Supplementary Note 6). Early in neurulation, changes in Shh response increased positional information (*ΔPI*(*g*_*t*_) > 0), but this contribution weakened over time. Cell rearrangements, by contrast, continuously eroded positional information (*ΔPI*(*d*_*t*_) < 0), with their detrimental effect growing stronger until ∼6 ss. Before ∼4 ss, the positive contribution from Shh response accumulation dominated, whereas after ∼4 ss, the negative contribution from cell rearrangement took over, driving the overall decline in positional information (Fig. 2d; Extended Data Fig. 5p-t). The early optimal readout window for positional information, therefore, resulted from the competing effects of signal integration and cell rearrangements.

We next asked how well, in principle, the cell’s DV position at earlier stages can predict its final DV position. Positional persistence between stage *t* and 12 ss, *PP*_12_(*d*_*t*_), remained near ∼1.2 bits from 0–6 ss (purple curve in Fig. 2b), defining a ceiling imposed by cell rearrangements on how much information about the final DV position can be preserved from earlier DV positions. Thus, even if Shh response perfectly encoded early position, at most ∼1.2 bits could be preserved at the final stage. We therefore hypothesized that although Shh-encoded positional information *PI*(*g*_*t*_) reached a peak of ∼1.5 bits at ∼4 ss (Fig. 1h), no more than 1.2 bits could be transferred to the final stage.

To test this, we computed cross-temporal positional information, *PI*_*t*’_(*g*_*t*_), which measures how well Shh response at stage *t* predicts DV position at stage *t*′ (Fig. 2e; Supplementary Notes 1). Positional information about the initial DV position, *PI*_0_(*g*_*t*_), rose to a maximum near ∼4 ss and then declined (orange curves in Fig. 2f). Interestingly, at any given time, Shh responses always carried more information about the initial DV position, *PI*_0_(*g*_*t*_), than about the current DV position, *PI*(*g*_*t*_) (orange versus green curves in Fig. 2f), and this ordering persisted even when DV positions were decoded from the full Shh response history (dark orange versus dark green in Fig. 2g; Extended Data Fig. 5a–e). By contrast, positional information about the final DV position, *PI*_12_(*g*_*t*_), increased from 0–6 ss, approached the ∼1.2-bit ceiling, and then declined after ∼9 ss (purple curve in Fig. 2f). Even when the full Shh response history was available, positional information about the final DV position remained capped near this ceiling (dark purple curve in Fig. 2g).

Taken together, these results support a model in which Shh response serves as a cellular “memory” of early DV position. Although alternative interpretations cannot be excluded, cross-temporal analyses show that cell rearrangements impose an explicit ceiling on information about final DV position, which Shh-encoded positional information can approach but not exceed.

### Morphogenesis defines the timing of optimal positional information

Cross-temporal information analyses suggested that the timing of maximal positional information depends on cell rearrangement dynamics. We therefore hypothesized that altering the pace of neural plate convergence could shift both the timing of the positional persistence minimum and the timing of the positional information maximum.

Zebrafish neural plate convergence is controlled by opposing adhesion systems, in which Cdh2-dependent cell–cell adhesion promotes convergence whereas Itga5-dependent cell–ECM adhesion resists it (Fig. 3a)^23^. We analysed maternal–zygotic *itga5^-/-^* (*MZitga5^-/-^*) embryos with faster neural plate convergence and *cdh2^-/-^* embryos with slower neural plate convergence (Fig. 3b,c; Supplementary Video 2). In these mutant embryos, the timing of positional persistence minimum—and therefore the maximal DV position rearrangements—shifted to earlier stages with faster convergence and to later stages with slower convergence (Fig. 3d). The maximum of Shh-encoded positional information shifted in parallel, plateauing earlier with faster convergence and peaking later with slower convergence (Fig. 3e).

**Fig. 3.**
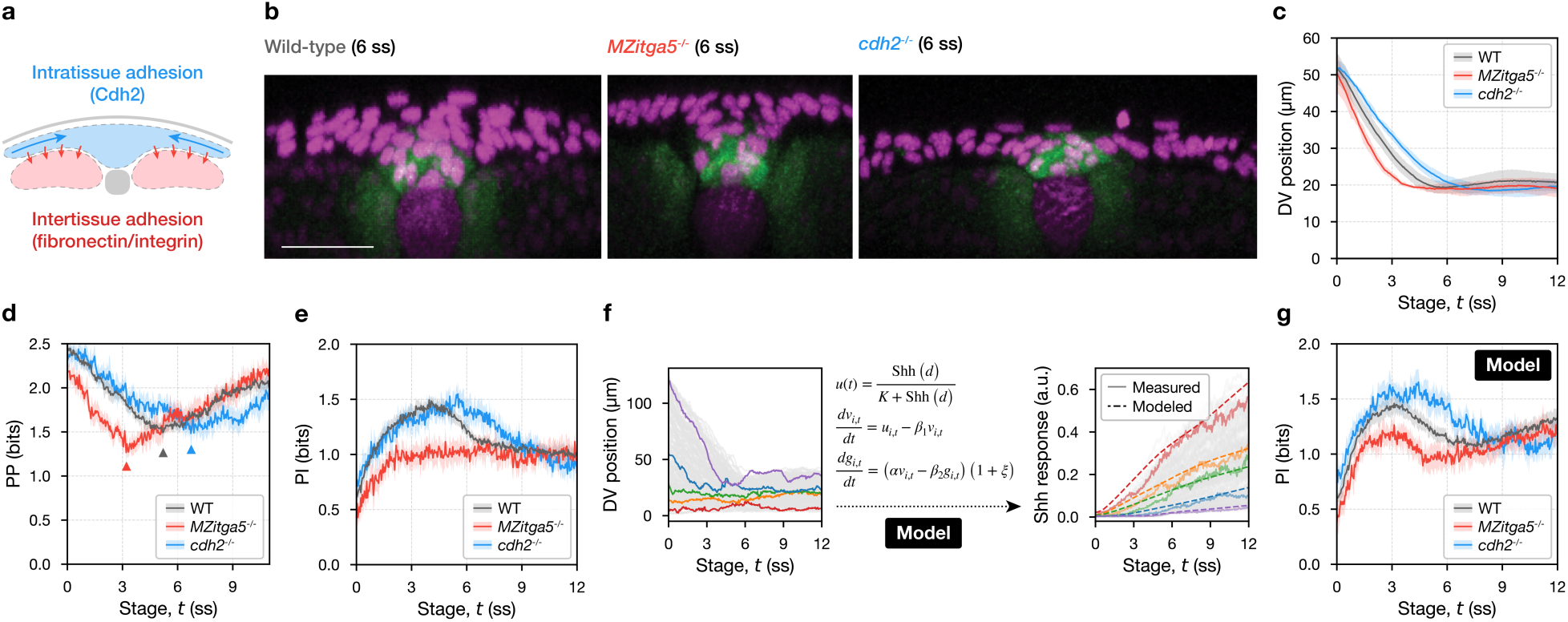
Genetic perturbations of neural plate convergence shift DV position dynamics and Shh-encoded positional information. **a,** Schematic of adhesion systems regulating neural plate convergence: Cdh2-mediated cell–cell adhesion promotes convergence, whereas fibronectin–integrin adhesion to the extracellular matrix opposes convergence. **b,** Representative transverse sections at 6 ss from wild type, *MZitga5^-/-^* and *cdh2^-/-^*embryos. Magenta, *Tg(sox19a:H2B-mCherry); Tg(shha:mem-mCherry)*; green, *TgBAC(ptch2:Kaede)*. Scale bar, 20 µm. **c,** DV position over time for each genotype. For each embryo, DV position was averaged across tracked ventral spinal-cord progenitors at each stage and then summarized as mean ± s.d. across embryos. **d,** Positional persistence between consecutive somite stages, *PP*_*t*+1_(*d*_*t*_), for each genotype (mean with 95% CI). Triangles indicate the stage of the minimum. **e,** Positional information for each genotype (mean with 95% CI). **f,** Schematic of the hybrid model (Supplementary Note 8). Empirically measured DV position trajectories are used as inputs (left). Right, measured Shh response and model-predicted Shh response. **g,** Positional information computed from model-predicted Shh responses driven by the measured DV position trajectories for wild type, *MZitga5^-/-^* and *cdh2^-/-^*. Mean with 95% CI.

To gain quantitative insight, we normalized morphogenesis progression across genotypes by rescaling developmental time (Supplementary Note 7). On this normalized timescale, positional information in *cdh2^-/-^*embryos was closely aligned with wild type, whereas *MZitga5^-/-^* embryos showed only partial alignment. Instead of a pronounced peak at the rescaled 4 ss, positional information in *MZitga5^-/-^* embryos plateaued around the rescaled 4 ss at ∼0.4 bits below wild type (Extended Data Fig. 6e–h). Because Shh response at any given stage was most informative about initial DV position, we interpret this plateau as reflecting earlier rearrangement of initial DV position in *MZitga5^-/-^* embryos (Extended Data Fig. 6c), which reduced the amount of early positional information that could be accumulated (Extended Data Fig. 6a).

To test whether cell movements alone can explain the positional information dynamics independently of changes in Shh signalling, we developed hybrid models to perform a controlled sufficiency test. These models are “hybrid” in the sense that we replayed each cell’s experimentally measured trajectory through a simulated Shh gradient and modelled cellular Shh response, assuming all cells shared identical dose-response dynamics upon Shh ligand exposure (Fig. 3f; Supplementary Note 8). We benchmarked multiple response-module variants and obtained consistent conclusions across models (Extended Data Fig. 7). We therefore focused on one simple best-performing model, whose parameters were fit once on wild-type embryos and then held fixed to predict Shh responses in all cells across genotypes without refitting. When driven by the DV position trajectories measured in wild-type, *MZitga5^-/-^*, and *cdh2^-/-^* embryos, the hybrid model reproduced the genotype-specific shifts in the timing of maximal positional information (Fig. 3g). Therefore, cell movement dynamics alone are sufficient to set the timing of the Shh-encoded positional information optimum.

### Notch-mediated heterogeneity prevents late accumulation of positional information

Despite the overall agreement between the hybrid model and the measurements, a key discrepancy remained. Across all response-module variants, positional information was predicted to increase again once DV position had stabilized (Fig. 3g; Extended Data Fig. 7b,e,g), which was not observed experimentally (Fig. 3e).

Because the model assumed a fixed, homogeneous Shh response rule across cells, we hypothesized that late-emerging cell-to-cell heterogeneity in Shh sensitivity could account for this discrepancy. Consistent with our hypothesis, we observed a negative contribution of positional information by late Shh response accumulation (*ΔPI*(*g*_*t*_) < 0) after ∼8ss (Fig. 2d). Additionally, positional information about the final DV position, *PI*_12_(*g*_*t*_), declined from its 1.2-bit ceiling after 8ss (Fig. 2f).

To compare early versus late Shh response directly, we photoconverted *TgBAC(ptch2:Kaede)* at 8 ss and imaged embryos at 12 ss, separating reporter signals accumulated before 8 ss (magenta) from those accumulated between 8 and 12 ss, defined here as late Shh response (cyan). Strikingly, the late Shh response displayed a pronounced salt-and-pepper pattern among neighbouring cells (Fig. 4a), reminiscent of patterns produced by lateral inhibition^24,25^. The late Shh response was also systematically lower in more lateral regions of the spinal cord (Fig. 4b), in line with the medial–lateral organization in which progenitor-like states are enriched medially and more neurogenic states laterally (Extended Data Fig. 8d)^26,27^. Together, these spatial signatures suggested that late Shh response became heterogeneous across cells during neurulation.

**Fig. 4.**
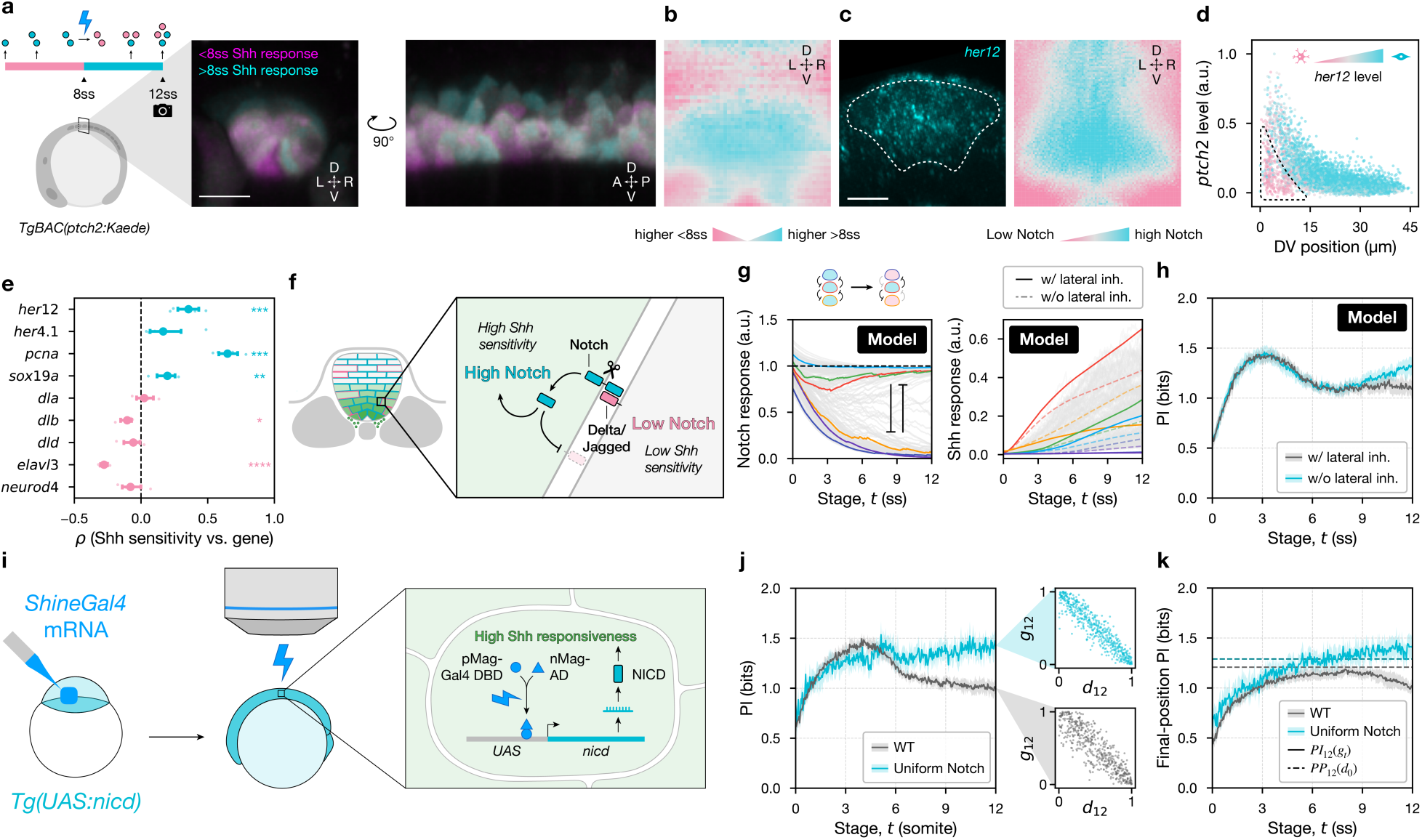
Notch activity drives heterogeneity in Shh sensitivity and limits positional information at late stages. **a,** *TgBAC(ptch2:Kaede)* embryos photoconverted at 8 ss to separate Shh response accumulated before 8 ss (magenta) from Shh response accumulated between 8 and 12 ss (late Shh response; cyan). Transverse section (middle) and sagittal section (right) at 12 ss. Scale bar, 20 µm. **b,** Transverse DV–LR mean profile of the within-embryo percentile difference between late and early Shh response (N = 3 embryos). **c,** Transverse section and DV–LR mean profile of *her12* mRNA at 10 ss (N = 6 embryos). Scale bar, 20 µm. **d,** DV position versus instantaneous Shh response measured by endogenous *ptch2* mRNA, coloured by Notch activity measured by endogenous *her12* mRNA at 10 ss. Data pooled from N=6 embryos (n=5,561 cells); medial floor plate excluded. **e,** Embryo-level Spearman correlations between inferred Shh sensitivity and expression of indicated Notch-regulated genes. Points, embryos; error bars, s.d. across embryos. Correlations were tested against zero using one-sample t-tests. **F,** Schematic of the hybrid model with Notch lateral inhibition (Supplementary Note 8). Left, neighbours updated over time from measured 3D trajectories; Notch lateral inhibition between dynamically exchanging neighbours generates effectively bimodal Notch activity that modulates Shh sensitivity. Right, corresponding modelled Shh response dynamics. **h,** Positional information from model-predicted Shh responses with Notch lateral inhibition (cyan) and without lateral inhibition (grey). Curves show mean with 95% CI. **i,** Schematic of uniform-Notch activation experiment. Light-activated ShineGal4 drives broad NICD expression in *Tg(UAS:nicd)* embryos during imaging. **j,** Positional information in wild type and uniform-Notch embryos. Insets, Shh response versus DV position at 12 ss; wild type subsampled to match cell numbers. Curves show mean with 95% CI. **k,** Positional information about final DV position, *PI*_12_(*g*_*t*_), for wild type and uniform-Notch embryos. Dashed lines, *PP*_12_(*d*_0_). Curves show mean with 95% CI.

Because Notch signalling mediates lateral inhibition^24,25^, maintains neural progenitor states^26,27^, and can modulate Shh sensitivity^28,29^, we asked whether the observed heterogeneity in Shh response reflected differences in Notch activity. We performed multiplex RNA measurements, using *ptch2* mRNA as a proxy for instantaneous Shh response and *her12* mRNA as a proxy for Notch activity^30,31^. Cells with low Notch activity were enriched among those with lower Shh response compared to the expectation based solely on their DV position (Fig. 4c,d). Consistently, inferred Shh sensitivity—defined as the deviation of Shh response from the expected DV position trend—correlated positively with progenitor markers activated by Notch signalling (*her12*^31^, *her4.1*^32^, *pcna*^33^, *sox19a*^34^) and negatively with most neurogenic markers repressed by Notch signalling (*dlb*^27^, *dld*^27^, *elavl3*^35^, *neurod4*^35^, except *dla*^27^) (Fig. 4e; Extended Data Fig. 8a–d; Supplementary Note 9). Together, cell-to-cell heterogeneity in Notch activity closely matches the heterogeneity in Shh sensitivity.

To test whether Notch-mediated lateral inhibition is sufficient to explain the lack of late positional-information accumulation, we added a lateral inhibition module upstream of the Shh response module in the hybrid model (Supplementary Note 8)^36,37^. Using the measured 3D trajectories, we updated each cell’s nearest neighbours at each time point, simulated Notch lateral inhibition between dynamically exchanging neighbours, and used the resulting Notch activity to modulate each cell’s Shh sensitivity (Fig. 4g). This extension generated diverse, effectively bimodal Notch activity that acted as an extrinsic source of variability and induced heterogeneous Shh sensitivity across cells. The resulting model captured the early positional information peak while eliminating the late accumulation, bringing simulations into close agreement with the measurements (Fig. 4h).

Because extending the hybrid model with a lateral inhibition module prevented positional information from increasing at late stages, we hypothesized that reducing heterogeneity in Notch activity would restore positional information accumulation. To test this prediction experimentally, we optogenetically enforced a uniformly high Notch activity by over-expressing Notch intracellular domain (NICD) throughout the imaging window (Fig. 4i; Extended Data Fig. 8e–g)^38,39^. Remarkably, uniform Notch activation significantly increased positional information at late stages (by ∼0.5 bits) over wild type (Fig. 4j), accompanied by a reduction of low-*ptch2* cells in lateral regions (Extended Data Fig. 8h,i). Importantly, positional information about final DV position, *PI*_12_(*g*_*t*_), continued to increase at late stages and ultimately exceeded the ceiling imposed by cell rearrangements, *PP*_12_(*d*_0_) (Fig. 4k). This later increase required that changes in Shh response remained informative about DV position once *PP*_12_(*d*_*t*_) rose above its earlier ∼1.2-bit ceiling from 0–6 ss (Fig. 2b,d). Taken together, these results demonstrate that Notch-driven heterogeneity prevents late accumulation of positional information.

### Early readout window conveys sufficient information for final ventral fates

Shh-encoded positional information peaks in the early readout window, but are cells utilizing this early window for their final fate decisions? To link positional information to fate, we mapped each lineage’s final cell fate (i.e., p3, pMN, p2 or p1) back onto its trajectories by multiplex *in situ* staining with fate markers immediately after live imaging (Fig. 5a–e; Extended Data Fig. 2e–h; Supplementary Method 3; Supplementary Video 3).

**Fig. 5.**
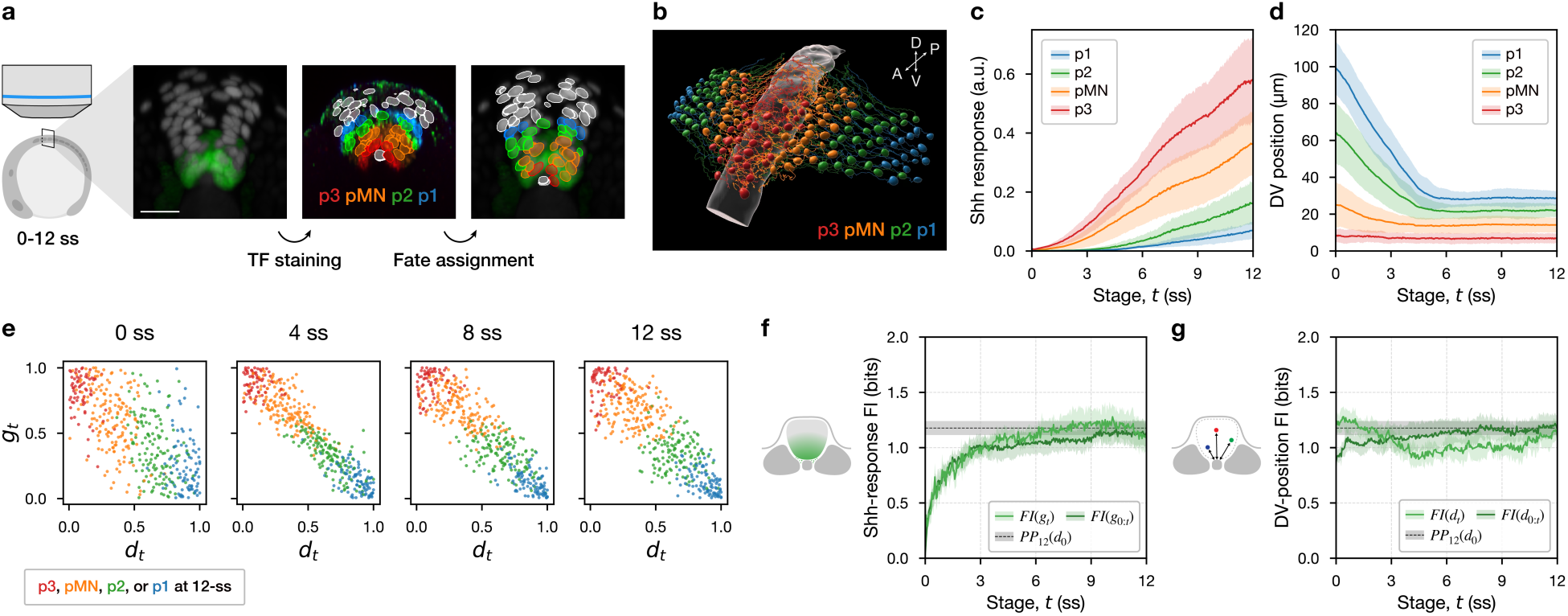
Fate mapping and timed Shh inhibition link the early readout window to ventral fate positioning. **a,** Workflow linking signalling histories to final fates. Embryos were imaged live from 0–12 ss, then fixed and stained for transcription-factor markers to assign each tracked nucleus to p3, pMN, p2 or p1 and map fate back onto its lineage trajectory. Scale bar, 20 µm. **b,** Example lineage reconstruction from one embryo, with initial positions coloured by fate at 12 ss. **c,d,** Shh response trajectories (**c**) and DV position trajectories (**d**) for lineage-traced cells grouped by 12-ss fate (mean ± s.d. across lineages; N=3 embryos). **e,** Relationship between within-embryo percentile-ranked Shh response *g*_*t*_ and DV position *d*_*t*_ at 0, 4, 8 and 12 ss, coloured by 12-ss fate. **f,** Fate information carried by Shh response at stage *t*, *FI*(*g*_*t*_) (light green), and by Shh response histories, *FI*(*g*_0:*t*_) (dark green) (Supplementary Notes 1 and 5). Dashed line, *PP*_12_(*d*_0_). Curves show mean with 95% CI. **g,** Fate information carried by DV position at stage *t*, *FI*(*d*_*t*_) (light green), and by DV position histories, *FI*(*d*_0:*t*_) (dark green). Display conventions as in **f**.

We defined fate information as the mutual information between final fate at 12 ss, *F*_12_, and either Shh response or DV position at stage *t* (Supplementary Note 1). Fate information carried by Shh response, *FI*(*g*_*t*_), rose rapidly and approached a plateau by ∼5 ss (Fig. 5f). Fate information did not improve when decoding final fate from the entire Shh response history relative to the snapshot Shh response (Fig. 5f; Extended Data Fig. 5k,m; Supplementary Note 5), indicating limited additional fate information in the Shh response history beyond snapshot readouts.

Notably, fate information closely matched positional information about final DV position (Fig. 5f compared with Fig. 2f,g), indicating that decoding final fate from Shh response is quantitatively equivalent to decoding final DV position from Shh response.

In contrast, fate information carried by DV position, *FI*(*d*_*t*_), was highest at the start of the imaging window and declined during convergence (Fig. 5g), consistent with the cross-temporal observation that Shh response was more predictive of earlier DV position than of instantaneous DV position during convergence (Fig. 2d,f). *FI*(*d*_*t*_) showed a modest late recovery after ∼10 ss (Fig. 5g), consistent with the observation that changes in DV position became weakly beneficial for positional information during this interval (Fig. 2g). Decoding final fate from full DV position trajectories likewise provided little improvement beyond the fate information carried by initial DV position, *FI*(*d*_0_) (Fig. 5g; Extended Data Fig. 5l,o; Supplementary Note 5).

Finally, decoding final fate from Shh response trajectories, DV position trajectories, or both jointly, converged to a similar value of ∼1.2 bits (Fig. 5f,g; Extended Data Fig. 5k–m). This remarkable agreement among all the information metrics is consistent with the aforementioned positional persistence ceiling, which limits how much information about final DV position can be preserved through neurulation—and therefore how much fate information can ultimately be extracted from Shh response as a readout of DV position. Thus, the early readout window is not only when Shh response is most informative about DV position but also when cells acquire most of the information relevant for later fate.

### Cells specify fate during the early readout window

Having established that cells accumulate sufficient information for ventral fates during the early readout window (0–5 ss), we asked whether this relationship is merely correlative or functionally required. We inhibited Shh signalling in three separate windows—before the early readout window (<0 ss), during the early readout window (0–5 ss), or after the early readout window (5–10 ss)—and assayed ventral fate patterning (p3, pMN, p2, p1, and p0) at 10 ss by multiplex *in situ* staining of marker genes (Fig. 6a–c; Supplementary Note 11). We assessed patterning phenotypes by quantifying fate proportions, marker intensity, mean DV position, and precision of each fate domain.

**Fig. 6.**
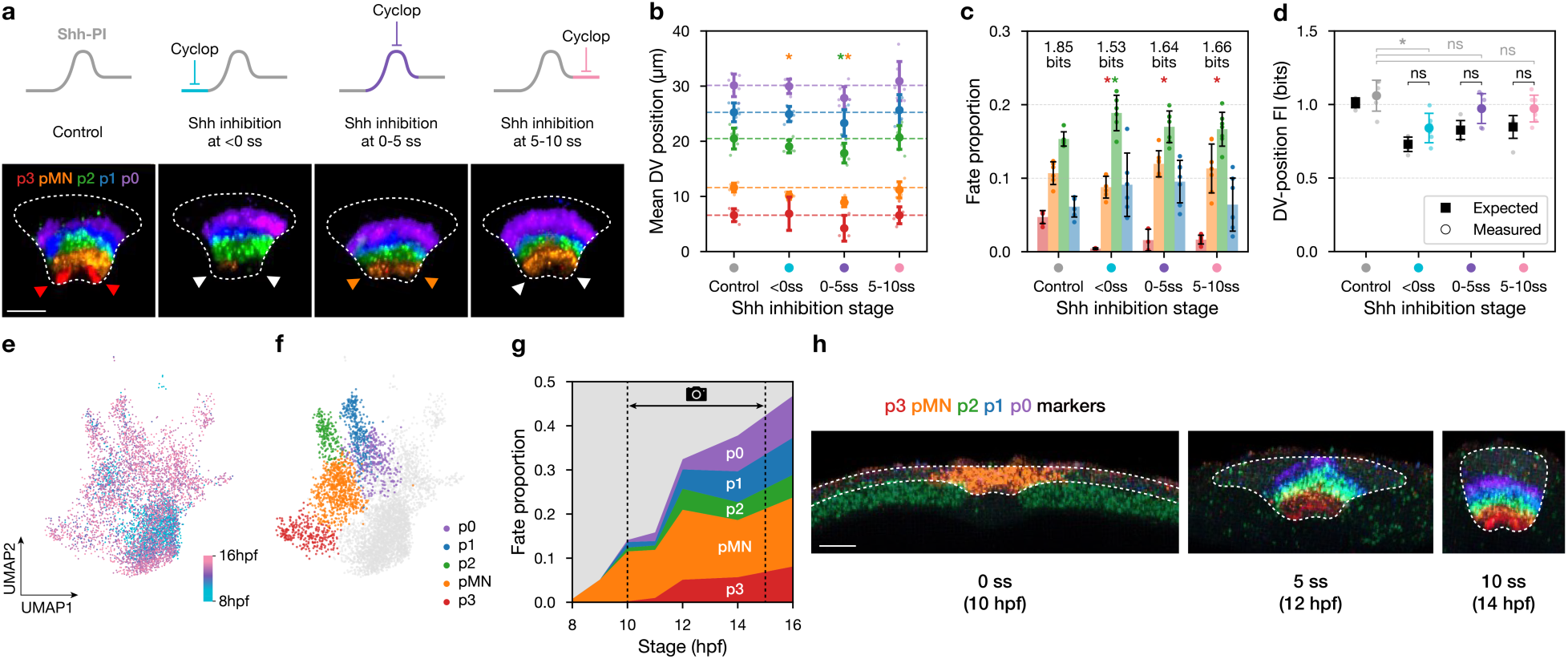
Early readout window aligns with emergence of ventral transcriptional states and peaks in fate regulator levels. **a,** Timed Shh inhibition with cyclopamine. Top, inhibition windows relative to the early readout window: before (<0 ss), during (0–5 ss), or after (5–10 ss). Bottom, representative 10-ss transverse sections stained for ventral fate markers. Arrowheads mark the inferred p3 position; colours denote the fate observed in that region (red, p3; orange, pMN; white, cells lacking p3–p1 markers). Scale bar, 20 µm. **b,** Mean DV position of each ventral fate domain. Points, embryos; large symbols, mean ± s.d.; dashed lines, control means. Asterisks, significantly different from control for the indicated fate, tested by one-way OLS with Holm correction (Supplementary Note 11). **c,** Proportions of ventral progenitor fates (mean ± s.d.); numbers, entropy within the p3–p1 population (bits). Points, embryos; bars, mean ± s.d. Statistics as in **b**. **d,** Fate information carried by DV position, *FI*(*d*_10_). Points, embryos; large symbols, mean ± s.d. Black squares indicate a composition-matched control baseline computed by resampling control fate labels to match each embryo’s observed p3–p1 fractions (Supplementary Note 11). Statistics as in b. **e,** UMAP of the 8–16 hpf spinal cord subset from the ZMAP zebrafish single-cell RNA-seq atlas, coloured by developmental stage. **f,** Ventral progenitor assignments (p3, pMN, p2, p1, p0) in the same dataset based on marker expression; light grey, dorsal progenitors and unassigned cells. Floor plate excluded. **g,** Proportions of ventral progenitor classes from 8–16 hpf. The 10–15 hpf interval (approximately 0–12 ss) brackets the live-imaging window used in this study. **h,** Representative trunk spinal cord sections stained by multiplex HCR RNA-FISH for ventral fate markers at 10 hpf (0 ss), 12 hpf (5 ss), and 14 hpf (10 ss). Dashed outlines, spinal cord. Scale bar, 20 µm.

Across all inhibition windows, p3 was strongly affected, with reduced p3 proportion and weaker p3 marker expression (Fig. 6c; Extended Data Fig. 9c–e). pMN proportion and marker expression were reduced when Shh signalling was inhibited before or after the early readout window, but not when Shh signalling was inhibited during the early readout window (Fig. 6c; Extended Data Fig. 9c–e). Strikingly, only inhibition during the early readout window produced a coherent ventral shift of the fate map, with each domain displaced ventrally by approximately one step: pMN expanded into the p3 territory, p2 into the pMN territory, p1 into the p2 territory, and p0 into the p1 territory (Fig. 6a,b). By contrast, inhibition before or after the early readout window produced no clear shift in mean DV position across fate domains, except for a mild expansion of pMN into the p3 territory when Shh signalling was inhibited before the early readout window (Fig. 6a,b). Thus, Shh signalling influences ventral fate proportions across windows, but ventral fate positions are specifically sensitive to Shh inhibition during the early readout window, when Shh response accumulates most fate information.

We next asked whether timed Shh inhibition reduced the fate information carried by DV position at 10 ss, *FI*(*d*_10_) (Supplementary Note 1). Relative to control, fate information showed a small decrease under Shh inhibition, but this reduction was not significant except when inhibition began before the early readout window (Fig. 6c). Mathematically, fate information depends on two factors: fate proportions (as quantified by fate entropy; Fig. 5d, bar labels) and the precision of fate boundaries. To separate these contributions, we constructed a proportion-matched control baseline by resampling control fate labels to match the p3–p1 fate proportions measured in each inhibition window (Fig. 5c, dark error bars; Supplementary Note 11). Compared with this proportion-matched baseline, fate information in the inhibition conditions showed no significant differences (Fig. 5d). We further verified independently that fate-boundary precision was unaffected by inhibition (Supplementary Note 11; Extended Data Fig. 9i,j). Together, timed Shh inhibition reduced fate information by changing fate proportions, but not by reducing the precision of the remaining fate domains.

Finally, we asked whether endogenous fate programs emerge during the same early readout window. To temporally dissect the emergence of ventral fate programs on a transcriptomic level, we analysed ZMAP, an integrated wild-type scRNA-seq resource spanning zebrafish development^40^, focusing on stages from 8–16 hpf, which encompass our imaging interval (10–15 hpf). Within the spinal cord subset, we performed unbiased clustering and assigned clusters to p3, pMN, p2, p1, and p0 based on established marker expression (Fig. 6e,f; Extended Data Fig. 10a; Supplementary Note 12). The proportion of cells assigned to these ventral fates increased sharply around ∼10–12 hpf, corresponding approximately to ∼0–6 ss (Fig. 6g). Multiplex *in situ* staining of fate markers showed that transcripts of ventral fate markers become detectable over the same interval. We first detected p3–p0 fate markers at 2 ss, and all embryos robustly expressed these fate markers by 5 ss (Fig. 6h; Extended Data Fig. 10b–d; Supplementary Note 13). Together with fate mapping and timed Shh inhibition, these transcriptional dynamics indicate that cells specify fate during the early readout window, when Shh response carries the most information, not when it is strongest.

## Discussion

For an organism to develop and function, cells must continuously interpret signaling cues to make decisions, yet the temporal dynamics of signal strength and signal quality need not coincide. Here, in a developmentally relevant *in vivo* context, we found that the window of peak Shh signaling strength is distinct from the window when signaling carries maximal positional information. Strikingly, cells align their fate decisions with this optimal information window — suggesting that developmental timing programs are not tuned for signal amplitude but rather for information content, a mathematically rigorous way of defining signal quality. This distinction has important implications for how we think about morphogen interpretation during tissue morphogenesis and cellular decision-making in general.

Our information-theoretic framework enables a direct comparison between two processes that are often analyzed separately: positional information gained through morphogen signal accumulation and positional information lost through cell rearrangements. By placing both on a common quantitative scale, in units of bits, we directly quantify the tug-of-war between how much information cells acquire through signaling and how much information survives physical reorganization of the tissue. Cell rearrangements preserved ∼1.2 bits between early and final DV positions, defining a morphogenesis-imposed ceiling for information transfer. Information metrics connecting Shh response to final position, DV position to final fate, and Shh response to final fate all converged near this same value, even when position was not directly used and when complete position or signaling histories were considered. Thus, morphogenesis sets a common limit through which position, signaling, and fate information can approach but cannot exceed.

Much of the classic literature assessed morphogen gradient precision from spatially averaged, mean expression profiles across embryos, leading to the conclusion that morphogen gradients are highly reproducible and precise^41–43^. We find that this reproducibility reflects the fact that variability between embryos is indeed minimal. Nevertheless, the dominant source of imprecision lies elsewhere: in the cell-to-cell variability within a single embryo among similarly positioned cells. Because mean profiles average over this within-tissue heterogeneity, they systematically underestimate the uncertainty each cell actually experiences. Quantifying positional information at true single-cell resolution is therefore essential for understanding what information is actually available to individual cells as they commit to a fate.

Obtaining the full signalling and positional histories of every cell in the ventral spinal cord also reveals information dynamics that are not accessible from snapshots. First, it enables comparison between information available from the snapshot morphogen response and the full signalling history. In the zebrafish spinal cord, decoding DV position from Shh response histories does not substantially exceed what is already present in the cumulative reporter readout at each stage, supporting the practical use of cumulative reporters as informative proxies in similar settings^41,43,44^. More broadly, lineages make cross-temporal information measurable: one can quantify the mutual information between a morphogen readout at one stage and cell position at another stage, or between cell positions at different stages. These cross-temporal analyses reveal information quantities invisible in snapshot analyses. For example, information about instantaneous position peaks early at ∼4 ss. However, whether this represents an optimal window for fate decisions depends on how much subsequent cell rearrangements shuffle positional order. Our cross-temporal analysis shows that morphogen response at ∼4 ss already carries near-maximal information about the final DV position, with little additional gain at later stages, consistent with an early window being the most informative for downstream fate decisions.

Positional persistence is especially valuable in this respect: it can only be quantified when cells are fully tracked over time, and it provides a direct, quantitative measure of how strongly tissue morphogenesis dynamics constrain information accumulation.

A natural strategy for cells that must read positional signals during morphogenesis would be to delay fate decisions until tissue movements have ceased, thereby minimizing the detrimental effect of cell rearrangements on positional information. Yet we find that cells in the zebrafish spinal cord do not adopt this strategy. Instead, they exploit an early window when positional information is at its peak, committing to fates before morphogenesis is complete. Crucially, our analysis shows that waiting until the end of morphogenesis yields no informational advantage: late-stage positional information does not exceed what was already available during the early readout window. If cell rearrangements proceed after fate specification, downstream mechanisms such as cell sorting or selective cell death can still refine patterns^45–47^, correcting errors without requiring a second increase in morphogen-encoded positional information. Thus, acting early is not a compromise — it is an informationally equivalent, and developmentally faster, solution.

A striking observation is that fate specification aligns in time with when signalling is most informative. This raises a broader question: is such temporal alignment simply the coincidence of two independently tuned processes, or are there mechanisms that couple the pace of fate programs to the information available in signalling? A related timing structure has been described in *Drosophila*, where positional information in the gap gene system peaks within a narrow window even without cell rearrangements, reflecting time dependence in the expression profiles themselves^5,6^. Defining what sets the gene expression dynamics of major fate regulators, and how these dynamics respond to changes in morphogen signalling dynamics, should help explain how fate programs are coordinated with morphogen information content.

This study has several limitations. First, continuous lineage tracking was restricted to the ventral spinal cord; extending these measurements across the full dorsoventral axis will require larger-volume imaging, likely at the cost of longer imaging intervals or lower spatial resolution.

Second, Shh response was measured with a single cumulative transcriptional reporter, *TgBAC(ptch2:Kaede)*, so the absolute magnitude and timing of information may vary with different signalling readouts. However, several lines of evidence support the interpretation that the early optimum reflects system-level information dynamics rather than reporter-specific kinetics. First, Shh-encoded positional information approached the theoretical ceiling defined by positional persistence, indicating that information transfer was nearly saturated. Second, the same rise-and-fall structure was recapitulated by a hybrid model in which Shh sensing was idealized and measured cell movement was the only empirical input. Finally, timed Shh perturbation showed that this early window is functionally required for correct fate positioning, and transcriptomic analyses showed that ventral progenitor states emerge during the same period. Together, these results support the conclusion that the early information optimum reflects the interaction between morphogen signalling and morphogenesis, rather than a peculiarity of the reporter.

Our study provides a first quantitative link between morphogen information to morphogenesis. Cell rearrangements do not merely add noise to an otherwise static readout; they actively define the temporal structure and set the ceiling of information transfer. Cell rearrangements during tissue morphogenesis are pervasive across embryogenesis and are often more extensive than in the neural plate^48^. We expect that applying lineage-resolved, cross-temporal positional information analyses across axes, morphogens, tissues and organisms will help define general strategies by which cells integrate morphogen signalling to specify fates in the right place and at the right time.

## Methods

### Zebrafish husbandry, staging and ethics

All zebrafish were raised, maintained and used for experiments in accordance with protocols approved by the Institutional Animal Care and Use Committees of Washington University in St. Louis. Embryos were obtained from natural matings, incubated at 28.5 °C and staged using standard morphological criteria and somite counts^49^.

### Zebrafish lines

Transgenic lines used in this study included: *TgBAC(ptch2:Kaede)*^21^, *Tg(8xgli-Xla.Cryaa:NLS-mCherry)*^50^ (this study, using the original construct), *Tg(sox19a:H2B-mCherry)*^51^, *Tg(sox19a:H2B-mNeonGreen)* (this study), *Tg(sox19a:membrane-mNeonGreen)* (this study), *Tg(−2.4shha-ABC:membrane-mCherry)*^52^, *Tg(−2.4shha-ABC:GFP)*^53^, and *Tg(UAS:myc-Notch1a-intra)*^39^.

Mutant lines included *cdh2^tm1^*^01^ ^54^ and *itga5^+20^* (this study). Maternal–zygotic itga5 mutants (MZitga5*^+20/+20^*) were produced by incrossing rescued homozygous adults. For cdh2*^tm101^*, heterozygous carriers were incrossed and embryos were imaged prior to genotyping; homozygous mutants were expected at Mendelian frequency.

### Transgenesis

To drive fluorescent reporters in neural progenitors, a 6,929-bp upstream regulatory fragment of *sox19a*^55^ was cloned into a Tol2 backbone^56^. Reporter cassettes encoded either H2B- or membrane-localization sequences followed by a GA linker (GAPAGGAGAA) fused to mNeonGreen or mCherry. Plasmids were assembled by Gibson assembly^45^ and co-injected with Tol2 transposase mRNA^56^ at the one-cell stage. Founders (F0) were outcrossed and stable lines were established by screening F1 offspring for the expected expression patterns.

### CRISPR mutagenesis and genotyping

For *itga5* editing, crRNA (AGGATACTTCTTGACCTCAG; IDT) was annealed with tracrRNA and complexed with Cas9 protein immediately before microinjection into one-cell embryos^57^. F0 outcrosses were genotyped by PCR (forward: CTGCGTTCAAACAAGAAATTG; reverse: AGGTGTCATCAAATCTGATCTCTTT), and a line carrying a 20-bp insertion was selected and propagated. To obtain viable *itga5* homozygous adults, embryos were rescued by one-cell injection of *itga5* mRNA as described previously^58^.

### Confocal imaging

Embryos were mounted individually in agarose moulds in egg water and imaged on an upright ZEISS LSM 980 confocal microscope^59^. For live time-lapse imaging, embryos were staged by completion of epiboly and imaged from 0 to 12 somite stages (ss) at 2-min intervals, beginning at bud stage (∼10 hpf). Volumes were acquired at 1 µm × 1 µm × 1 µm isotropic voxel size over a 303 µm × 303 µm × 145 µm field, encompassing trunk somites 2–6 (additional acquisition details in Supplementary Method 2). For fixed or live snapshot (non–time-lapse) imaging, embryos were mounted in PBS (fixed) or egg water (live) and imaged at 1 µm isotropic resolution. For time-course fixed embryos collected by hpf at 28.5 °C, we empirically mapped 10 hpf to 0 ss and 15 hpf to 12 ss and linearly interpolated intermediate stages for plotting.

### HCR RNA-FISH

HCR RNA-FISH was performed following the manufacturer’s protocol (Molecular Instruments). Unless otherwise noted, solution exchanges and washes (including probe and hairpin steps) were performed on an InsituPro robot (CEM Corporation). Embryos were fixed in 4% paraformaldehyde (PFA) at 28.5 °C for exactly 4 h, washed, pre-hybridized at 37 °C and hybridized overnight at 37 °C with 1 nM probe sets. After post-hybridization washes, embryos were incubated overnight in hairpin amplification mixtures at room temperature, washed and mounted for imaging. Where feasible, quantitative intensity analyses were acquired in the infrared channel (647) to reduce depth-dependent attenuation and background.

### Immunofluorescence

For Nkx6.1 staining, embryos were fixed in 4% PFA at 28.5 °C for exactly 4 h and washed in PBST (PBS + 0.1% Tween-20). Samples were blocked for 1 h in blocking buffer (10% foetal bovine serum, 1% DMSO, 0.1% Tween-20, 0.1% Triton X-100 in PBS), then incubated overnight at 4 °C with mouse anti-Nkx6.1 (DSHB F55A10, supernatant) at 1:20 in blocking buffer. PBST washes were performed on an InsituPro robot (CEM Corporation). Secondary staining was performed with Alexa Fluor 647 goat anti-mouse IgG (Invitrogen A32733TR) at 1:500 in blocking buffer. Embryos were washed and mounted for confocal imaging.

### Photoconversion of *TgBAC(ptch2:Kaede)* and spatial profiling of early versus late reporter accumulation

Embryos at 8 ss were exposed to 405 nm light for 1 min under a stereomicroscope (OLYMPUS MVX10) to photoconvert Kaede from green to red (pseudocoloured cyan and magenta in plots). After photoconversion, embryos were kept in the dark until 12 ss and then imaged by confocal microscopy.

For analysis, the pre-8-ss (photoconverted) and post-8-ss (newly-synthesized) nuclear signals were quantified per nucleus and rank-normalized independently to percentiles in [0,1] within each embryo. The per-cell difference was defined as (post-8-ss percentile – pre-8-ss percentile). To generate spatial heatmaps, DV position was represented as the percentile of nucleus-to-notochord distance, and mediolateral (left–right) bias was represented as (left − right)/(left + right). For each embryo, voxel intensities were averaged over shared DV/left–right coordinates and then averaged across embryos to obtain population heatmaps (Detail in Supplementary Note 10).

### Optogenetic Notch activation

To uniformly activate Notch, 50 pg ShineGal4 mRNA^38^ (zebrafish-codon optimized) was injected at the one-cell stage into *Tg(UAS:myc-Notch1a-intra)* embryos^39^. Embryos were kept in the dark until bud stage. During time-lapse imaging from 0–12 ss, the 488 nm excitation used for Kaede imaging simultaneously activated ShineGal4, driving widespread NICD (Notch intracellular domain) expression during the imaging window. mRNAs were synthesized from linearized templates using mMESSAGE mMACHINE SP6 kits (Invitrogen) and purified following the manufacturer’s instructions.

### Pharmacological perturbation of Shh signalling

For Shh inhibition, embryos were dechorionated at dome stage and maintained in 1× Danieau solution with matched solvent exposure. Cyclopamine (100 µM; LC Laboratories C-8700) was added at 28.5 °C starting in one of three defined windows: 5–10 hpf (before 0 ss), 10–12 hpf (during 0–5 ss), or beginning at 12 hpf (after 5 ss). Cyclopamine was washed out by three rinses in fresh 1× Danieau at the end of the specified window and embryos were then maintained in vehicle until fixation. Vehicle controls were treated identically with matched solvent exposure and rinse timing. Embryos were fixed at 10 ss for HCR RNA-FISH.

### Nuclear and tissue segmentation

Neural progenitor nuclei were segmented with StarDist-3D^60^ (StarDist v0.8.5) trained using the ZeroCostDL4Mic workflow^61^ on independently acquired high-resolution stacks; representative performance is shown in Extended Data Fig. 1a, and full training details are provided in Supplementary Method 1.

Spinal cord, notochord and floor plate masks were generated using 3D U-Net–style convolutional networks^62^ trained on fixed embryos with transgenic and/or HCR-based ground-truth masks; training details and validation are provided in Supplementary Method 1 with examples in Extended Data Fig. 1b.

For downstream quantification, segmentation outputs were imported into Imaris as Surfaces, including a notochord surface predicted by the 3D U-Net.

### Lineage reconstruction and trajectory curation

After time-lapse acquisition, nuclei were segmented and imported into Imaris for tracking. Initial tracks were generated using Brownian motion tracking with a maximum step of 5 µm to link nuclei between frames, and were then manually extended and curated to follow all cells present at the initial time point across the full movie unless they exited the imaging volume. Floor plate cells were excluded based on medial position and low *TgBAC(ptch2:Kaede)* signal at the final time point. Tracking workflow, quality control and examples are shown in Extended Data Fig. 2, with detailed procedures in Supplementary Method 2.

In transplantation-based ground-truth tests, the automatic tracking step achieved 99.6% correct auto-linked connections, and automatic tracking followed by manual correction achieved 100% accuracy for reconstructed lineages (see Supplementary Method 2 for details).

### Quantification of Shh response and DV position

For time-lapse analyses, Shh response was quantified per nucleus as the mean nuclear Kaede intensity exported from Imaris. DV position was quantified as the shortest distance from each nucleus surface to the notochord surface in Imaris (spot-to-surface / surface-distance mode; unsigned shortest distance). The notochord surface was generated from the 3D U-Net notochord mask imported into Imaris. For benchmarking, centroid-to-surface distance was also computed in Python, but all primary analyses use the Imaris shortest surface-to-surface distance.

For fixed-embryo assays (HCR and immunofluorescence), DV position was computed using the same notochord-surface reference and shortest-distance definition to enable direct comparison between live and fixed datasets.

### Fluorescence background correction and intensity normalization

To reduce systematic imaging artifacts, fluorescence channels were background-corrected before downstream normalization and information-theoretic analyses. For *TgBAC(ptch2:Kaede)* and *ptch2* mRNA, background was estimated within each embryo and snapshot by fitting the reporter intensity as a function of dorsoventral (DV) position and decomposing this DV-dependent profile into an exponential-decay component plus a constant offset; the fitted background component was subtracted from per-cell values. For HCR channels used for transcription-factor quantification, the DV-dependent intensity profile was instead modelled using a Gaussian peak plus constant offset; the fitted background term was subtracted. These fits were performed per embryo and per snapshot (timepoint for time-lapse data) to avoid conflating biological dynamics with imaging drift.

For analyses that pool cells across embryos, within-embryo rank normalization was performed independently at each somite stage. Specifically, for each embryo and stage, Shh response and DV position values across cells were converted to percentiles in [0,1] and then pooled across embryos for stage-specific analyses.

### Live-to-fixed fate mapping

Immediately after time-lapse imaging, embryos were fixed overnight in 4% PFA at 4 °C and processed for HCR RNA-FISH. Ventral progenitor domains were assigned using marker sets (p3: *nkx2.2a*/*nkx2.2b*/*nkx2.9*; pMN: *olig2*; p2: *lbx2*; p1: *prdm12b*) and identities were called manually from multiplex signal patterns. Live-imaged lineages were mapped to fixed identities by manual registration of nuclei at the final live time point to nuclei in the fixed specimen, guided by skin nuclei, floor-plate position, and the overall neural progenitor population. This workflow and validation are described in Supplementary Methods.

### Quantification of ventral domain positions and boundary precision in fixed embryos

Embryos were fixed and stained by multiplex HCR RNA-FISH for p3–p0 markers together with DAPI. Spinal cord and notochord volumes were segmented using a 3D U-Net, imported into Imaris as surfaces, and nuclei were detected from DAPI and restricted to nuclei within the spinal cord mask. Cell types were assigned in Imaris using the built-in supervised classification workflow trained on HCR intensity features from manually labelled nuclei and then applied uniformly across all embryos in the dataset using the same trained model. DV position was computed as the shortest 3D distance from each nucleus (spot) to the notochord surface.

Boundary precision was assessed by (i) fate information between DV position and fate labels and (ii) within-domain dispersion of DV position (standard deviation), as described in Supplementary Note 11.

### ZMAP scRNA-seq resource analysis

To place ventral gene programs on a transcriptome-level timeline, we analysed ZMAP, focusing on spinal cord cells from 8–16 hpf. After filtering to this subset, the analysis encompassed 66 libraries across 6 independent studies. Time was represented as hours post-fertilization (hpf) as annotated in the original datasets. Cells were reanalysed using ZMAP’s existing annotations as a starting point, followed by reclustering of the spinal cord subset; ventral progenitor identities (p3, pMN, p2, p1, p0) were assigned based on canonical marker programs. Floor plate cells were excluded before computing ventral-class fractions, which were calculated as fractions of the spinal cord population across time. Full details are provided in Supplementary Note 11.

### Mutual information estimation

For each embryo and each somite stage, Shh response and DV position values were rank-normalized to [0,1], and cells were then pooled across embryos within each condition and stage. Continuous variables were discretized by quantile binning into equiprobable bins, and mutual information was computed from the resulting joint contingency table. To control finite-sample upward bias in the plug-in estimator, we used a subsampling–extrapolation (“direct”) debiasing procedure^19,20^: for each of *n*_chain_Monte Carlo chains, plug-in mutual information was computed on nested subsamples and extrapolated linearly to the infinite-sample limit (1/*n* → 0). Reported mutual information is the mean of the extrapolated intercepts across chains.

Uncertainty is reported as mean ± 1.96×s.d. across chains, reflecting variability introduced by the subsampling–extrapolation procedure. In simulation benchmarks using synthetic datasets with known ground-truth mutual information, this interval achieved ∼95% coverage of the true value. Because chains are generated by repeated subsampling from the same underlying dataset, they are not independent replicates; standard error–based confidence intervals that further divide by 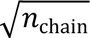 would therefore underestimate uncertainty and are inappropriate. Bin resolution was chosen conservatively using shuffle-null controls. Full details are provided in Supplementary Note 2–3.

### Trajectory-based decoding of DV position and fate from histories

All history-based decoding analyses follow the same strategy (detailed in Supplementary Note 5). At each somite stage *t*, we form a per-lineage history feature vector from measurements up to *t* (Shh response history, DV position history, or their concatenation, depending on the panel). We then train supervised decoders using outer 5-fold cross-validation across lineages, pool out-of-fold predictions across folds, and quantify decoding-based information by computing mutual information between predicted and measured targets using the same estimation conventions as in the corresponding snapshot analyses (Supplementary Note 2).

Decoder choice and feature-set choice were benchmarked separately for continuous versus categorical targets. For continuous DV position decoding, we selected ridge regression (Extended Data Fig. 5a–c). For categorical fate decoding (12-ss fate), we selected a Random Forest model (Extended Data Fig. 5d–f).

### Hybrid modelling of signalling readout from trajectories

We implemented a hybrid modelling strategy to isolate the contribution of morphogenesis—cell-specific DV position trajectories in absolute distance units (µm)—to cumulative Shh response dynamics, while holding the Shh ligand field and sensing/response rule fixed (Supplementary Note 8). Using wild-type lineage-traced embryos, we fitted each candidate model once by minimizing the root-mean-square error of log-transformed Shh response values across all lineages and timepoints, which balances early and late parts of the trajectory and prevents late high-intensity frames from dominating the fit. We benchmarked multiple sensing/response architectures (Extended Data Fig. 7) and assessed cross-condition transfer in the units most relevant for information analyses by additionally evaluating a within-embryo percentile loss.

Based on percentile-loss generalization under morphogenesis perturbation (prioritizing *MZitga5^-/-^*), we selected the sensing logic “Hill1_2layers” (Extended Data Fig. 7a; Supplementary Note 8). The resulting parameter set was then held fixed and applied unchanged to morphogenesis-perturbed mutants so that differences in predicted Shh response arise from substituting the experimentally measured DV position trajectories rather than refitting.

To simulate Notch-mediated lateral inhibition, we extended the hybrid model with an upstream lateral-inhibition module with tuned (not fit) parameters: nearest-neighbour relations were inferred from the measured 3D trajectories, Notch states were iteratively updated to generate a bimodal distribution, and the resulting Notch state modulated the Shh response cascade (Extended Data Fig. 7f,h,k; Supplementary Note 8). Positional-information quantities computed from model-predicted Shh responses used the same mutual-information pipeline as for experimental data (Supplementary Note 8).

### Statistics and reproducibility

Sample sizes are reported as embryos (N; biological replicates) and as the number of lineage trajectories or cells (n) contributing to pooled mutual-information analyses. For time-lapse lineage datasets, sample sizes were: wild type, N = 9 embryos (n = 1,561 lineages); *cdh2*^-/-^, N = 4 (n = 566); *MZitga5*^-/-^, N = 4 (n = 618); uniform Notch activation, N = 3 (n = 495); and fate-mapped wild type, N = 3 (n = 445). Unless stated otherwise, error bars represent standard deviation across embryos (for embryo-level summaries) or 95% CI for mutual-information estimates (Supplementary Note 2).

No embryos were excluded based on quantitative outcomes. Time-lapse datasets were excluded only if large embryo rotation prevented consistent anatomical registration, if <100 µm of the anteroposterior trunk region of interest was contained in the imaging window, or if a substantial portion of the spinal cord moved outside the lateral field of view during acquisition. Each time-lapse movie contains one embryo; thus, biological replicates derive from independent clutches.

Blinding was not used for time-lapse lineage analyses because tracking and linkage were performed using predefined computational pipelines without condition-dependent manual decisions. For staining-based datasets requiring supervised classification, embryos were pooled into a single multi-embryo image file (as a time dimension), the same trained classifier was applied across the pooled dataset, and condition identity was revealed only during downstream analysis by embryo/time identifier. Statistical tests and multiple-comparison corrections are reported in figure legends; biological replicate N was used as the unit of inference, and Holm–Bonferroni correction was used unless otherwise specified. Analyses were performed in Python.

## Code availability

Scripts for mutual-information estimation and hybrid forward modelling are available at: https://git.ista.ac.at/jrenaud/pi_while_morphogenesis_methods.

## Data availability

The processed data used in this manuscript, along with the corresponding plotting codes, are available at: https://git.ista.ac.at/jrenaud/pi_while_morphogenesis_methods.

## Acknowledgements

We thank members of the Tsai and Tkacik labs and Dr. Lilianna Solnica-Krezel for critical discussions and reading of the manuscript. We are grateful to A.J. Federico and David Bruckner for the earlier version of the analysis and stimulating discussion, the Peng Huang and Scott Holley labs for sharing transgenic fish, Yohanns Bellaiche and Pedro Hernandez Cerda for sharing optogenetic reagents, and the Solnica-Krezel lab for sharing laboratory equipment. This work was supported by US National Institutes of Health (NIH) Maximizing Investigator’s Research Award (MIRA) grant (R35GM150759) to C.-T.C. and T.Y.-C.T., Taiwan Ministry of Education fellowship to C.-T.C., and by the European Research Council (ERC-2023-SyG, ‘Dynatrans’, grant no. 101118866) to J.R. and G.T.

## Author contributions

C.-T.C. and T.Y.-C.T. conceived and designed the study. C.-T.C. developed and performed the zebrafish experiments and deep-learning tracking pipelines. J.R. and C.-T.C. developed the information-theoretic framework and associated analyses, and J.R. developed the hybrid modelling. C.-T.C. and J.R. performed the data analyses, and C.-T.C. rendered the plots. T.Y.-C.T. and G.T. supervised the research. C.-T.C. and T.Y.-C.T. wrote the manuscript with substantial input and critical revisions from J.R. and G.T. All authors approved the final manuscript.

## Extended Data Figure Legends

**Extended Data Fig. 1. Deep-learning segmentation of nuclei and reference tissues.**

**a,** Representative confocal transverse optical section from a 10-ss neural tube used for nucleus-segmentation benchmarking (independent test image; not used for model training or validation). From left to right: raw image; manually curated ground-truth nuclei; StarDist-3D nucleus instance segmentation; nuclei detected by Imaris (Spots). Colored overlays indicate individual nucleus instances.

**b,** Benchmark of nucleus-detection error modes for StarDist-3D versus Imaris Spots across Spot diameter settings, evaluated on an independent test set (not used for model training or validation). For each embryo, points show the fraction of ground-truth nuclei classified as split (one ground-truth nucleus matched to >1 prediction) or missing (no prediction matched to a ground-truth nucleus). Large symbols show mean ± s.d. across embryos. The y axis is plotted linearly from 0 to 0.1 and on a log scale above 0.1.

**c,** Representative 3D U-Net tissue segmentation across the live time-lapse window used for lineage analyses (example confocal transverse optical sections; not part of the segmentation training/validation datasets). Top row: confocal transverse optical sections at 0, 4, 8 and 12 ss from embryos carrying *Tg(sox19a:H2B-mCherry)* and *Tg(shha:mem-mCherry)* (magenta) together with the Shh-response reporter *TgBAC(ptch2:Kaede) (green)*. Bottom row: corresponding tissue masks (spinal cord, floor plate and notochord) overlaid on the same sections. Scale bar, 20 µm.

**d,** Representative 3D U-Net segmentation of reference tissues in fixed embryos stained with DAPI across stages (example confocal optical sections; not part of the segmentation training/validation datasets). Top row: DAPI optical sections at 0, 5, 7 and 12 ss. Bottom row: corresponding masks for spinal cord, notochord, somites and skin overlaid on the same sections. Scale bar, 20 µm.

**Extended Data Fig. 2. Quality control for lineage reconstruction and live-to-fixed fate mapping.**

**a,** Schematic of sparse cell transplantation used to generate “ground-truth” donor cells for quality control of lineage reconstruction. Donor cells were labelled with membrane-mNeonGreen and Dextran-AF647.

**b,** Representative confocal optical sections from a transplanted embryo at 0-, 6-, and 12-somite stages (0, 300, and 600 min) showing sparse donor-cell labelling that enables unambiguous manual tracking across frames. Scale bar, 20 µm.

**c,** Comparison of automatically reconstructed donor-cell trajectories with manually curated ground-truth trajectories. Tracks are shown only for donor cells. A representative misconnection is highlighted (cyan): an Imaris track that is incorrectly linked into the donor lineage set.

**d,** Accuracy of automatic frame-to-frame linking for donor-cell tracking using Brownian-motion linking (maximum step size 5 µm; no gap closing), comparing segmentations from Custom StarDist-3D versus Imaris Spots (benchmark dataset only; not used for model training/validation). Error categories are illustrated schematically (left). For each method, points show per-cell linking error (%) for the indicated error modes; counts indicate the number of affected cases. Manual curation produced 100% correct linking for the evaluated donor-cell set.

**e,** Workflow for live-to-fixed fate mapping. After live imaging, embryos were fixed and stained by multiplex HCR RNA-FISH to assign each tracked nucleus to a ventral progenitor fate domain (p3, pMN, p2, p1) in the fixed specimen, then mapped back to the terminal frame of each live lineage.

**f,** Schematic of sparse-labelling strategy used to quality-control live-to-fixed mapping: sparse donor-cell labels are imaged live, then re-identified after fixation and HCR processing to verify correspondence across modalities.

**g,** Representative confocal optical sections showing sparse labels in the live embryo, after HCR processing, and the resulting correspondence used for identity matching. Scale bar, 20 µm.

**h,** Live-to-fixed identity matching success rate across embryos. Bars show the number of successfully matched cells out of the total evaluated cells per embryo.

**Extended Data Fig. 3. Selection of Shh-response and dorsoventral (DV) position metrics and an upper bound on reporter delay.**

**a–c,** Analysis of embryos expressing the Shh-response reporter *TgBAC(ptch2:Kaede)* (N = 36 embryos; n = 26,557 cells).

**d–f,** Analysis of embryos expressing *Tg(8xgli-Xla.Cryaa:NLS-mCherry)* (GBS) (N = 6 embryos; n = 4,666 cells).

For the cell-level scatter plots in **b,c,e,f,** we randomly subsampled 4,000 cells per reporter to balance cell density for visual comparison.

**a,d,** Representative confocal optical transverse sections showing reporter signal in the trunk spinal cord at 10 ss (*ptch2:Kaede*, green, a; *GBS:mCherry-NLS*, red, d). Dashed outline, spinal cord. Scale bar, 20 µm.

**b,e,** Single-cell Shh response versus raw DV position (µm) for *TgBAC(ptch2:Kaede)* (b) and GBS (e). Points, cells. Reporter values are normalized to unit mean amplitude to enable comparison across reporters.

**c,f,** As in **b,e** after within-embryo rank normalization of Shh response and DV position to percentiles (variables *g*_*t*_ and *d*_*t*_). Points, cells. Left, *g*_*t*_ binned by *d*_*t*_ (bin width 0.1; mean ± s.d.). Right, *d*_*t*_ binned by *g*_*t*_ (mean ± s.d.).

**g,** Comparison of fitted Shh-gradient decay length (µm) between *TgBAC(ptch2:Kaede)* and GBS. Points, embryos; bars, mean ± s.d. across embryos. *P* value from two-sided *t*-test.

**h,** Positional information (bits) at ∼10 ss computed from each reporter (N = 36 embryos for *ptch2*:Kaede; N = 6 embryos for GBS). Points, embryos; bars, mean ± s.d. across embryos. *P* value from two-sided *t*-test.

**i,** Confocal optical transverse sections of reporter mRNA at 10 ss (*ptch2:Kaede* mRNA, green; *GBS:mCherry* mRNA, red). Dashed outline, spinal cord. Scale bar, 20 µm.

**j,k,** Time courses of mRNA gradient amplitude (j) and decay length (k) for *ptch2:Kaede* and *GBS:mCherry* transcripts (N = 5–12 embryos per transgene per stage). Points, embryos; curves show mean ± s.d. across embryos.

**l,** Sequential Shh inhibition to bound the effective *TgBAC(ptch2:Kaede)* readout delay. Cyclopamine was applied at 0, 3, 6, or 9 ss and maintained until 12 ss. The *ptch2:Kaede* gradient amplitude at 12 ss was normalized to the no-inhibition control and plotted versus inhibition duration (N = 8 embryos per condition; points, embryos; mean ± s.d. across embryos). Dashed line, linear fit to condition means. Green line, x-intercept (mean ± s.d. across bootstrap replicates), giving an upper bound on the effective delay that includes drug onset, pathway shutdown, and Kaede fluorescence dynamics.

**m,** Corresponding analysis in the hybrid model using the measured DV-position trajectories from the N = 8 wild-type embryos. Applying the same sequential-inhibition protocol *in silico* yields an approximately linear dependence of the Shh-response amplitude at 12 ss on inhibition duration (points, simulated embryos; mean ± s.d.), supporting interpretation of panel **l**’s intercept as a conservative upper bound on the intrinsic signalling-to-readout delay.

**n–r,** Mutual information computed using alternative DV-position metrics, plotted against the notochord-surface shortest-distance metric (N = 18 embryos across the 0–12 ss time course). Points, embryo time points; dashed line, identity. DV-position metrics were computed using the tissue-segmentation model shown in Extended Data Fig. 1d.

**Extended Data Fig. 4. Quality control of mutual-information estimation, binning resolution and sample size.**

**a,** Example of direct subsampling–extrapolation debiasing for a pooled wild-type estimate (illustrated for PI at 4 ss). Points show the naïve plug-in mutual information computed on nested subsamples of size *n*, plotted versus 1/*n*. Coloured lines show independent linear fits (“chains”) across repeated subsampling/fit realizations; the black point at 1/*n* = 0 indicates the extrapolated infinite-sample intercept, reported as mean with 95% CI.

**b,** Example of dependence on bin number *Q* for pooled positional information at 4 ss. Green line, debiased estimate (mean with 95% CI across debiasing chains). Grey line, shuffle-null control generated by permuting Shh response relative to DV position 30 times (mean with 95% CI across permutations, using the same estimator settings). Vertical dashed line marks the bin number used for pooled estimates (*Q* = 30 ∼*Q*_*max*_(*n*_*pooled*,*WT*_)).

**c,** Example of dependence on bin number *Q* for per-embryo positional information at 4 ss. For each *Q*, positional information was computed separately in each embryo and then summarized across embryos. Green line, mean with 95% CI across embryos; grey line, shuffle-null control (mean with 95% CI across embryos, using within-embryo permutations). Vertical dashed line marks the bin number used for per-embryo estimates (*Q* = 10 ∼*Q*_*max*_(*n*_*embryo*,*WT*_)).

**d,** Lookup curve *n*_*min*_(*Q*)∼*Q*_*max*_(*n*) across bin numbers *Q* (dashed), summarizing the minimum sample size required for shuffle-null bias to be negligible under the chosen estimator settings.

Coloured points show the (*Q*, *n*) pairs used across datasets/conditions (including wild type and perturbations), confirming that all analyses were performed above the corresponding *n*_*min*_(*Q*) threshold.

**e,** Effect of binning resolution on binned mutual-information estimates: comparison of estimates obtained using reduced bin numbers *Q* versus a high-resolution reference (*Q* = 30). Each point corresponds to a matched estimate (same stage/MI type), plotted against the *Q* = 30 value; dashed line, identity.

**f,** Summary of the binning-resolution benchmark in **e**, reporting agreement metrics (slope and *R*^2^) as a function of *Q*, evaluated relative to the *Q* = 30 reference.

**g,** k-nearest-neighbour (kNN) mutual-information estimates benchmarked against the *Q* = 30 binned reference. Points show matched estimates across stage/MI type; dashed line, identity.

**h,** Dependence of kNN estimates on neighbourhood size k, summarized as agreement metrics relative to the *Q* = 30 reference (as in **f**), to assess estimator stability across *k*.

**i,** Agreement between pooled and per-embryo binned estimates: per-embryo mutual information estimates (computed separately in each embryo) plotted against the corresponding pooled-data estimates (pooling embryos at each stage after within-embryo rank normalization) (both estimate use Q=10). Dashed line, identity.

**j,** Summary of per-embryo-versus-pooled agreement in **i** as a function of the number of embryos pooled (*N*) (*Q* = 10), reporting slope and *R*^2^.

**k,** Agreement between pooled and per-embryo binned estimates as in **i** but using kNN method (both estimates use *k* = 3).

**l,** Analogous per-embryo-versus-pooled comparison for kNN mutual-information estimates (*k* = 3), plotted as in **j**; dashed line, identity.

**m,** Comparison of positional-information dynamics computed by retaining one descendant per lineage after division (*n* = 1,561 lineages throughout) versus retaining both descendants after division (*n* increases from 1,561 to 2,099).

**Extended Data Fig. 5. Benchmarking decoder-based positional and fate information and testing robustness to Shh-response delay.**

**a,b,** Example single-lineage time traces used for decoder benchmarking. a, DV position *d*_*t*_trajectories for a subset of lineages. **b,** Corresponding Shh-response histories *g*_*t*_ for the same lineages.

**c–e,** Example decoder performance for predicting DV position at different target stages from Shh-response history. Scatter plots compare decoded versus measured DV position for c, *d*_0_, d, *d*_6_, and e, *d*_12_ using a ridge-regression decoder; dashed line indicates identity.

**f–h,** Benchmarking alternative decoders for extracting trajectory-based positional information from Shh-response history *g*_0:*t*_ about three DV-position targets (Supplementary Note 5). Curves show decoder-based positional information versus stage *t* for predicting f, instantaneous DV position *d*_*t*_, g, initial DV position *d*_0_, and h, final DV position *d*_12_}. Methods shown include ridge, lasso, elastic net, random forest, gradient-boosted trees, kNN, and SVR (legend).

**i,j,** Testing robustness to effective Shh-response delay. Positional information dynamics computed after shifting the Shh-response input by an assumed delay (0–3 ss): **i,** instantaneous positional information *PI*(*g*_*t*_); **j,** trajectory-based positional information from histories *PI*(*g*_0:*t*_). Allowing delayed readout does not remove the late decline in positional information.

**k–m,** Decoder-based trajectory-based fate information about fate at 12 ss, *F*_12_, using histories up to stage *t*. Curves show fate information extracted from k, Shh-response histories *g*_0:*t*_, **l,** DV-position histories *d*_0:*t*_, and m, joint histories (*g*_0:*t*_, *d*_0:*t*_), benchmarked across decoder classes as in **f–h** (legend).

**n,o,** Fate-decoding performance of the retained decoder (Random Forest) using the full history to 12 ss. Bubble plots compare decoded (y-axis) versus observed (x-axis) *F*_12_ using **n**, Shh-response history *g*_0:12_ or **o**, DV-position history *d*_0:12_. Bubble area indicates the fraction of cells.

**p–t,** Local “PI gain” decomposition on the cross-temporal PI surface (Supplementary Note 6). **p,** Example illustrating how local changes near an anchor stage (example shown at 3 ss) are decomposed into a Shh-response contribution and a DV-position contribution. **q,** Shh-response contribution *ΔPI*(*g*_*t*_) estimated from a local slope along the Shh-response axis at fixed target stage. **r,** DV-position contribution *ΔPI*(*d*_*t*_) estimated from a local slope along the DV-position axis at fixed Shh-response stage. **s,** Cumulative sums of *ΔPI*(*g*_*t*_) and *ΔPI*(*d*_*t*_) across stages, showing their opposing contributions to the net change in positional information. **t,** Reconstructing the positional information time course by adding the cumulative contributions in **s** to the initial positional information *PI*(*g*_0_), and comparing with the measured pooled positional information trajectory.

**Extended Data Fig. 6. Shh-response profile fits, positional information, and positional persistence under morphogenesis perturbations.**

**a,b,** Cross-temporal positional information from Shh response, summarized as cuts through *PI*_*t*’_(*g*_*t*_). **a,** information about the initial DV position, *PI*_0_(*g*_*t*_). **b,** information about the final DV position, *PI*_12_(*g*_*t*_). Curves show mean with 95% CI for each genotype.

**c,d,** Cross-temporal positional persistence, summarized as cuts through *PP*_*t*’_(*d*_*t*_). **c,** persistence about the initial DV position, *PP*_0_(*d*_*t*_). **d,** persistence about the final DV position, *PP*_12_(*d*_*t*_).

Curves show mean with 95% CI for each genotype.

**e,** Mean DV-position dynamics plotted versus morphogenesis-normalized stage, *t*° (Supplementary Note 8). Solid lines, trajectories versus morphogenesis-normalized stage; dashed lines, the same trajectories versus original somite stage. Curves show mean ± s.d. across embryos.

**f,g,** Positional persistence versus morphogenesis-normalized stage. **f,** 1-somite-stage interval persistence, *PP*_*t*+1_(*d*_*t*_). **g,** persistence about the initial DV position, *PP*_0_(*d*_*t*_). Curves show mean with 95% CI.

**h,** Positional information about instantaneous DV position plotted versus morphogenesis-normalized stage, *PI*(*g*_*t*_).

**Extended Data Fig. 7. Benchmarking and using the hybrid forward model.**

**a,** Model-selection summary across alternative sensing logics. For each sensing logic, performance is shown for the single best-performing parameterization (selected using wild type and then evaluated across conditions). Top, root-mean-square (RMS) error in log space between predicted and measured Shh response in absolute units. Bottom, RMS error after within-embryo rank normalization (rank-percentiles). Points, embryos; large symbols, mean ± s.d. across embryos. Models are ordered by the maximum (worst) mean RMS rank-percentile error across conditions.

**b–d,** Positional information computed from model-predicted Shh responses using alternative sensing logics without added noise. **b,** instantaneous positional information, *PI*(*g*_*t*_). **c,** positional information about initial DV position, *PI*_0_(*g*_*t*_). **d,** positional information about final DV position, *PI*_12_(*g*_*t*_). Curves show mean with 95% CI.

**e,f,** Positional information for the same model set after adding either **e**, stage-independent (“constant”) noise to the predicted Shh response or **f,** a neighbour-coupled lateral-inhibition module that generates cell-to-cell heterogeneity in Shh sensitivity. Display conventions as in **b–d**.

**g,h,** Positional information computed from model-predicted Shh responses when substituting experimentally measured DV trajectories from morphogenesis-perturbed embryos. **g,** no added noise. **h,** lateral-inhibition setting. Curves show mean with 95% CI.

**i–k,** One-dimensional slices through cross-temporal positional information, *PI*_*t*′_(*g*_*t*_), shown for **i,** no added noise, **j,** constant-noise setting, and **k,** lateral-inhibition setting. Curves show mean ± s.d. across embryos; mutual-information estimates are summarized as mean with 95% CI within each embryo.

**Extended Data Fig. 8. Notch-state correlates of Shh sensitivity and effects of uniform Notch activation.**

**a,** Confocal optical cross-section showing endogenous *ptch2* mRNA at 10 ss (left) and the corresponding DV–LR (dorsal-ventral–left-right) mean expression profile across the spinal cord (right). Dashed outline, spinal cord. Scale bar, 20 µm.

**b,** Definition of Shh sensitivity from single-cell *ptch2* expression at 10 ss, illustrated using the *her12* dataset (N = 6 embryos; n = 5,561 cells). Left, *ptch2* mRNA versus DV position for individual cells (points, cells; colours, embryos). Middle, rank-normalized *ptch2* level in [0,1]. Right, Shh sensitivity defined as the residual (deviation) from the expected DV trend (dashed line), with points coloured by sensitivity (higher = more Shh-sensitive; lower = less Shh-sensitive).

**c,** Relationship between Shh sensitivity and representative Notch-associated targets at 10 ss: *her12* (Notch-associated/progenitor; left; N = 6 embryos; n = 5,561 cells) and *elavl3* (neurogenic; right; N = 6 embryos; n = 5,052 cells). Points, cells; black line, mean Shh sensitivity within 0.1-wide expression bins; error bars, s.d. across cells within bins.

**d,** DV–LR expression profiles (top, confocal optical sections; bottom, corresponding DV–LR mean maps) for Notch-associated targets at 10 ss used in this study: *her12* (N = 6; n = 5,561), *her4.1* (N = 6; n = 5,584), *pcna* (N = 5; n = 4,557), *sox19a* (N = 5; n = 4,584) (cyan; progenitor/Notch-associated) and *dla* (N = 5; n = 4,470), *dlb* (N = 5; n = 4,651), *dld* (N = 5; n = 4,453), *elavl3* (N = 6; n = 5,052), *neurod4* (N = 5; n = 4,528) (magenta; neurogenic/Notch-negative). Dashed outlines, spinal cord. Scale bars, 20 µm.

**e,** DV–LR mean profiles of *her12* mRNA at 12 ss in control (dark) versus uniform Notch activation (light, 0–12 ss) embryos generated by ShineGal4-driven NICD expression (ShineGal4 mRNA injection in *Tg(UAS:nicd)* embryos). Dark, N = 10 embryos; Light, N = 13 embryos; uninjected WT, N = 11 embryos.

**f,g,** Embryo-level summaries for uninjected WT (N = 11 embryos) and injected embryos under dark (N = 10 embryos) versus light activation (N = 13 embryos): mean *her12* level (f) and *ptch2* gradient amplitude (g). Points, embryos; large symbols, mean ± s.d. across embryos.

**h,i,** Representative dorsal views of a 10-µm ventral optical slab at 12 ss showing *TgBAC(ptch2:Kaede)* signal. In wild type (h), lateral cells with reduced *ptch2:Kaede* are indicated (arrowheads); under uniform Notch activation (i), these low-*ptch2:Kaede* lateral cells are reduced in prevalence. Images are representative; the same phenotype was observed in all WT embryos (N = 9) and uniform-Notch embryos (N = 3). Scale bars, 20 µm.

**Extended Data Fig. 9. Fate-configuration quantification for lineage-traced and timed Shh-inhibition datasets.**

**a,** Cell-type fractions for lineage-traced ventral progenitors at 12 ss (p3, pMN, p2, p1). Bars, mean; error bars, s.d. across embryos; dots, embryos. Text indicates the entropy of the fate distribution (bits).

**b,** Trajectory-based fate information for decoding 12 ss fate *F*_12_ across stages. Dark green, decoding from DV-position history *d*_0:*t*_ alone; black, decoding from the combined DV-position and Shh-response histories (*d*_0:*t*_, *g*_0:*t*_).

**c, d,** Sampling metrics for the timed Shh-inhibition dataset: total cell number per embryo (c) and p3–p1 cell number per embryo (d) across inhibition windows (Control, <0 ss, 0–5 ss, 5–10 ss). Dots, embryos; horizontal bars, mean ± s.d. across embryos.

**e,** Cell-type fractions (including p3-p1 and p0) across timed Shh-inhibition windows. Bars, mean; error bars, s.d. across embryos; dots, embryos. Asterisks indicate significant differences versus control for the indicated cell type.

**f,** Entropy of the p3–p1 fate distribution (bits) across inhibition windows. Dots, embryos; large symbols, mean ± s.d. across embryos. Asterisks indicate significant differences versus control.

**g,** Fate-information-to-entropy ratio (FI/entropy) across inhibition windows (definition in Supplementary Note 10). Dots, embryos; large symbols, mean ± s.d. across embryos.

**h,** Marker expression level within each ventral domain under timed Shh inhibition. Bars, mean; error bars, s.d. across embryos; dots, embryos. Asterisks indicate significant differences versus control for the indicated marker/domain combination.

**i, j,** Within-domain positional variability quantified as the s.d. of DV position within each domain. Values are plotted as a function of the mean DV position of the domain (i) and as a function of the domain’s cell-type fraction (j). Points, embryos; error bars, mean ± s.d. across embryos; symbol shapes denote domains (p0–p3) as indicated.

Sample sizes for timed Shh inhibition (c–j): Control N = 6 embryos, n = 3,000 cells; <0 ss N = 6 embryos, n = 2,456 cells; 0–5 ss N = 7 embryos, n = 3,341 cells; 5–10 ss N = 7 embryos, n = 3,407 cells.

**Extended Data Fig. 10. Transcriptional dynamics of dorsoventral markers during neurulation.**

**a,** UMAP feature plots for reclustered spinal cord cells from the ZMAP single-cell atlas at 10 hpf (∼0 ss), 12 hpf (∼6 ss) and 14 hpf (∼10 ss). Left, ventral progenitor-class annotation using canonical dorsoventral (DV) markers (p3, red; pMN, orange; p2, green; p1, blue; p0, purple). Right, expression of representative DV transcription-factor markers (italics denote gene names): *nkx2.2a*, *nkx2.2b*, *nkx2.9*, *olig2*, *lbx2*, *nkx6.1*, *nkx6.2*, *prdm12b*, *dbx1a*, *dbx1b*, *irx3a*, *pax3a* and *msx1b*.

**b,** Representative confocal cross-sections of multiplex HCR RNA-FISH showing combinatorial transcription-factor staining used to assign ventral progenitor classes (p3–p0; marker combination indicated in the overlay). Images are from embryos staged in hours post fertilization (hpf, 28.5 °C) and mapped onto somite-stage units (ss) by linear conversion: 10–15 hpf → 0–12 ss in 1-hpf steps (10, 11,…, 15 hpf maps to 0, 2.4,…, 12 ss). Top, example embryo with predominant pMN-marker (*olig2*) signal; bottom, example embryo with detectable expression of multiple p3–p0 markers at this stage. Scale bar, 20 µm.

**c,** Ventral-class fractions across neurulation stages quantified from multiplex HCR RNA-FISH as in **b**, using the same hpf→ss mapping (10–15 hpf → 0–12 ss). Curves show mean ± s.d. across embryos. Sample sizes by hpf: 10 hpf (N = 5 embryos; n = 3,268 cells), 11 hpf (N = 6; n = 4,663), 12 hpf (N = 4; n = 2,464), 13 hpf (N = 7; n = 3,931), 14 hpf (N = 6; n = 2,992), 15 hpf (N = 5; n = 2,715).

**d,** Amplitude dynamics of domain-enriched transcription factors quantified from single-gene HCR RNA-FISH across neurulation stages. For each gene and stage, N = 6 embryos (independent sets; one gene stained per embryo). Curves show mean ± s.d. across embryos.

## Supplementary Video Legends

**Supplementary Video 1.**

Lineage reconstructions of ventral spinal-cord progenitors from a wild-type embryo. Left, 3D reconstruction of tracked nuclei within the spinal-cord mask (notochord surface shown for reference) with Shh response overlaid. Right, single-cell Shh response versus DV position across stages (0–12 ss), with points coloured by embryo (stage indicated by the colour bar).

**Supplementary Video 2.**

Neural plate convergence dynamics under morphogenesis perturbations. Transverse sections from time-lapse movies of *MZitga5^-/-^*, wild type, and *cdh2^-/-^* embryos (top to bottom), illustrating faster, normal, and slower convergence, respectively. A representative lineage originating from the lateral neural plate is marked in each embryo.

**Supplementary Video 3.**

Lineage reconstructions with fate mapping. Left, 3D lineage reconstruction from a fate-mapped wild-type embryo with nuclei coloured by the assigned 12-ss ventral progenitor fate (p3, pMN, p2, p1; notochord surface shown for reference). Right, Shh response versus DV position for the same tracked cells, with points coloured by final fate (stage indicated by the colour bar).

