## Supplementary Figures for "Cells specify fate within an optimal window of positional information determined by morphogenesis"

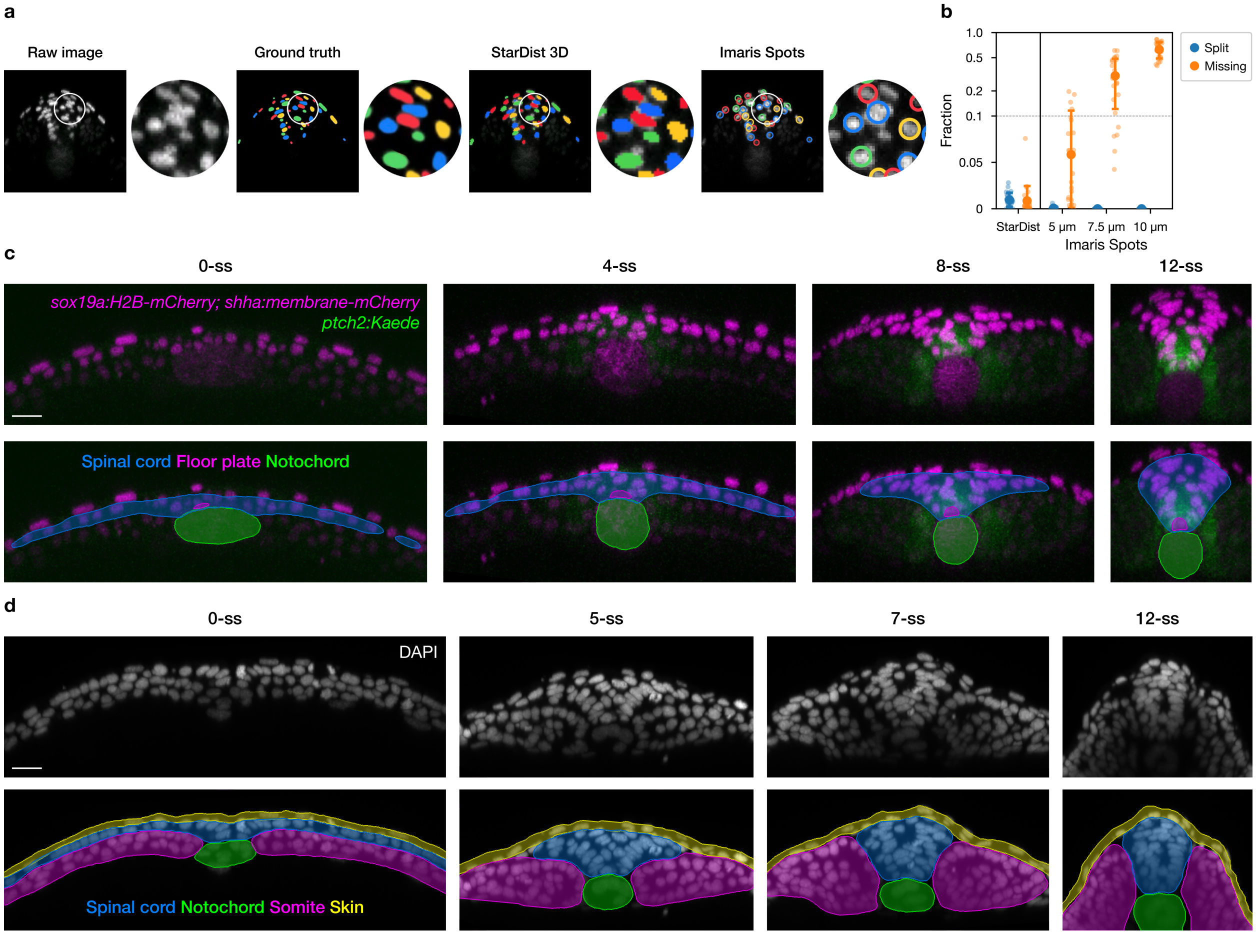


### Extended Data Fig. 1 | Deep-learning segmentation of nuclei and reference tissues

**a,** Representative confocal transverse optical section from a 10-ss neural tube used for nucleus-segmentation benchmarking (independent test image; not used for model training or validation). From left to right: raw image; manually curated ground-truth nuclei; StarDist-3D nucleus instance segmentation; nuclei detected by Imaris (Spots). Colored overlays indicate individual nucleus instances.

**b,** Benchmark of nucleus-detection error modes for StarDist-3D versus Imaris Spots across Spot diameter settings, evaluated on an independent test set (not used for model training or validation). For each embryo, points show the fraction of ground-truth nuclei classified as split (one ground-truth nucleus matched to >1 prediction) or missing (no prediction matched to a ground-truth nucleus). Large symbols show mean ± s.d. across embryos. The y axis is plotted linearly from 0 to 0.1 and on a log scale above 0.1.

**c,** Representative 3D U-Net tissue segmentation across the live time-lapse window used for lineage analyses (example confocal transverse optical sections; not part of the segmentation training/validation datasets). Top row: confocal transverse optical sections at 0, 4, 8 and 12 ss from embryos carrying *Tg(sox19a:H2B-mCherry)* and *Tg(shha:mem-mCherry)* (magenta) together with the Shh-response reporter *TgBAC(ptch2:Kaede) (green)*. Bottom row:

corresponding tissue masks (spinal cord, floor plate and notochord) overlaid on the same sections. Scale bar, 20 µm.

**d,** Representative 3D U-Net segmentation of reference tissues in fixed embryos stained with DAPI across stages (example confocal optical sections; not part of the segmentation training/validation datasets). Top row: DAPI optical sections at 0, 5, 7 and 12 ss. Bottom row: corresponding masks for spinal cord, notochord, somites and skin overlaid on the same sections. Scale bar, 20 µm.

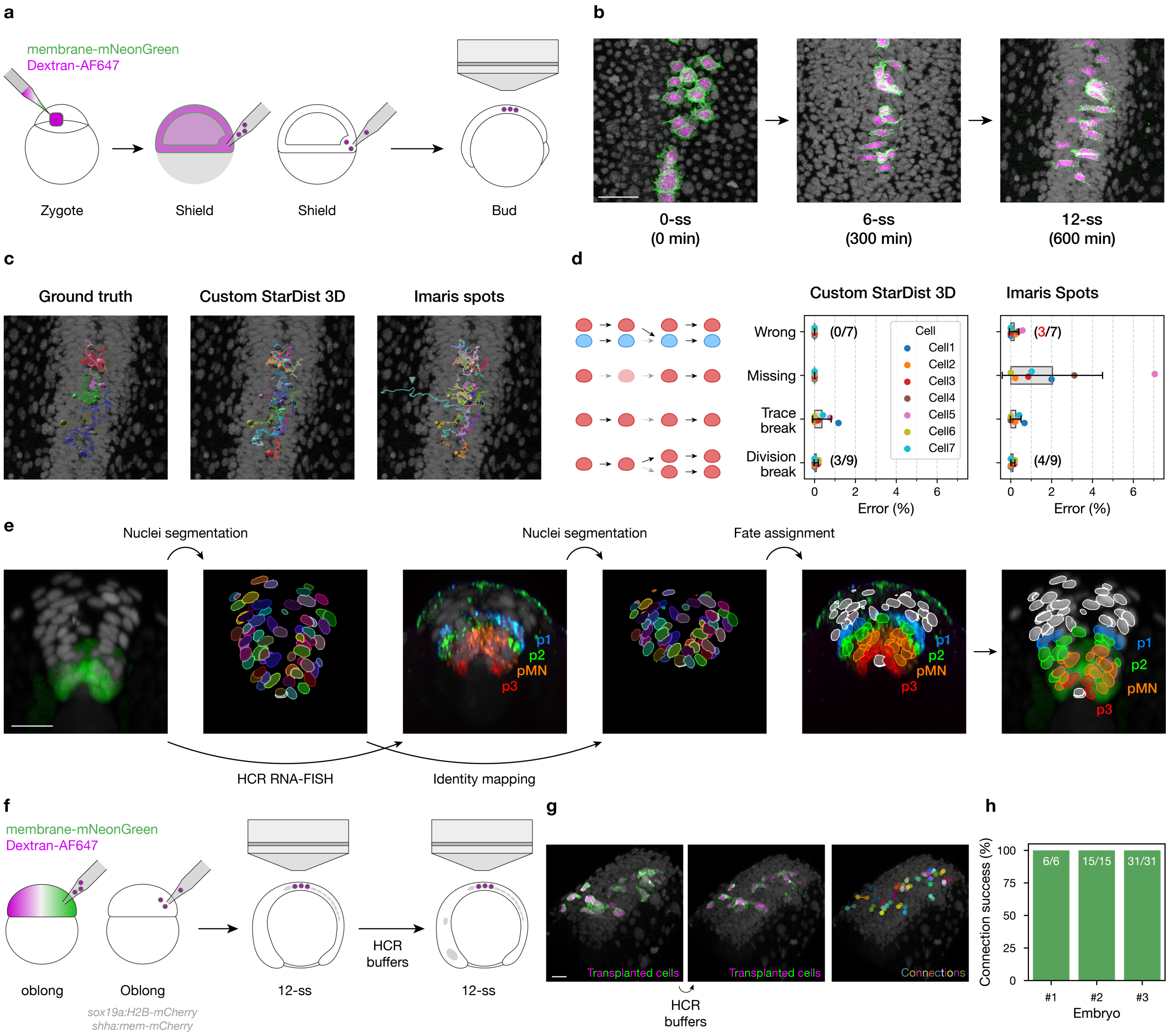


### Extended Data Fig. 2 | Quality control for lineage reconstruction and live-to-fixed fate mapping

**a,** Schematic of sparse cell transplantation used to generate "ground-truth" donor cells for quality control of lineage reconstruction. Donor cells were labelled with membrane-mNeonGreen and Dextran-AF647.

**b,** Representative confocal optical sections from a transplanted embryo at 0-, 6-, and 12-somite stages (0, 300, and 600 min) showing sparse donor-cell labelling that enables unambiguous manual tracking across frames. Scale bar, 20 μm.

**c,** Comparison of automatically reconstructed donor-cell trajectories with manually curated ground-truth trajectories. Tracks are shown only for donor cells. A representative misconnection is highlighted (cyan): an Imaris track that is incorrectly linked into the donor lineage set.

**d,** Accuracy of automatic frame-to-frame linking for donor-cell tracking using Brownian-motion linking (maximum step size 5 µm; no gap closing), comparing segmentations from Custom StarDist-3D versus Imaris Spots (benchmark dataset only; not used for model training/validation). Error categories are illustrated schematically (left). For each method, points show per-cell linking error (%) for the indicated error modes; counts indicate the number of affected cases. Manual curation produced 100% correct linking for the evaluated donor-cell set.

**e,** Workflow for live-to-fixed fate mapping. After live imaging, embryos were fixed and stained by multiplex HCR RNA-FISH to assign each tracked nucleus to a ventral progenitor fate domain (p3, pMN, p2, p1) in the fixed specimen, then mapped back to the terminal frame of each live lineage.

**f,** Schematic of sparse-labelling strategy used to quality-control live-to-fixed mapping: sparse donor-cell labels are imaged live, then re-identified after fixation and HCR processing to verify correspondence across modalities.

**g,** Representative confocal optical sections showing sparse labels in the live embryo, after HCR processing, and the resulting correspondence used for identity matching. Scale bar, 20 µm.

**h,** Live-to-fixed identity matching success rate across embryos. Bars show the number of successfully matched cells out of the total evaluated cells per embryo.

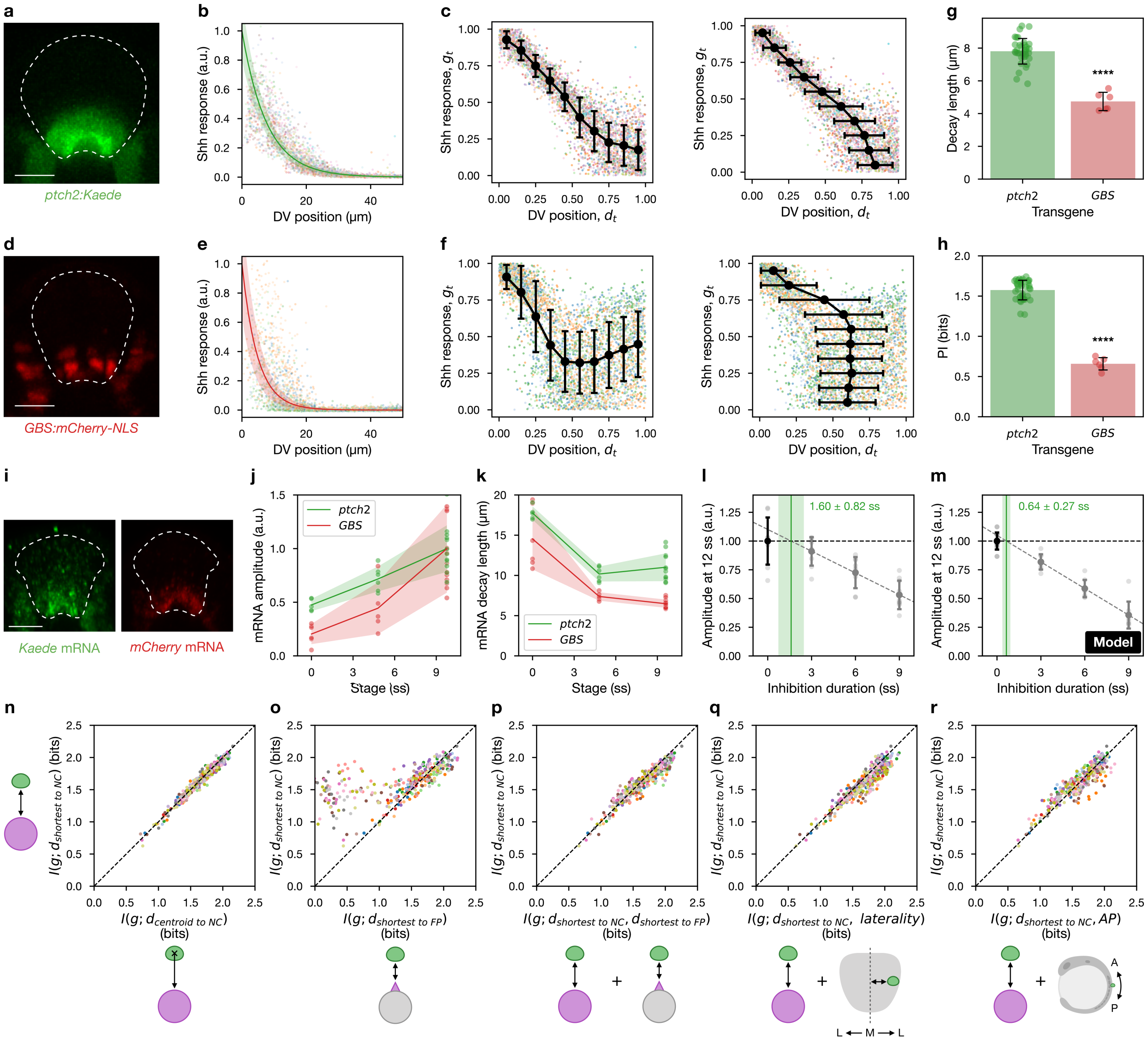


**Extended Data Fig. 3 | Selection of Shh-response and dorsoventral (DV) position metrics and an upper bound on reporter delay**

**a–c,** Analysis of embryos expressing the Shh-response reporter *TgBAC(ptch2:Kaede)* (N = 36 embryos; n = 26,557 cells).

**d–f,** Analysis of embryos expressing *Tg(8xgli-Xla.Cryaa:NLS-mCherry)* (GBS) (N = 6 embryos; n = 4,666 cells).

For the cell-level scatter plots in **b,c,e,f,** we randomly subsampled 4,000 cells per reporter to balance cell density for visual comparison.

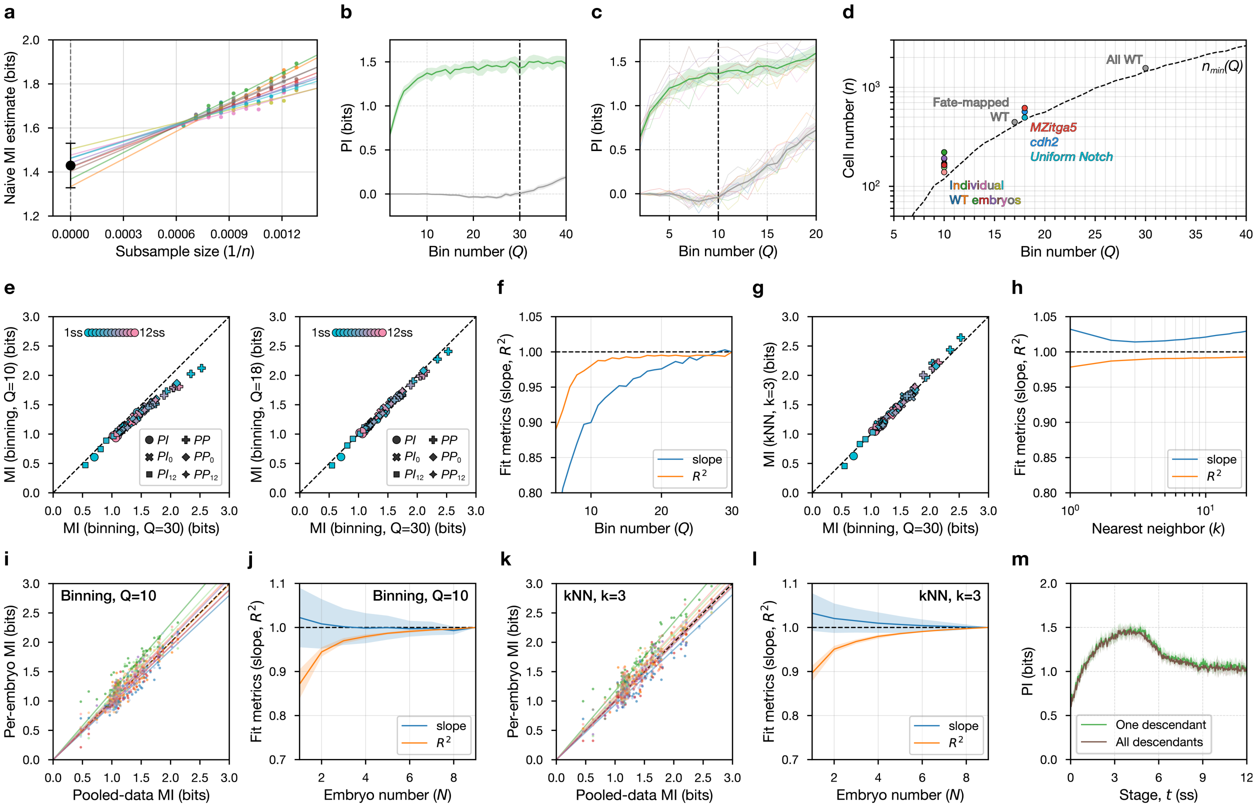


### Extended Data Fig. 4 | Quality control of mutual-information estimation, binning resolution and sample size

**a,** Example of direct subsampling–extrapolation debiasing for a pooled wild-type estimate (illustrated for PI at 4 ss). Points show the naïve plug-in mutual information computed on nested subsamples of size $n$, plotted versus $1/n$. Coloured lines show independent linear fits (“chains”) across repeated subsampling/fit realizations; the black point at $1/n = 0$ indicates the extrapolated infinite-sample intercept, reported as mean with 95% CI.

**b,** Example of dependence on bin number $Q$ for pooled positional information at 4 ss. Green line, debiased estimate (mean with 95% CI across debiasing chains). Grey line, shuffle-null control generated by permuting Shh response relative to DV position 30 times (mean with 95% CI across permutations, using the same estimator settings). Vertical dashed line marks the bin number used for pooled estimates ($Q = 30 \sim Q_{max}(n_{pooled,WT})$).

**c,** Example of dependence on bin number $Q$ for per-embryo positional information at 4 ss. For each $Q$, positional information was computed separately in each embryo and then summarized across embryos. Green line, mean with 95% CI across embryos; grey line, shuffle-null control (mean with 95% CI across embryos, using within-embryo permutations). Vertical dashed line marks the bin number used for per-embryo estimates ($Q = 10 \sim Q_{max}(n_{embryo,WT})$).

**d,** Lookup curve $n_{min}(Q) \sim Q_{max}(n)$ across bin numbers $Q$ (dashed), summarizing the minimum sample size required for shuffle-null bias to be negligible under the chosen estimator settings.

Coloured points show the $(Q, n)$ pairs used across datasets/conditions (including wild type and perturbations), confirming that all analyses were performed above the corresponding $n_{min}(Q)$ threshold.

**e,** Effect of binning resolution on binned mutual-information estimates: comparison of estimates obtained using reduced bin numbers $Q$ versus a high-resolution reference ($Q = 30$). Each point corresponds to a matched estimate (same stage/MI type), plotted against the $Q = 30$ value; dashed line, identity.

**f,** Summary of the binning-resolution benchmark in **e**, reporting agreement metrics (slope and $R^2$) as a function of $Q$, evaluated relative to the $Q = 30$ reference.

**g,** k-nearest-neighbour (kNN) mutual-information estimates benchmarked against the $Q = 30$ binned reference. Points show matched estimates across stage/MI type; dashed line, identity.

**h,** Dependence of kNN estimates on neighbourhood size k, summarized as agreement metrics relative to the $Q = 30$ reference (as in **f**), to assess estimator stability across $k$.

**i,** Agreement between pooled and per-embryo binned estimates: per-embryo mutual information estimates (computed separately in each embryo) plotted against the corresponding pooled-data estimates (pooling embryos at each stage after within-embryo rank normalization) (both estimate use Q=10). Dashed line, identity.

**j,** Summary of per-embryo-versus-pooled agreement in **i** as a function of the number of embryos pooled ($N$) ($Q = 10$), reporting slope and $R^2$.

**k,** Agreement between pooled and per-embryo binned estimates as in **i** but using kNN method (both estimates use $k = 3$).

**l,** Analogous per-embryo-versus-pooled comparison for kNN mutual-information estimates ($k = 3$), plotted as in **j**; dashed line, identity.

**m,** Comparison of positional-information dynamics computed by retaining one descendant per lineage after division ($n$ = 1,561 lineages throughout) versus retaining both descendants after division ($n$ increases from 1,561 to 2,099).

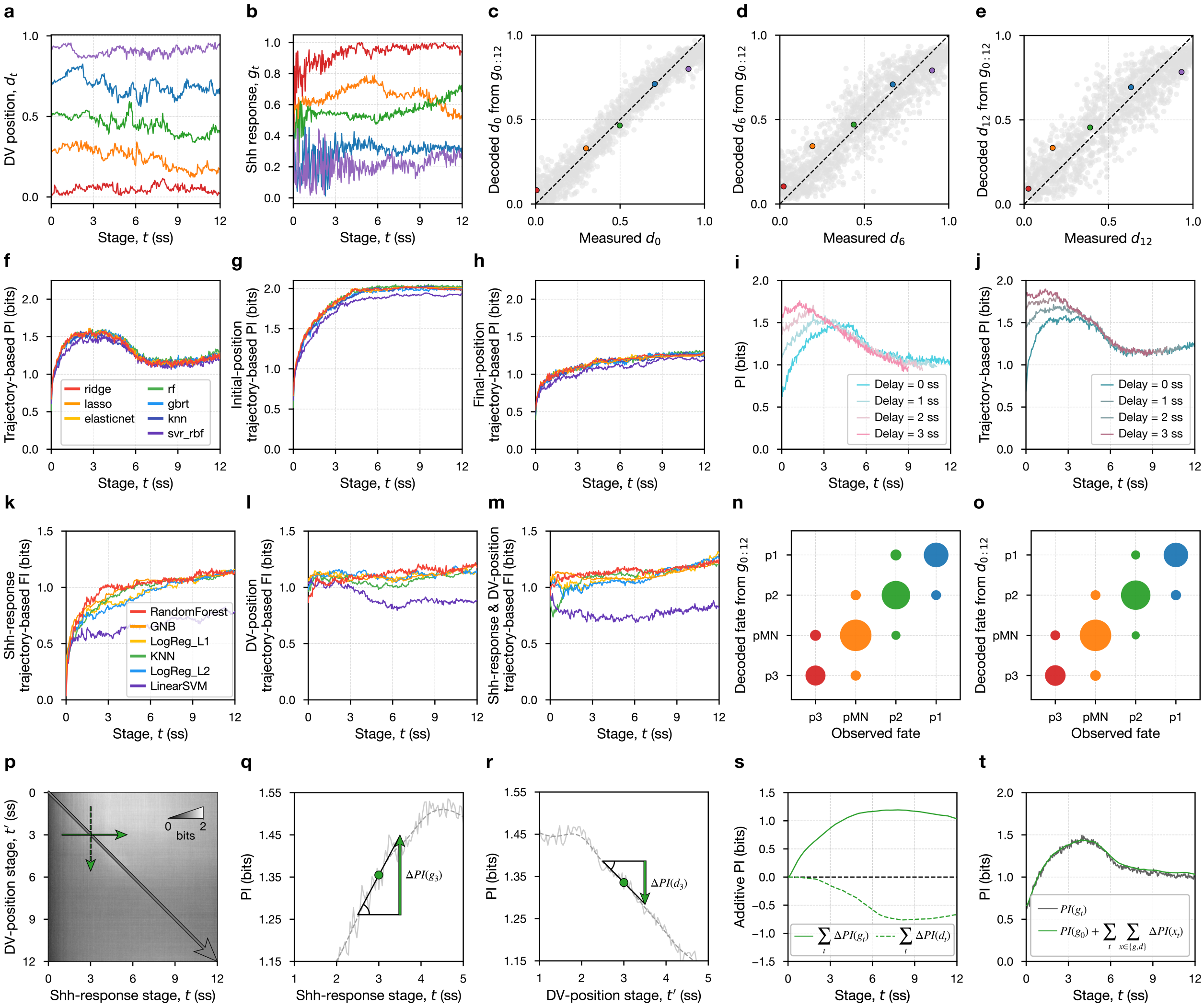


**Extended Data Fig. 5 | Benchmarking decoder-based positional and fate information and testing robustness to Shh-response delay**

**a,b,** Example single-lineage time traces used for decoder benchmarking. a, DV position $d_t$ trajectories for a subset of lineages. **b,** Corresponding Shh-response histories $g_t$ for the same lineages.

**c–e,** Example decoder performance for predicting DV position at different target stages from Shh-response history. Scatter plots compare decoded versus measured DV position for c, $d_0$, d, $d_6$, and e, $d_{12}$ using a ridge-regression decoder; dashed line indicates identity.

**f–h,** Benchmarking alternative decoders for extracting trajectory-based positional information from Shh-response history $g_{0:t}$ about three DV-position targets (Supplementary Note 5). Curves show decoder-based positional information versus stage $t$ for predicting f, instantaneous DV position $d_t$, g, initial DV position $d_0$, and h, final DV position $d_{12}\}$. Methods shown include ridge, lasso, elastic net, random forest, gradient-boosted trees, kNN, and SVR (legend).

**i,j,** Testing robustness to effective Shh-response delay. Positional information dynamics computed after shifting the Shh-response input by an assumed delay (0–3 ss): **i,** instantaneous positional information $PI(g_t)$; **j,** trajectory-based positional information from histories $PI(g_{0:t})$. Allowing delayed readout does not remove the late decline in positional information.

**k–m,** Decoder-based trajectory-based fate information about fate at 12 ss, $F_{12}$, using histories up to stage $t$. Curves show fate information extracted from k, Shh-response histories $g_{0:t}$, **l,** DV-position histories $d_{0:t}$, and m, joint histories $(g_{0:t}, d_{0:t})$, benchmarked across decoder classes as in **f–h** (legend).

**n,o,** Fate-decoding performance of the retained decoder (Random Forest) using the full history to 12 ss. Bubble plots compare decoded (y-axis) versus observed (x-axis) $F_{12}$ using **n**, Shh-response history $g_{0:12}$ or **o**, DV-position history $d_{0:12}$. Bubble area indicates the fraction of cells.

**p–t,** Local "PI gain" decomposition on the cross-temporal PI surface (Supplementary Note 6). **p,** Example illustrating how local changes near an anchor stage (example shown at 3 ss) are decomposed into a Shh-response contribution and a DV-position contribution. **q,** Shh-response contribution $\Delta PI(g_t)$ estimated from a local slope along the Shh-response axis at fixed target stage. **r,** DV-position contribution $\Delta PI(d_t)$ estimated from a local slope along the DV-position axis at fixed Shh-response stage. **s,** Cumulative sums of $\Delta PI(g_t)$ and $\Delta PI(d_t)$ across stages, showing their opposing contributions to the net change in positional information. **t,** Reconstructing the positional information time course by adding the cumulative contributions in **s** to the initial positional information $PI(g_0)$, and comparing with the measured pooled positional information trajectory.

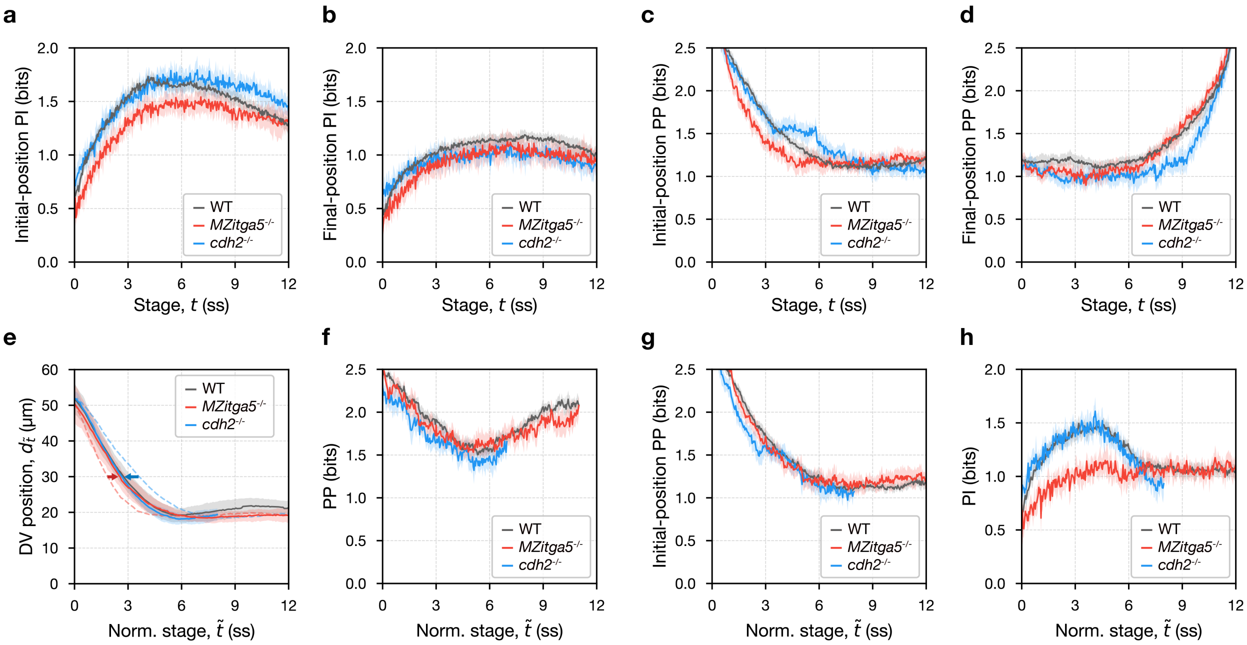


**Extended Data Fig. 6 | Shh-response profile fits, positional information, and positional persistence under morphogenesis perturbations**

**a,b,** Cross-temporal positional information from Shh response, summarized as cuts through $PI_{t'}(g_t)$. **a,** information about the initial DV position, $PI_0(g_t)$. **b,** information about the final DV position, $PI_{12}(g_t)$. Curves show mean with 95% CI for each genotype.

**c,d,** Cross-temporal positional persistence, summarized as cuts through $PP_{t'}(d_t)$. **c,** persistence about the initial DV position, $PP_0(d_t)$. **d,** persistence about the final DV position, $PP_{12}(d_t)$. Curves show mean with 95% CI for each genotype.

**e,** Mean DV-position dynamics plotted versus morphogenesis-normalized stage, $\tilde{t}$ (Supplementary Note 8). Solid lines, trajectories versus morphogenesis-normalized stage; dashed lines, the same trajectories versus original somite stage. Curves show mean ± s.d. across embryos.

**f,g,** Positional persistence versus morphogenesis-normalized stage. **f,** 1-somite-stage interval persistence, $PP_{t+1}(d_t)$. **g,** persistence about the initial DV position, $PP_0(d_t)$. Curves show mean with 95% CI.

**h,** Positional information about instantaneous DV position plotted versus morphogenesis-normalized stage, $PI(g_t)$.

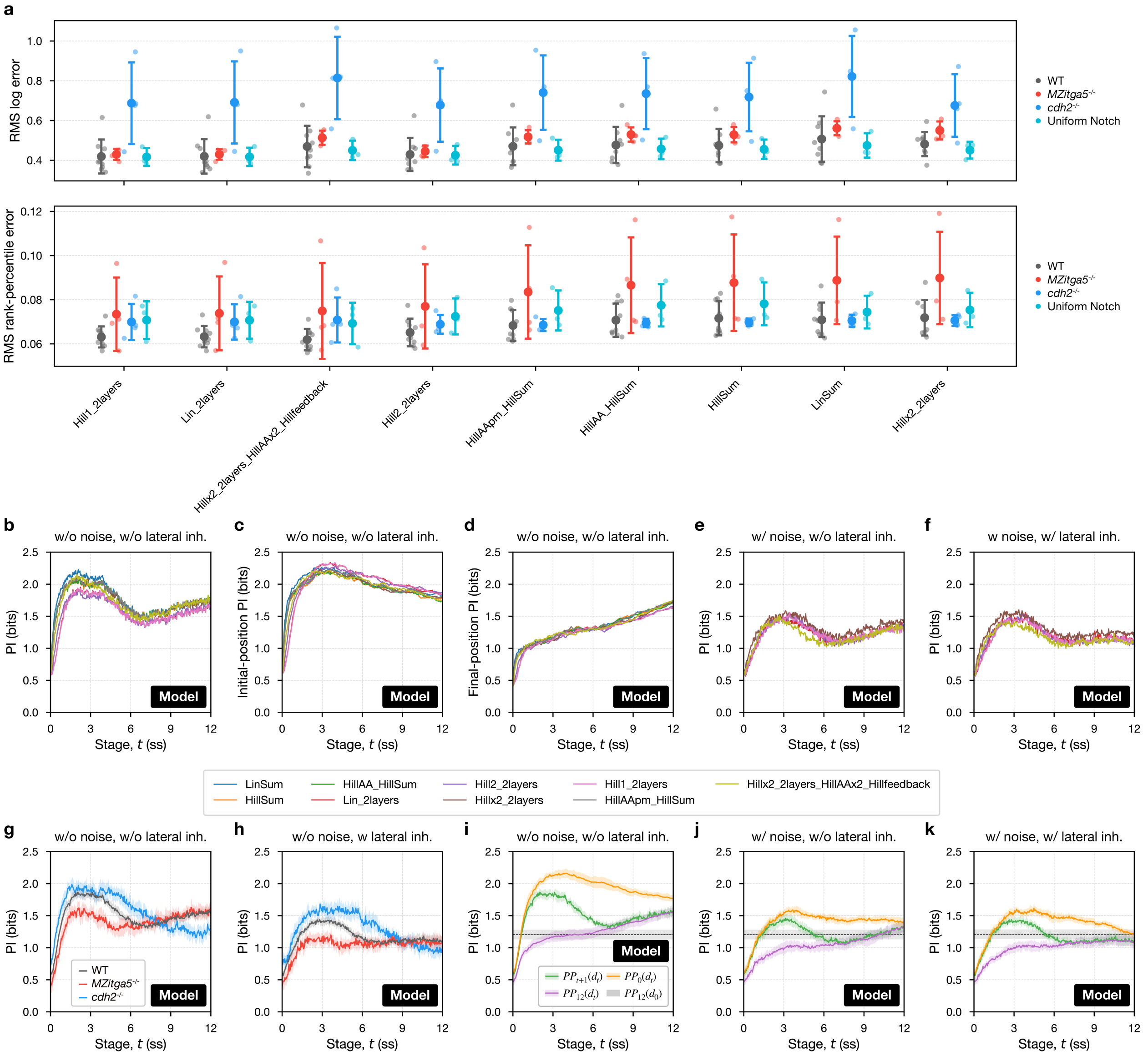


**Extended Data Fig. 7 | Benchmarking and using the hybrid forward model**

**a,** Model-selection summary across alternative sensing logics. For each sensing logic, performance is shown for the single best-performing parameterization (selected using wild type and then evaluated across conditions). Top, root-mean-square (RMS) error in log space between predicted and measured Shh response in absolute units. Bottom, RMS error after within-embryo rank normalization (rank-percentiles). Points, embryos; large symbols, mean ± s.d. across embryos. Models are ordered by the maximum (worst) mean RMS rank-percentile error across conditions.

**b–d,** Positional information computed from model-predicted Shh responses using alternative sensing logics without added noise. **b,** instantaneous positional information, $PI(g_t)$. **c,** positional information about initial DV position, $PI_0(g_t)$. **d,** positional information about final DV position, $PI_{12}(g_t)$. Curves show mean with 95% CI.

**e,f,** Positional information for the same model set after adding either **e**, stage-independent ("constant") noise to the predicted Shh response or **f,** a neighbour-coupled lateral-inhibition module that generates cell-to-cell heterogeneity in Shh sensitivity. Display conventions as in **b–d**.

**g,h,** Positional information computed from model-predicted Shh responses when substituting experimentally measured DV trajectories from morphogenesis-perturbed embryos. **g,** no added noise. **h,** lateral-inhibition setting. Curves show mean with 95% CI.

**i–k,** One-dimensional slices through cross-temporal positional information, $PI_{t'}(g_t)$, shown for **i,** no added noise, **j,** constant-noise setting, and **k,** lateral-inhibition setting. Curves show mean ± s.d. across embryos; mutual-information estimates are summarized as mean with 95% CI within each embryo.

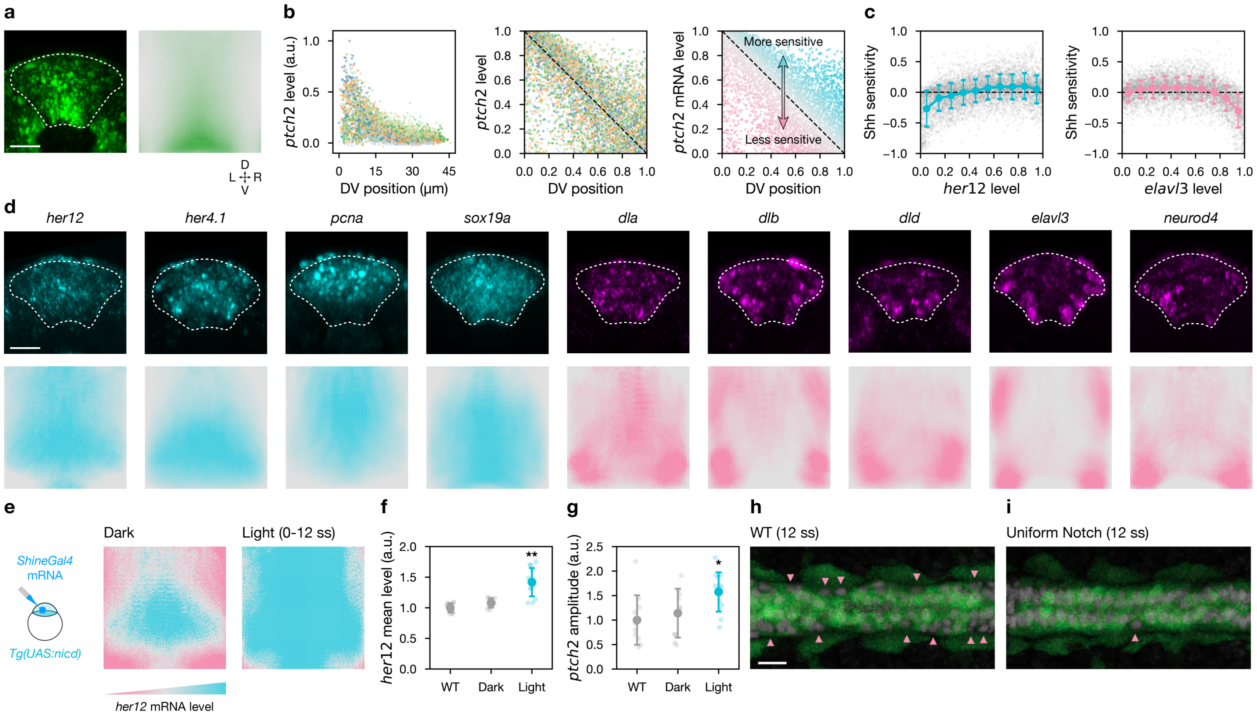


### Extended Data Fig. 8 | Notch-state correlates of Shh sensitivity and effects of uniform Notch activation

**a,** Confocal optical cross-section showing endogenous *ptch2* mRNA at 10 ss (left) and the corresponding DV–LR (dorsal-ventral–left-right) mean expression profile across the spinal cord (right). Dashed outline, spinal cord. Scale bar, 20 µm.

**b,** Definition of Shh sensitivity from single-cell *ptch2* expression at 10 ss, illustrated using the *her12* dataset (N = 6 embryos; n = 5,561 cells). Left, *ptch2* mRNA versus DV position for individual cells (points, cells; colours, embryos). Middle, rank-normalized *ptch2* level in [0,1]. Right, Shh sensitivity defined as the residual (deviation) from the expected DV trend (dashed line), with points coloured by sensitivity (higher = more Shh-sensitive; lower = less Shh-sensitive).

**c,** Relationship between Shh sensitivity and representative Notch-associated targets at 10 ss: *her12* (Notch-associated/progenitor; left; N = 6 embryos; n = 5,561 cells) and *elavl3* (neurogenic; right; N = 6 embryos; n = 5,052 cells). Points, cells; black line, mean Shh sensitivity within 0.1-wide expression bins; error bars, s.d. across cells within bins.

**d,** DV–LR expression profiles (top, confocal optical sections; bottom, corresponding DV–LR mean maps) for Notch-associated targets at 10 ss used in this study: *her12* (N = 6; n = 5,561), *her4.1* (N = 6; n = 5,584), *pcna* (N = 5; n = 4,557), *sox19a* (N = 5; n = 4,584) (cyan; progenitor/Notch-associated) and *dla* (N = 5; n = 4,470), *dlb* (N = 5; n = 4,651), *dld* (N = 5; n = 4,453), *elavl3* (N = 6; n = 5,052), *neurod4* (N = 5; n = 4,528) (magenta; neurogenic/Notch-negative). Dashed outlines, spinal cord. Scale bars, 20 µm.

**e,** DV–LR mean profiles of *her12* mRNA at 12 ss in control (dark) versus uniform Notch activation (light, 0–12 ss) embryos generated by ShineGal4-driven NICD expression (ShineGal4 mRNA injection in *Tg(UAS:nicd)* embryos). Dark, N = 10 embryos; Light, N = 13 embryos; uninjected WT, N = 11 embryos.

**f,g,** Embryo-level summaries for uninjected WT (N = 11 embryos) and injected embryos under dark (N = 10 embryos) versus light activation (N = 13 embryos): mean *her12* level (f) and *ptch2* gradient amplitude (g). Points, embryos; large symbols, mean ± s.d. across embryos.

**h,i,** Representative dorsal views of a 10-µm ventral optical slab at 12 ss showing *TgBAC(ptch2:Kaede)* signal. In wild type (h), lateral cells with reduced *ptch2:Kaede* are indicated (arrowheads); under uniform Notch activation (i), these low-*ptch2:Kaede* lateral cells are reduced in prevalence. Images are representative; the same phenotype was observed in all WT embryos (N = 9) and uniform-Notch embryos (N = 3). Scale bars, 20 µm.

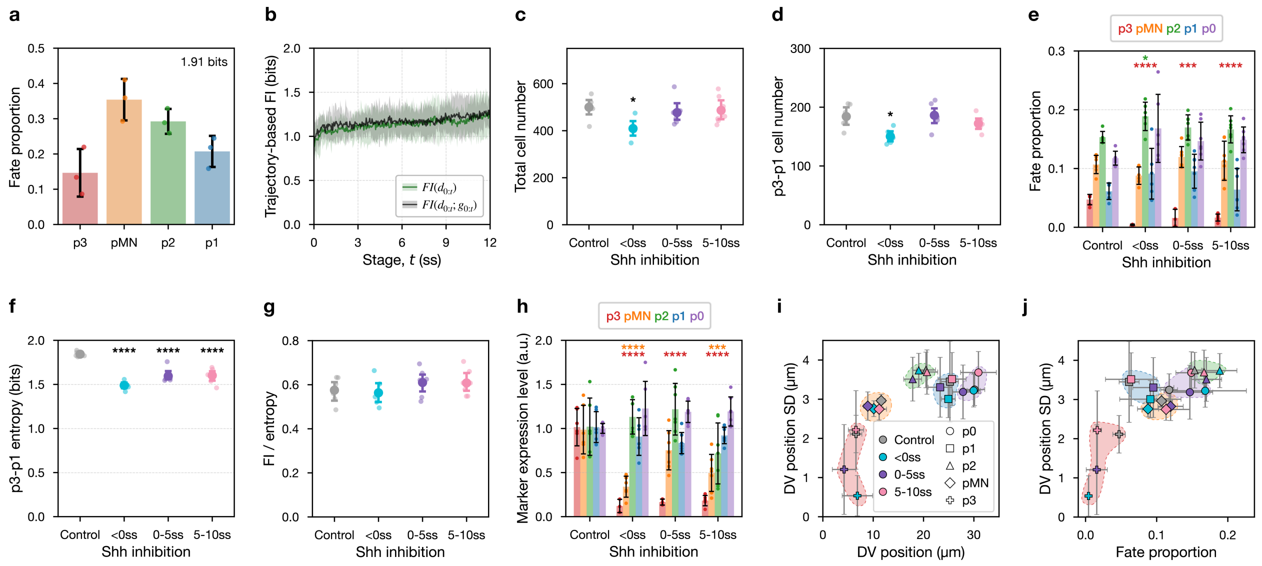


### Extended Data Fig. 9 | Fate-configuration quantification for lineage-traced and timed Shh-inhibition datasets

**a,** Cell-type fractions for lineage-traced ventral progenitors at 12 ss (p3, pMN, p2, p1). Bars, mean; error bars, s.d. across embryos; dots, embryos. Text indicates the entropy of the fate distribution (bits).

**b,** Trajectory-based fate information for decoding 12 ss fate $F_{12}$ across stages. Dark green, decoding from DV-position history $d_{0:t}$ alone; black, decoding from the combined DV-position and Shh-response histories $(d_{0:t}, g_{0:t})$.

**c, d,** Sampling metrics for the timed Shh-inhibition dataset: total cell number per embryo (c) and p3–p1 cell number per embryo (d) across inhibition windows (Control, <0 ss, 0–5 ss, 5–10 ss). Dots, embryos; horizontal bars, mean ± s.d. across embryos.

**e,** Cell-type fractions (including p3-p1 and p0) across timed Shh-inhibition windows. Bars, mean; error bars, s.d. across embryos; dots, embryos. Asterisks indicate significant differences versus control for the indicated cell type.

**f,** Entropy of the p3–p1 fate distribution (bits) across inhibition windows. Dots, embryos; large symbols, mean ± s.d. across embryos. Asterisks indicate significant differences versus control.

**g,** Fate-information-to-entropy ratio (FI/entropy) across inhibition windows (definition in Supplementary Note 10). Dots, embryos; large symbols, mean ± s.d. across embryos.

**h,** Marker expression level within each ventral domain under timed Shh inhibition. Bars, mean; error bars, s.d. across embryos; dots, embryos. Asterisks indicate significant differences versus control for the indicated marker/domain combination.

**i, j,** Within-domain positional variability quantified as the s.d. of DV position within each domain. Values are plotted as a function of the mean DV position of the domain (i) and as a function of the domain’s cell-type fraction (j). Points, embryos; error bars, mean ± s.d. across embryos; symbol shapes denote domains (p0–p3) as indicated.

Sample sizes for timed Shh inhibition (c–j): Control N = 6 embryos, n = 3,000 cells; <0 ss N = 6 embryos, n = 2,456 cells; 0–5 ss N = 7 embryos, n = 3,341 cells; 5–10 ss N = 7 embryos, n = 3,407 cells.

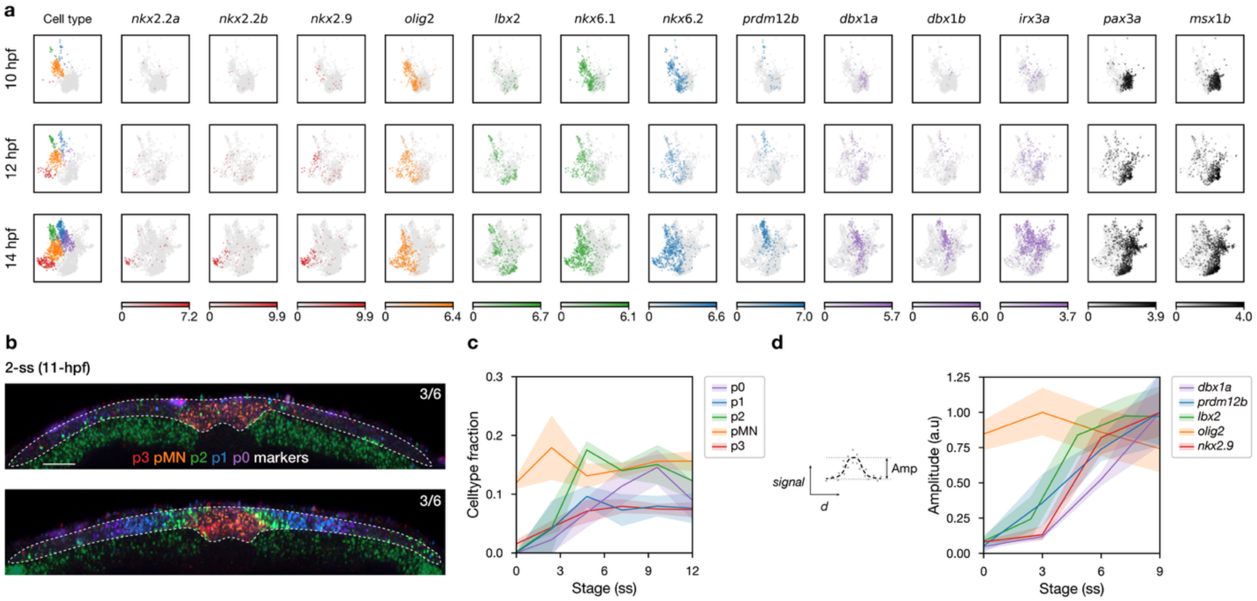


**Extended Data Fig. 10 | Transcriptional dynamics of dorsoventral markers during neurulation**

**a,** UMAP feature plots for reclustered spinal cord cells from the ZMAP single-cell atlas at 10 hpf (~0 ss), 12 hpf (~6 ss) and 14 hpf (~10 ss). Left, ventral progenitor-class annotation using canonical dorsoventral (DV) markers (p3, red; pMN, orange; p2, green; p1, blue; p0, purple). Right, expression of representative DV transcription-factor markers (italics denote gene names): *nkx2.2a*, *nkx2.2b*, *nkx2.9*, *olig2*, *lbx2*, *nkx6.1*, *nkx6.2*, *prdm12b*, *dbx1a*, *dbx1b*, *irx3a*, *pax3a* and *msx1b*.

**b,** Representative confocal cross-sections of multiplex HCR RNA-FISH showing combinatorial transcription-factor staining used to assign ventral progenitor classes (p3–p0; marker combination indicated in the overlay). Images are from embryos staged in hours post fertilization (hpf, 28.5 °C) and mapped onto somite-stage units (ss) by linear conversion: 10–15 hpf → 0–12 ss in 1-hpf steps (10, 11, …, 15 hpf maps to 0, 2.4, …, 12 ss). Top, example embryo with predominant pMN-marker (*olig2*) signal; bottom, example embryo with detectable expression of multiple p3–p0 markers at this stage. Scale bar, 20 µm.

**c,** Ventral-class fractions across neurulation stages quantified from multiplex HCR RNA-FISH as in **b**, using the same hpf→ss mapping (10–15 hpf → 0–12 ss). Curves show mean ± s.d. across embryos. Sample sizes by hpf: 10 hpf (N = 5 embryos; n = 3,268 cells), 11 hpf (N = 6; n = 4,663), 12 hpf (N = 4; n = 2,464), 13 hpf (N = 7; n = 3,931), 14 hpf (N = 6; n = 2,992), 15 hpf (N = 5; n = 2,715).

**d,** Amplitude dynamics of domain-enriched transcription factors quantified from single-gene HCR RNA-FISH across neurulation stages. For each gene and stage, N = 6 embryos (independent sets; one gene stained per embryo). Curves show mean ± s.d. across embryos.
