## Supplementary Information for "Cells specify fate within an optimal window of positional information determined by morphogenesis"

### Supplementary Methods

#### Supplementary Methods 1 | Deep learning segmentation

##### a. Nucleus segmentation

In the time-lapse experiments, neural-progenitor nuclei and notochord membrane signal are both present in the same mCherry channel in embryos carrying *Tg(sox19a:H2B-mCherry)*[1] together with *Tg(-2.4shha-ABC:membrane-mCherry)*[2]. We therefore required a nucleus-segmentation model that remains accurate despite the presence of notochord membrane signal in the same channel. To obtain reliable nuclear ground truth for training, we collected high-resolution snapshot 3D z-stacks from embryos that additionally carried a green membrane marker to improve boundary separation during ground-truth annotation. We then downsampled and augmented these data to match the effective resolution and noise characteristics of the time-lapse setting, and trained StarDist-3D using the mCherry channel as input and nuclear instance masks as output.

###### i. High-resolution imaging and ground truth generation

For ground-truth annotation, we imaged embryos carrying *Tg(sox19a:H2B-mCherry)* and *Tg(-2.4shha-ABC:membrane-mCherry)*, together with *Tg(sox19a:membrane-mNeonGreen)*, which was used to facilitate separation of adjacent nuclei during annotation. We acquired snapshot 3D z-stacks from 36 embryos spanning 0–12 somite stages using a ZEISS LSM Airyscan microscope. After preprocessing, stacks were represented at 0.25 µm × 0.25 µm × 0.25 µm voxel size. We generated nuclear instance ground-truth masks in ilastik[3] via pixel classification followed by object identification, using a small set of manually annotated nuclei as foreground and membrane/background as background.

###### ii. Downsampling and augmentation

To obtain 1 µm × 1 µm × 1 µm training data, each high-resolution stack was converted into 64 low-resolution realizations using phase-offset subsampling. Briefly, the high-resolution volume was partitioned into 4 × 4 × 4 voxel blocks, and each realization was constructed by selecting one fixed relative voxel position within each block (one of 64 possible offsets). This yields 64 paired low-resolution mCherry volumes and matching ground-truth masks per embryo. We applied random spatial shifts and rotations identically to images and ground-truth masks. To increase robustness to differences in acquisition conditions, we additionally applied additive noise, Gaussian blur, and simulated overexposure to the images only, thereby spanning the range of signal quality encountered across time-lapse imaging and fixed-embryo datasets.

###### iii. StarDist-3D training

We trained StarDist-3D (StarDist v0.8.5)[4] using the StarDist 3D ZeroCostDL4Mic notebook[5]. The model input was the augmented mCherry channel (nuclear signal plus notochord membrane signal) and the target was the corresponding nuclear instance ground-truth mask. Training used patch_size = 64 × 200 × 200 (Z × Y × X), batch_size = 2, number_of_steps = 149, n_rays = 192, and initial_learning_rate = 0.0003, for 500 epochs. Every 100 epochs, we switched to a different set of phase-offset downsampled image/ground-truth pairs with independently drawn augmentations. The model used for time-lapse nucleus segmentation was the checkpoint with the lowest validation loss within the final 100 epochs.

##### iv. Quality control

Representative nucleus segmentations produced by the custom-trained StarDist-3D model showed clear separation of adjacent nuclei and fewer obvious failure modes than Imaris Spots, which we used as a widely used baseline approach (Extended Data Fig. 1a). To quantify segmentation error modes, we scored split events (a single reference nucleus containing more than one predicted nucleus) and missing events (no prediction overlapping a reference nucleus) against an ilastik-assisted instance-mask reference set in an independent test dataset that was not used for model training or validation. We benchmarked these error modes against nuclei detected by Imaris Spots across a range of spot-diameter parameters and found that the custom StarDist-3D model had substantially lower split/missing error rates (~1%) than Imaris Spots (typically >5%) across parameter choices (Extended Data Fig. 1b).

Because the reference instance masks were generated by semi-automated ilastik annotation, they can contain systematic errors (for example, occasional merging of adjacent nuclei despite membrane-assisted separation). Therefore, the split/missing fractions should be interpreted as approximate performance metrics relative to this reference, rather than absolute ground-truth error rates. More generally, learning-based segmentation models can in some cases generalize beyond imperfections in training labels and achieve better practical performance than the annotations they were trained on[6,7]. As an independent end-to-end check, transplantation-based lineage-tracing quality control yielded 99.6% correct frame-to-frame connections before manual correction, indicating that residual segmentation and tracking errors are rare at the level relevant for downstream analyses (Extended Data Fig. 2a–d; Supplementary Methods 2).

#### b. Tissue segmentation

Many analyses require accurate spatial relationships between cells and surrounding tissues (for example, distance to the notochord). To delineate these tissues consistently and without manual bias, we trained 3D U-Net–style convolutional neural networks[8] to segment tissue masks from volumetric reference channels.

##### i. Spinal cord / notochord / floor plate segmentation in the *sox19a/shha* transgene context

To obtain tissue masks matched to the live-imaging transgene context, we fixed *Tg(sox19a:H2B-mCherry)*; *Tg(-2.4shha-ABC:membrane-mCherry)* embryos across 0–12 somite stages and performed HCR RNA-FISH for the spinal cord marker *sox19a* (647) and floor-plate markers *shhb*, *foxa1*, and *spon1b* (488). We then used ilastik to classify voxels and generate ground-truth object masks for spinal cord, floor plate, and notochord, leveraging the *shha* transgene signal to aid notochord identification. Exported ilastik object predictions were used as ground-truth targets for network training. Representative segmentations are shown in Extended Data Fig. 1c.

For training, each 3D volume was percentile-normalized by clipping intensities to the 1st–99th percentiles and linearly rescaling to [0,1]. No data augmentation was applied. We trained a 3D U-Net with three downsampling and three upsampling levels (two 3×3×3 conv–batch-norm–ReLU blocks per level; 2×2×2 max pooling for downsampling; 2×2×2 transposed convolutions with skip connections for upsampling), using 32 base feature channels doubling with depth (32→64→128) and a 256-channel bottleneck, followed by a 1×1×1 output convolution. Training used Adam (learning rate $1\times10^{-4}$) with an L1 loss between the predicted mask and the ilastik ground-truth mask. Volumes were split into training (100 z-stacks) and validation (20 z-stacks) sets, and early stopping was applied when validation loss did not improve for 50 epochs. Batch size was 3.

We trained three separate models, each predicting a single tissue mask (spinal cord, notochord, or floor plate) from the same input. The best-performing checkpoints (lowest validation loss) occurred at epoch 152 (spinal cord), epoch 267 (notochord), and epoch 113 (floor plate).

###### ii. Generic tissue segmentation from DAPI reference channels

To generate a model usable in embryos without the *sox19a/shha* transgene context, we fixed embryos and performed HCR RNA-FISH for spinal cord (*sox19a*, 488), notochord (*tbxta*, 647), skin (*krt4*, 647), and somites (*fn1b*, 546), together with DAPI as the structural reference channel. ilastik was used to generate ground-truth object masks for each tissue, and exported object predictions were used directly as training targets. Representative segmentations are shown in Extended Data Fig. 1d. This generic segmentation was used for fixed-embryo analyses requiring consistent DV position quantification, including the fate-mapping and cyclopamine experiments.

We trained a 3D U-Net with the same overall architecture as above, but with a single 4-channel output (spinal cord, notochord, skin, somite) predicted jointly from the DAPI input. Inputs were percentile-clipped (1st–99th) and min–max rescaled to [0,1] per volume. Volumes were split into training (100 z-stacks) and validation (10 z-stacks) sets, and the best-performing checkpoint (lowest validation loss) was selected at epoch 246. This generic segmentation pipeline was used for fixed-embryo analyses, including the datasets underlying Fig. 6.

### Supplementary Methods 2 | Lineage reconstruction

#### a. Time-lapse imaging and staging

###### i. Imaging setup

A single embryo at the bud stage (staged by completion of epiboly and yolk closure) was selected, manually dechorionated, and mounted dorsally for time-lapse imaging. Imaging was performed on a ZEISS LSM980 confocal microscope. We acquired a 303 × 303 × 145 $\mu m^3$ volume at 1 μm × 1 μm × 1 μm voxel size, collecting green and red channels simultaneously with 2× line averaging every 2 min for 10 h (301 timepoints).

During the final ~100 min of imaging (51 frames, around 10 ss), embryos reproducibly rolled posteriorly as tail extension began. To maintain coverage of the same anatomical region, we translated the imaging volume by 1 μm every 2 min to follow this global movement.

Imaging was performed at 23 °C, at which embryos develop at approximately half the rate of standard 28.5 °C conditions. Empirically, embryos reached 12 ss after 10 h of imaging at 23 °C, corresponding to ~15 hpf at 28.5 °C. We therefore treat 10 h at 23 °C as approximately equivalent to 5 h at 28.5 °C. For stage alignment, we assumed an approximately constant somite formation rate over the imaging window and defined somite stage (ss) using a linear conversion of 1 ss per 50 min.

#### b. Image preprocessing and segmentation

We used Fiji's 3D Drift Correction to remove global sample drift. After drift correction, we cropped the dataset to retain only the region that was fully captured throughout the entire imaging window. We then segmented (1) nuclei using the StarDist-3D model described in Supplementary Methods 1a and (2) the notochord using the 3D U-Net model described in Supplementary Methods 1b-i. Segmentation outputs were imported into Imaris as Surfaces, with nuclei and notochord represented as separate objects.

#### c. Lineage reconstruction in Imaris

##### i. Automatic tracking

We performed lineage reconstruction in Imaris using automatic Brownian-motion tracking (distance-based linking). We set the maximum allowed displacement to 5 µm and did not allow gap closing. This displacement threshold is smaller than the short-axis diameter of a nucleus and typically yielded ~100–150 trajectories per embryo after automatic tracking.

##### ii. Manual curation

We manually curated tracks by connecting fragments to reconstruct complete lineages, yielding >150 and up to ~220 complete lineages per embryo. We ensured that each starting cell contributed at least one lineage to the final time point unless its descendants exited the imaging volume. Manual curation time varied across embryos (~6–12 h per embryo).

After curation, floor-plate cells were identified and excluded based on their medial location and low *ptch2:Kaede* signal at the final time point.

##### iii. Handling segmentation errors

When segmentation errors occurred, we corrected them when feasible. For correctable split errors (one nucleus segmented into two objects), we merged the objects in Imaris and reconnected the track. For errors that were not readily correctable (for example, two nuclei merged into one object or a missing object), we left a gap at that time point and connected the track segments before and after the gap. For gaps caused by missing or incorrect segmentation, we imputed missing scalar measurements by linear interpolation (for example, $g = (g_{\text{prev}} + g_{\text{next}})/2$).

#### d. Output features for downstream analyses

We exported, for each lineage $i$ at each time point $t$: (1) the mean Kaede-channel intensity, used as Shh response $g_{i,t}$; (2) the shortest 3D distance from the nuclear surface to the notochord surface, used as DV position $d_{t,i}$; and (3) the nuclear centroid coordinates $(x_{i,t}, y_{i,t}, z_{i,t})$, used for neighbourhood-based analyses and lateral-inhibition simulations (Supplementary Note 8).

#### e. Quality control of tracking fidelity

##### i. Consistency checks from acquisition parameters

Time-lapse imaging was acquired at 2-min intervals. Under this time interval, frame-to-frame displacements were small (mean 1.23 ± 0.70 µm; median 1.10 µm), well below a typical nuclear diameter. This supports conservative distance-based linking and reduces the likelihood of identity swaps. Incorrect links typically introduced a break in the lineage graph, which became apparent and was fixed during manual curation.

##### ii. Transplantation-based ground-truth test

To directly assess lineage reconstruction accuracy, we performed cell transplantation to generate a small number of sparsely labelled donor cells that can be followed unambiguously by their unique label over time (Extended Data Fig. 2a,b). In a representative dataset, we followed 7 donor-derived lineages that collectively divided into 16 cells (3,718 frame-to-frame connections). We then applied the full tracking pipeline while withholding the unique-label channel and compared the resulting tracks to the donor-cell ground truth (Extended Data Fig. 2c). Automatic tracking produced highly accurate frame-to-frame connections when using StarDist-3D instance segmentations (99.6%), and manual curation resolved the remaining rare errors, yielding 100% correct reconstruction for the evaluated donor-lineage set (Extended Data Fig. 2d).

##### iii. Comparison to alternative tracking pipelines

We benchmarked segmentation-based tracking against an Imaris Spots workflow using identical stringent tracking settings (5 µm maximum step; no gaps). While spot-based tracking achieved high overall auto-linking performance (97.5%), it exhibited substantial track loss (2.04%), which greatly reduced the feasibility of manual reconstruction of complete lineages. Moreover, despite identical stringent settings, we observed identity swaps in 3 out of 7 donor-derived lineages during automatic spot-based linking. Such swaps are difficult to detect and correct without an independent label and would compromise downstream trajectory-based analyses. Together, these comparisons indicate that accurate instance segmentation is an essential prerequisite for complete, low-swap lineage recovery in this system, which in turn is required for analyses that rely on continuous signalling histories. Notably, our pipeline exceeds state-of-the-art lineage-tracking pipelines[9,10], which report ~5% identity swaps, whereas we observed 0% swaps in our transplantation-based benchmark (over 3,718 connections). In addition, for comparable trajectory lengths (~300 frames), published pipelines typically require manual curation for >95% of lineages, whereas in our datasets ~1/2–1/3 of lineages required manual edits (embryo dependent), and most breaks arose at cell divisions and were straightforward to correct.

### Supplementary Methods 3 | Fate mapping

#### a. HCR RNA-FISH

Immediately after time-lapse imaging, embryos were fixed overnight in 4% PFA at 4 °C and processed for HCR RNA-FISH. Ventral progenitor domains were assigned using the following marker sets: p3 (*nkx2.2a, nkx2.2b, nkx2.9*; 647 channel), pMN (*olig2*; 546 channel), p2 (*lbx2*; 514 channel) and p1 (*prdm12b*; 488 channel).

#### b. Fate assignment

Because transcription-factor mRNAs can be cytosol-enriched and weakly nuclear, domain identities were assigned manually based on multi-channel HCR signal patterns. We did not use automated classification to keep fate calls conservative and to avoid misassignment in cells with low or ambiguous transcript levels.

#### c. Live-to-fixed identity mapping

To map live-imaged lineages to fixed identities, nuclei from the terminal frame of the time-lapse were manually registered to HCR-stained nuclei in the fixed samples (Extended Data Fig. 2e). Registration proceeded hierarchically using (1) skin nuclei as an outer-shell landmark, (2) floor-plate cells as a single medial row, and (3) the overall neural progenitor population to resolve local correspondences. Each lineage-traced progenitor was then matched to a fixed nucleus, and its fate was assigned from the HCR channels. The combined effort of lineage tracing and fate mapping for one embryo required ~40 h of manual work by one person.

#### d. Quality control of live-to-fixed identity mapping

To validate the identity-mapping procedure, we performed cell transplantation to generate sparsely labelled cells, imaged embryos live at 12 ss, fixed them immediately, processed them through the HCR workflow in buffer-only conditions (no probes or hairpins), and then imaged them again (Extended Data Fig. 2f,g). Live-to-fixed identity mapping was then performed as described above without access to the sparse-label channel. Across three embryos, we correctly matched all cells (52/52; 100%) from the fixed samples back to their live counterparts (Extended Data Fig. 2h).

### Supplementary Notes

#### Supplementary Note 1 | Information-theoretic quantities

##### a. Notation and stage indexing

Throughout this study, we use mutual information to quantify statistical dependence between variables in a common unit (bits). This allows us to compare, on the same scale: (1) how Shh response relates to DV position, (2) how DV position is preserved across developmental stages, and (3) how Shh response or DV position relates to ventral fate (estimation details in Supplementary Note 2).

All stage indices $t$ and $t'$ refer to developmental stage in somite stages (ss), treated as a continuous variable spanning 0–12 ss. For mutual-information analyses, Shh response and DV position are represented as within-embryo, stage-wise rank-normalized variables (Supplementary Note 2a-ii), denoted $g_t$ (Shh response) and $d_t$ (DV position). We also consider their histories, starting from time 0:

$$g_{0:t} \coloneqq (g_0, g_1, \dots, g_t),\ d_{0:t} \coloneqq (d_0, d_1, \dots, d_t).$$

##### b. Positional information

We define positional information as a formalization of Wolpert's idea[11,12]: how much a morphogen readout (here, Shh response) specifies position (here, DV position). For a target stage $t'$, we define **DV positional information** as

$$PI_{t'}(g) \coloneqq I(g;\, d_{t'}),$$

where the Shh-response predictor $g$ can be either a snapshot $g_t$ or a trajectory $g_{0:t}$. Importantly, the Shh-response stage $t$ does not have to be equal to the DV-position stage $t'$; when $t \neq t'$, we refer to these as cross-temporal positional-information quantities.

In the instantaneous case $t' = t$, we typically drop the first subscript:

$$PI(g_t) = PI_t(g_t) = I(g_t; d_t)$$

and for the trajectory-based predictors (Supplementary Note 5),

$$PI(g_{0:t}) = PI_t(g_{0:t}) = I(g_{0:t}; d_t).$$

##### c. Positional persistence

To quantify how strongly DV position is preserved across time, we define **positional persistence** as the mutual information between DV positions at two different stages $t$ and $t'$:

$$PP_{t'}(d_t) \coloneqq I(d_t;\, d_{t'}).$$

Intuitively, high positional persistence indicates that relative DV order is largely preserved between $t$ and $t'$, whereas low positional persistence indicates substantial DV rearrangement over that interval.

Unless stated otherwise, "positional persistence" in the main text refers to a fixed 1-ss interval:

$PP_{t+1}(d_t) = I(d_t;\, d_{t+1})$.

#### d. Fate information

To quantify how strongly an upstream variable predicts ventral fate, we define **fate information** as the mutual information between a predictor $x$ and a discrete fate label $F$ defined at a reference stage:

$FI(x) \coloneqq I(x;\, F)$,

where x may be any snapshot or trajectory variable used in the manuscript, for example $x \in \{g_t, d_t, g_{0:t}, d_{0:t}\}$.

Conceptually, patterning information has a directional flow: position → morphogen response → fate. Morphogen response encodes positional information, which is then interpreted to specify fate; fate is therefore the downstream readout rather than a predictor of position.

Accordingly, although the following quantity can be written in multiple notations,

$FI(x = d_{t'}) = I(d_{t'}; F) = PI_{t'}(g = F)$,

we refer to it as fate information carried by position, not "positional information carried by fate".

### Supplementary Note 2 | Mutual information estimation

#### a. Preprocessing

##### i. Constant cell number per stage

In our dataset, we captured dividing cells and tracked both daughters after division. Although retaining both daughters did not qualitatively change the dynamics of positional information $PI(g_t)$ (Extended Data Fig. 4m), we adopted a one-lineage-per-parent convention because several key quantities are cross-temporal, such as $PI_{t'}(g_t)$ and $PP_{t'}(d_t)$.

When a lineage *i* divides between stages $t'$ and $t$ (with $t' < t$), cross-temporal estimation requires relating a single parent value $x_{i,t'}$ to values at $t$. If both daughters $i_1$ and $i_2$ are retained, then the same parent observation $x_{i,t'}$ must be paired with both $y_{i_1,t}$ and $y_{i_2,t}$, effectively duplicating that parent value in the dataset. This duplication induces dependence between samples and can bias mutual-information estimates.

We therefore retain only one daughter branch after each division, so that each parent state contributes at most one observation to any cross-temporal estimate. This yields a more conservative sampling scheme that preserves as much independence as possible between observations.

##### ii. Within-embryo rank normalization and pooled estimation across embryos

Because lineage-resolved datasets were acquired embryo-by-embryo across multiple clutches and imaging days over two years, embryo-to-embryo variation can arise from biological differences and imaging conditions. To mitigate nuisance variation in fluorescence scaling and geometric scale, we rank-normalized, for each embryo $emb$ and each stage $t$, both Shh response and DV position to linearly spaced percentiles in $[0,1]$.

Ranks were computed within the set of lineage-traced cells in each embryo. Because lineages were reconstructed continuously over the full imaging window, the number of tracked lineages in embryo $emb$ is constant across stages; we denote this number by $n_{emb}$. For a variable $x$ measured for lineage $i$ at stage $t$ in embryo $emb$, we replaced each value by its ordinal rank among all lineages in that embryo at that stage, divided by $n_{emb}$:

$$x_{i,t,emb}^{\text{norm}} = \frac{\text{rank}_{t,emb}\left(x_{i,t,emb}^{\text{obs}}\right)}{n_{emb}}.$$

The resulting rank-normalized quantities are denoted $g_t$ for Shh response and $d_t$ for DV position.

After normalization, we pooled lineages across embryos within each condition to form empirical datasets for mutual-information estimation across stages. Throughout this note, N denotes the number of embryos pooled for an estimate and n denotes the corresponding number of pooled lineages.

#### b. Binning estimator

##### i. Quantile-binned plugin mutual information (naive estimator)

For a fixed condition and stage, we start from paired pooled observations $\{(X_i, Y_i)\}_{i=1}^{n}$, where each pair corresponds to one sampled lineage at that stage (under the one-lineage-per-ancestor convention described above). Here $i$ indexes sampled lineages in the pooled dataset, and $n$ is the number of pooled lineage observations used for that estimate. When stage indexing matters, we write variables explicitly as $g_{i,t}$ and $d_{i,t}$; in this subsection we suppress $t$ because the estimator is applied at a single fixed stage at a time.

We discretize each continuous variable into $Q$ equipopulated bins (quantile binning), yielding discrete pairs $\left(\widehat{X}_i, \widehat{Y}_i\right)$. From the set of these pairs, we build a $Q \times Q$ contingency table with counts $T_{xy}$ and empirical probabilities $\hat{p}_{xy} = T_{xy}/n$, from which we derive a plug-in ("naïve", see debiasing below) mutual-information estimate (in bits):

$$\hat{I}_{\text{naive}}\left(\hat{X}; \hat{Y}\right) = \sum_{x,y} \hat{p}_{xy} \left(\frac{\hat{p}_{xy}}{\hat{p}_x \hat{p}_y}\right).$$

When one variable is discrete (for example, a fate label $F$), we directly use its native categories and quantile-bin only the continuous variable.

##### ii. Direct (debiased) estimator via subsampling–extrapolation

Plug-in estimates are upward biased at finite sample size. We therefore implemented a debiased ("direct") estimate to extrapolate to the infinite-sample limit $1/n \to 0$, using a subsampling–extrapolation procedure inspired by Slonim et al[13].

For $c = 10$ Monte Carlo chains, we randomly permuted the $n$ paired observations and formed nested subsamples of sizes

$$\{n_j = \text{round}(f_j n)\}_j,$$

with fractions $f_j \in [f_{\text{min}}, 1]$ (we used $f_{\text{min}} = 0.5$), evenly spaced in $1/f$ (in practice, we used $1/f \in [1, 1.111, 1.222 \dots\ 2]$). For each chain, we computed $\hat{I}_{\text{naive}}(n_j)$ for each subsample size $n_j$and fitted a linear dependence on inverse sample size,

$$\hat{I}_{\text{naive}}(n_j) \approx a_0 + a_1(1/n_j).$$

The debiased estimate for that chain is the intercept $a_0$, corresponding to an extrapolation to infinite sample size. We report

$$\hat{I}_{\text{direct}} = \langle a_0 \rangle$$

as the mean across chains. An example extrapolation is shown in Extended Data Fig. 4a.

To quantify uncertainty in this extrapolation, we compute the standard deviation across chain intercepts, $SD_{\text{chains}}$, and report a 95% confidence interval

$$\hat{I}_{\text{direct}} \pm t_{0.975,\, c-1}\, SD_{\text{chains}}.$$

Using simulation benchmarks with known ground-truth mutual information and matched sample sizes/binning settings, this CI construction achieved ~95% empirical coverage of the true mutual information. We therefore report $\hat{I}_{\text{direct}}$ together with this 95% CI as a practical, empirically calibrated summary of uncertainty with respect to the infinite-sample extrapolation.

##### iii. Shuffle control and choice of bin-resolution

Using more bins increases discretization resolution and allows the estimator to capture finer dependence structure (green curves in Extended Data Fig. 4b,c). However, at fixed pooled sample size $n$, increasing $Q$ makes the $Q \times Q$ contingency table sparser and increases finite-sample upward bias, so that mutual-information estimates can remain positive even when the variables are statistically independent (grey curves in Extended Data Fig. 4b,c). We estimate this bias with a shuffle-based control alongside each mutual-information curve and choose the largest $Q$ for which the shuffle-null estimate is negligible, $Q_{\text{max}}(n)$, preventing discretization choices from introducing a spurious positive baseline.

Starting from pooled paired observations $\{(X_i, Y_i)\}_{i=1}^{n}$, we destroy dependence by randomly permuting one variable across cells while keeping the other fixed. Let π be a random permutation of $\{1, \dots, n\}$, and define shuffled pairs $\{(X_i,\ Y_{\pi(i)})\}_{i=1}^{n}$. We then compute mutual information on the shuffled dataset using the same binning resolution $Q$ and the same direct estimator used for the empirical estimate. We denote the resulting null estimate by $\hat{I}_{\text{shuffle}}$ and plot it as a reference baseline beneath the corresponding empirical curve where applicable (Fig. 1h,i; Extended Data Fig. 4b,c).

To choose $Q$ conservatively when pooled cell numbers differ across conditions, we precomputed a lookup table $n_{\min}(Q)$ (Extended Data Fig. 4c), defined as the minimum sample size for which the shuffle-null mutual-information estimate is not significantly greater than zero under matched estimator settings. For each candidate $Q$ and sample size $n$, we generated 30 independent synthetic datasets with $X \perp Y$ by drawing $X$ and $Y$ independently from $\text{Uniform}(0,1)$, computed the direct estimator on each replicate, and performed a one-tailed one-sample *t*-test of whether the mean shuffle-null estimate exceeds zero at $\alpha = 0.05$. We used a one-tailed test because the practical concern is an upward, strictly positive estimator baseline under independence.

We used this lookup table $n_{\min}(Q)$ to make sure to perform all our analyses above the corresponding bias-control threshold (Extended Data Fig. 4d), choosing the following values for the bin number: $Q = 10$ for per-embryo estimates (Fig. 1i and Fig. 6d); $Q = 30$ for wild-type pooled analyses with sufficient sampling (Figs. 1h and 2); $Q = 18$ for morphogenesis-perturbed and Notch-perturbed datasets (Figs. 3 and 4), with wild-type curves recomputed using $Q = 18$ for direct comparison; and $Q = 17$ for fate-mapped wild-type data (Fig. 5).

Within a given plot, mutual-information estimates are all compared using a single shared value of $Q$.

##### c. kNN estimator[14]

As a complementary method that avoids discretization, we also estimated mutual information using k-nearest-neighbour (kNN) estimator implemented in NPEET (https://github.com/gregversteeg/NPEET; continuous–continuous: ee.mi; mixed discrete–continuous: ee.micd), with its default parametrization.

#### Supplementary Note 3 | Estimator benchmarking and parameter sensitivity

Here we benchmark how estimator settings affect mutual-information estimates under the empirical distributions and sample sizes used in this study. The goal is to verify that the qualitative conclusions in the main text are not artefacts of binning resolution, pooled sample size, or estimator choice, and to motivate the practical comparability rules used across figures.

##### a. High-resolution empirical reference for benchmarking

To obtain an empirical, high-resolution reference, we pooled all traced wild-type cells ($N = 9$ embryos; $n = 1{,}561$ cells). This combined dataset supports high-resolution quantile binning at

$Q$ = 30 under our bias-control criteria (Extended Data Fig. 4b). Using this dataset, we computed the same set of information-theoretic quantities used throughout the manuscript—positional information and positional persistence—evaluated across integer somite stages, yielding 71 matched points spanning the observed dynamic range. We treat the $Q = 30$ binned estimates on the $n = 1{,}561$ dataset as an empirical reference for testing whether alternative settings preserve (1) the absolute mutual-information scale, quantified by the slope of a regression through the origin (Extended Data Fig. 4f,k, blue curves), and (2) the relative mutual-information relationship across the 71 matched estimates, quantified by the corresponding $R^2$ (Extended Data Fig. 4f,k, orange curves).

#### b. Effect of bin number $Q$ at fixed high $n$

We first quantified how reducing binning resolution changes mutual-information estimates while holding the underlying dataset and all definitions fixed (Extended Data Fig. 4e, illustrated for $Q = 10$ and $Q = 18$). Reducing $Q$ primarily compresses the mutual-information scale (slope <1), while preserving the relative relationships across the 71 matched estimates (high $R^2$) (Extended Data Fig. 4f). Increasing $Q$ improves absolute agreement with the $Q = 30$ reference, consistent with binning acting as an explicit resolution parameter that can be tightened when sample size permits (Extended Data Fig. 4b,c).

#### c. kNN estimator compared to the high-$Q$ binned reference

We next compared a continuous k-nearest-neighbour (kNN) estimator to the same $Q = 30$ binned reference using the $n = 1{,}561$ pooled wild-type dataset. The kNN estimate (illustrated for k = 3) closely matches the binned reference across the same 71 points (Extended Data Fig. 4g), indicating these two estimators converge when sample size supports high-resolution estimation.

We then scanned $k$ to test estimator sensitivity (Extended Data Fig. 4h). Agreement remains high across a broad range of $k$; yet the overall scale varies with k in a non-monotonic manner, emphasizing that $k$ is not a direct analogue of the "resolution" parameter $Q$ used in quantile binning (Extended Data Fig. 4h).

#### d. Dependence on embryo pooling $N$

Because we pooled embryos for most analyses, we quantified how mutual-information estimates depend on the number of embryos pooled ($N$) and the resulting pooled cell number ($n$). Using wild-type embryos, we constructed pooled datasets at multiple intermediate values of $N$, recomputed mutual-information estimates, and compared them to the $N = 9$ pooled wild-type reference. We show a representative pooled-versus-per-embryo comparison at $Q = 10$ (Extended Data Fig. 4i) and summarize their agreement as a function of $N$ (Extended Data Fig. 4j).

When $N \geq 3$ embryos, fitted slopes are close to the $N = 9$ reference and agreement is high ($R^2 > 0.95$). Accordingly, we repeated each perturbation experiment on at least three embryos for comparisons to wild type.

We repeated the same embryo-subsampling analysis using the kNN estimator. Again, agreement improves as embryos are pooled (Extended Data Fig. 4k,l).

##### e. Estimator choice in the main figures

Although the binning estimator and the k-nearest-neighbour estimator agree well in practice—both in the relative ordering of mutual-information values across stages and quantity types, and reasonably in absolute scale under our sample sizes (Extended Data Fig. 4e–h)—we use the binning estimator for most positional-information and positional-persistence curves in the main figures for three practical reasons.

First, the bin number $Q$ is an explicit and auditable resolution parameter that forces a controlled discretization and can be held fixed within a plot, enabling like-for-like comparisons across conditions even when pooled cell numbers differ. Second, our shuffle-null control procedure provides a direct check for spurious bias in the estimate, which we used to screen the possible values of $Q$ (Extended Data Fig. 4b–d). Third, the subsampling–extrapolation debiasing yields an empirical uncertainty on the infinite-sample extrapolation, reported as confidence intervals on the debiased values (Extended Data Fig. 4a). We do not have an comparably interpretable uncertainty estimate for the kNN estimator under these settings.

##### f. When we use kNN in this study

We use kNN estimates for two purposes: (1) to cross-check our binning estimates without variable discretization under matched distributions and sample sizes (Extended Data Fig. 4g,h,k,l), and (2) to evaluate mutual information between multivariate quantities, without discretization-imposed ceilings (Extended Data Fig. 3p-r, when considering multivariate references for DV position).

Our binning estimator suffers from the curse of dimensionality. Discretizing a $p$-dimensional variable into $Q$ bins yields $Q^p$ joint states: controlling finite-sample bias at a given accuracy therefore requires sample size $n$ to scale with $Q^p$. With finite $n$, as in this study, multivariate binning requires reducing $Q$ substantially, which imposes a discretization ceiling on estimated mutual information and can skew conclusions.

#### Supplementary Note 4 | Choice of Shh response and DV position metric

In this study, we quantify positional information as mutual information between a Shh-response variable $g$ and a DV-position variable $d$. Both variables can be defined in multiple biologically reasonable ways (for example, using different reporters or different distance metrics), and these choices affect the value of $I(g; d)$. We therefore aimed to adopt definitions that are simple, biologically interpretable, and capture the strongest measurable dependence between Shh response and DV position in our data. We also prioritized definitions that keep $g$ and $d$ low-dimensional, because when variables become multivariate the binning estimator quickly becomes impractical at our sample sizes (Supplementary Note 3f). We therefore only add dimensions if they provide substantial information gains.

#### a. Shh response, *g*

We considered two Shh signalling reporters: an endogenous-regulatory reporter, *TgBAC(ptch2:Kaede)*[15], and a synthetic multimerized Gli-binding-site reporter, *Tg(8xgli-Xla.Cryaa:NLS-mCherry)* (GBS)[16]. We compared these reporters based on (1) their DV fluorescence dynamic range, which determines the DV span that can be robustly sampled, and (2) the mutual information between reporter signal and DV position, which quantifies how well each reporter predicts DV position in our information-theoretic analyses.

At 10 ss, *ptch2:Kaede* showed a broader DV-dependent range than the GBS reporter (Extended Data Fig. 3a–f), consistent with a longer fitted decay length (Extended Data Fig. 3g), and carried higher mutual information with DV position (Extended Data Fig. 3h). Consistent with these protein-level comparisons, reporter mRNA measurements also supported a more spatially extended *ptch2:Kaede* signal relative to *GBS: NLS-mCherry* over the same period (Extended Data Fig. 3i–k). We therefore used *ptch2:Kaede* as the primary Shh-response variable throughout the study.

#### b. DV position, *d*

##### i. Centroid distance versus nuclear-surface distance

As a convenient output from Imaris, we initially used the shortest 3D distance from the nuclear surface to the notochord surface when benchmarking Shh reporter choice. We then asked whether a centroid-to-surface distance metric, computed in Python because it is not exported directly by Imaris, would improve positional-information estimates. The two distance definitions were highly similar and yielded near-identical positional information across stages (Extended Data Fig. 3n), making the choice between them inconsequential for our conclusions. We therefore used the nuclear-surface–to–notochord-surface distance as our default DV-position metric.

##### ii. Choice of reference structure: notochord versus floor plate

Because both the notochord and the floor plate secrete Shh, the tissue used as a reference to define DV position could, in principle, affect how well Shh response specifies position. We therefore compared DV coordinates referenced to the floor plate versus the notochord, using the criterion that the most relevant reference should maximize positional information carried by Shh response. Notochord-referenced distance consistently captured substantial positional information, whereas floor-plate–referenced distance did not (Extended Data Fig. 3o). We also tested whether using both coordinates jointly increased positional information, but observed no substantial gain relative to notochord distance alone (Extended Data Fig. 3p). We therefore defined $d$ as notochord-referenced distance throughout the study.

##### iii. Testing additional spatial covariates: laterality and anteroposterior position

A central result of this study is that positional information, positional persistence, and fate information can be placed on the same information-theoretic scale and converge to a shared ~1.2-bit bound. This interpretation assumes that the relevant spatial structure in *ptch2:Kaede* levels is along the DV axis, so that $d$ captures the main positional variable that the Shh response encodes. We therefore tested whether augmenting $d$ with additional spatial coordinates—left–right position or anteroposterior position—captured additional information about the spatial distribution of *ptch2:Kaede*. These covariates produced, at most, small gains (Extended Data Fig. 3q,r) in positional information, indicating that notochord-referenced DV position is the dominant geometric predictor of Shh response in this dataset. We therefore did not include laterality or anteroposterior position in the primary definition of $d$.

### Supplementary Note 5 | Decoding-based analyses

#### a. Continuous decoding of DV position from Shh-response history

To quantify how precisely DV position can be inferred from a cell's full Shh-response history, we aimed to estimate the trajectory-based positional information,

$$PI_{t'}(g_{0:t}) = I(g_{0:t};\, d_{t'}),$$

using the notation in Supplementary Note 1. However, $g_{0:t}$ is a high-dimensional time series, making direct mutual-information estimation impractical at our sample sizes due to the curse of dimensionality. We therefore adopt a decoding-based approach[17]: we train supervised regression models to predict DV position from $g_{0:t}$, and then compute the mutual information between decoded positions and measured positions. This provides a tractable, interpretable estimate of how much positional information can be extracted from Shh response histories.

##### i. Inputs and targets

Let $g_{i,t}$ denote the Shh response of lineage $i$ at stage $t$. For each stage $t$, we represent lineage $i$ by its Shh-response history up to $t$,

$$x_{i,t} = g_{i,0:t}.$$

The corresponding regressor matrix $X_t$ has one row per lineage and one column per included time point. We trained regression models directly on these high-dimensional histories, without dimensionality reduction, to predict DV position $d$; the resulting predictions are denoted $\hat{d}_{i,t}$.

Depending on the analysis, the positional target was $d \in \{d_t,\, d_0,\, d_{12}\}$, corresponding to instantaneous, initial, or final DV position (Extended Data Fig. 5a–e).

##### ii. Decoding-based mutual information estimation

Decoders were trained using outer 5-fold cross-validation across lineages. For each fold, the model was trained on a subset of training lineages and used to predict $d$ for held-out lineages. We pooled predictions across folds to obtain a paired dataset $\{(\hat{d}_i, d_i)\}_{i=1}^{n}$, and computed decoding-based positional information as

$$I_{dec}(g_{0:t}; d) := I\big(\hat{d}(g_{0:t});\, d\big),$$

using the same binning estimator and shuffle control as in Supplementary Note 2. To keep $I(\hat{d}(g_{0:t});\ d)$ directly comparable to the corresponding snapshot estimate $I(g_t; d)$, we used the same bin number $Q$ as in the matched snapshot analysis.

**iii. Decoder benchmarking and selection**

We benchmarked multiple regression models spanning linear shrinkage and nonlinear approaches (ridge, lasso, elastic net, random forest, gradient boosting, kNN and SVR-RBF). Across stages and targets, ridge regression provided the largest decoding-based information under cross-validation (Extended Data Fig. 5f–h) and was therefore retained for the position-decoding analyses shown in the main figures (Fig. 1h-i, Fig. 2g, Fig. 5f-g).

#### b. Decoding of discrete fate labels from continuous histories

To test whether histories up to stage $t$ support fate assignment, we decoded ventral fate at 12 ss, $F_{12}$, from lineage histories of Shh response ($g_{0:t}$), DV position ($d_{0:t}$), or the combination of both ($g_{0:t}, d_{0:t}$) (Extended Data Fig. 5k–o). Because the target is a discrete fate category rather than a continuous position, we used a classification model rather than a regression model.

**i. Inputs and targets**

For each lineage $i$, we constructed a history vector $x_{i,t}$ from $x \in \{g,\ d,\ (g, d)\}$, sampled up to stage $t$, yielding a regressor matrix $X_t$ with one row per lineage and one column per included time point. Models were trained directly on these high-dimensional histories, without dimensionality reduction, to predict the fate label $F_i$. The corresponding predicted labels are denoted $\hat{F}_i$.

**ii. Cross-validation and decoding-based fate information**

As for decoding positions, we used 5-fold cross-validation across lineages. We pooled the predictions for held-out lineages across the folds to obtain a single set of decoded labels $\{\hat{F}_{12,i}(t)\}_{i=1}^{n}$ matched to true labels $\{F_{12,i}\}_{i=1}^{n}$. We quantified the decoding-based fate information as

$$I_{dec}(x_{0:t}; F) := I\big(\hat{F}_{12}(t)\ ;\ F_{12}\big),$$

computed from the 4 × 4 contingency table of decoded versus true fate labels (Extended Data Fig. 5n,o). Finite-sample bias was corrected using the same subsampling–extrapolation procedure as in Supplementary Note 2.

**iii. Classifier benchmarking and selection**

We benchmarked several classifiers (random forest, logistic regression with L1/L2 penalties, linear SVM, kNN and Gaussian naive Bayes). Random forest achieved the highest decoding fate information and was therefore used for discrete fate decoding (Extended Data Fig. 5k–o).

#### c. Comparing snapshot and decoding estimates

Continuous histories (for example, $d_{0:t}$ or $g_{0:t}$) contain more information in general than their snapshot counterparts (for example, $d_t$ or $g_t$), In particular, for true mutual information values,

$$I(g_{0:t};\, d)\ \geq I(g_t;\, d).$$

However, direct estimation of $I(g_{0:t}; d)$ is not tractable because $g_{0:t}$ is high-dimensional. We therefore use decoding to map the history $g_{0:t}$ to a one-dimensional prediction $\hat{d}$ (or a predicted label $\hat{F}$) and then estimate $I(\hat{d}; d)$ (or $I(\hat{F}; F)$). This dimensionality reduction introduces an information bottleneck: $\hat{d}$ or $\hat{F}$ is not guaranteed to preserve all information about the target that is present in the full history. Consequently, decoding-based information estimates are not guaranteed to exceed snapshot estimates. If $I_{\text{dec}} > I$, the history contains additional information beyond the snapshot and the decoder can access it. If $I_{\text{dec}} < I$, the comparison is ambiguous because decoder limitations could hide information present in the history (Fig 5f,g).

### Supplementary Note 6 | Breaking down the temporal evolution of $\boldsymbol{PI_t}$: a local $\boldsymbol{\Delta PI}$ analysis

To separate changes in positional information driven by changes in Shh response from changes driven by changes in DV position, we analysed how cross-temporal positional information varies locally around each developmental stage. Cross-temporal positional information is computed as in Supplementary Note 2 and follows the notation in Supplementary Note 1:

$$PI_{t'}(g_t) = I(g_t;\, d_{t'}),$$

where $g_t$ is the Shh response measured at stage $t$, and $d_{t'}$ is the DV position measured at stage $t'$. Conceptually, $PI_{t'}(g_t)$ quantifies how well the Shh response sampled at stage $t$ predicts the DV position measured at stage $t'$.

It is useful to view $PI_{t'}(g_t)$ as a function of the two variables $(t, t')$, defined on a 2D plane. Local changes near the diagonal ($t = t'$) can then be decomposed into two directional contributions, corresponding to updating either the Shh-response measurement (along the $t$ axis) or the DV-position target (along the $t'$ axis), while holding the other variable fixed (Fig. 2c).

#### a. Local contributions around the diagonal

We carry out this decomposition focusing on the diagonal $t' = t$, which corresponds to the instantaneous positional information

$$PI_t(g_t) = I(g_t;\, d_t).$$

##### i. Shh-response contribution, $\boldsymbol{\Delta PI(g_t)}$

To capture how positional information changes when the Shh response is sampled slightly earlier or later than $t$ for prediction, while the DV position being predicted is held at the anchor stage $t$, we define:

$\Delta PI_g(t) \approx \frac{\partial}{\partial t} PI_{t'}(g_t)|_{t'=t}$ (see also Extended Data Fig. 5p,q).

Intuitively, $\Delta PI_g(t)$ captures the local benefit (or cost, when $< 0$) of using slightly later Shh response to predict position at stage $t$.

##### ii. DV-position contribution, $\boldsymbol{\Delta PI(d_t)}$

Similarly, to capture how positional information changes when the DV position to predict is evaluated slightly later or earlier than $t$, while the Shh response sample is held at the anchor stage $t$, we define:

$\Delta PI_d(t) \approx \frac{\partial}{\partial t'} PI_{t'}(g_t)|_{t'=t}$ (see also Extended Data Fig. 5p,r),

These two quantities correspond to the components illustrated in Fig. 2c and quantified in Fig. 2d. We detail in the next section how we evaluated them.

#### b. Numerical estimation used in this study

We estimate $\Delta PI(g_t)$ and $\Delta PI(d_t)$ from one-dimensional slices through the surface $PI_{t'}(g_t) = I(g_t;\, d_{t'})$, first applying Gaussian smoothing and then estimating the local derivative using a centred finite difference (method visualized in Extended Data Fig. 5p–t).

For each anchor stage $t$:

i. To estimate $\boldsymbol{\Delta PI(g_t)}$, we take the slice $\boldsymbol{PI_t(g_\tau)}$ as a function of $\boldsymbol{\tau}$ with $\boldsymbol{t}$ fixed, smooth this slice with a Gaussian filter with $\boldsymbol{\sigma = 0.5}$ ss, and then evaluate the slope at $\boldsymbol{\tau = t}$ with the central difference formula, with step $\boldsymbol{\delta = 0.04}$ ss

$$\boldsymbol{\Delta PI(g_t) \approx \frac{PI_t^{sm}(g_{t+\delta}) - PI_t^{sm}(g_{t-\delta})}{2\delta}}.$$

ii. To estimate $\boldsymbol{\Delta PI(d_t)}$, we take the slice $\boldsymbol{PI_\tau(g_t)}$ as a function of $\boldsymbol{\tau}$ with $\boldsymbol{t}$ fixed, smooth this slice with the same Gaussian filter, and then compute the slope at $\boldsymbol{\tau = t}$ with the same formula:

$$\boldsymbol{\Delta PI(d_t) \approx \frac{PI_{t+\delta}^{sm}(g_t) - PI_{t-\delta}^{sm}(g_t)}{2\delta}}.$$

Slopes have units of bits per ss. The sign convention matches Fig. 2d and Extended Data Fig. 5q–r, where $\Delta PI(g_t)$ is typically positive early and $\Delta PI(d_t)$ is typically negative.

#### c. Stage-binned summaries and variability display

To summarize trends over time, we bin slope estimates by integer stage $T$ using anchor stages $t \in [T - 0.5,\, T + 0.5)$. For each bin, and for both contributions, we compute the mean and 95% CI, consistent with the display in Fig. 2d.

#### d. Validity of the approximation of the signal as the sum of both contributions

Finite differences provide only an approximation to derivatives, and the resulting slopes need not integrate back exactly to the full signal over long times. In practice, $PI_{t'}(g_t)$ is sufficiently smooth that summing the contributions $\Delta PI(g_t)$ and $\Delta PI(d_t)$ over time closely tracks the observed profile $PI_t(g_t)$, reproducing the rise-and-fall structure with only minor drift (Extended Data Fig. 5t). This supports the usefulness of the decomposition not only locally, but also across the full experiment: it provides an intuitive accounting of how Shh-response-driven increases and DV-position-driven decreases combine over time to generate the rise-and-fall structure.

### Supplementary Note 7 | Morphogenesis-normalized time $\tilde{t}$

Across embryos and perturbations, morphogenetic progression can be faster or slower. To compare information-theoretic quantities on a common "morphogenesis clock," we define a morphogenesis-normalized stage $\tilde{t}$ by applying a per-embryo linear rescaling of somite stage (Extended Data Fig. 6e–h).

For each embryo $emb$, let $\bar{d}_{emb}(t)$ denote the embryo-averaged raw position at stage $t$. We define the embryo's position midpoint

$$\bar{d}_{emb}^{\mathrm{mid}} = \frac{1}{2}\left(\min_t \bar{d}_{emb}(t) + \max_t \bar{d}_{emb}(t)\right),$$

and define the embryo's centre stage $t_{emb}^{\mathrm{centre}}$ as the stage at which $\bar{d}_{emb}(t)$ is closest to $\bar{d}_{emb}^{\mathrm{mid}}$.

We then define the WT reference centre stage $t_{\mathrm{WT}}^{\mathrm{centre}}$ as the mean of $t_{emb}^{\mathrm{centre}}$ across WT embryos, and compute one normalization factor for each embryo:

$$m_{emb} = \frac{t_{\mathrm{WT}}^{\mathrm{centre}}}{t_{emb}^{\mathrm{centre}}}.$$

Finally, we map each original stage $t$ (ss) to morphogenesis-normalized stage

$$\tilde{t} = m_{emb}\, t.$$

This rescaling aligns embryos by forcing their morphogenetic midpoints to occur at the same $\tilde{t}$, while preserving the ordering of stages for each embryo.

Because sampling is uniform in $t$ but not in $\tilde{t}$ after this embryo-specific rescaling, we resampled each lineage's measurements onto a uniform grid in normalized time ($\Delta\tilde{t} = 0.04$ somite stages) using linear interpolation, and recomputed all quantities as in the main analysis.

### Supplementary Note 8 | Hybrid models

To test whether morphogenetic DV-position dynamics alone are sufficient to set the timing at which Shh response best encodes DV position, we developed numerical "hybrid" models that combine (1) a static Shh ligand field $c(d)$, defined as a function of absolute DV distance $d$ (μm), with (2) a fixed, cell-autonomous response module that maps ligand exposure into a cumulative Shh-response readout. We call these models “hybrid” because the ligand field and response module are idealized and held constant over time, whereas the DV-position trajectories are provided as independent inputs. The signalling part (ligand field and response module) was fit on wild-type data and then held fixed across conditions, while DV-position trajectories were taken directly from lineage-traced measurements and used embryo-by-embryo.

By varying only the cell trajectories, while keeping the ligand field and response rule fixed, these models isolate the contribution of morphogenesis to the Shh-response dynamics and the timing of information transfer, providing a controlled sufficiency test for the changes we observed in mutants (Fig. 3f–g; Extended Data Fig. 7).

For most simulations, the only geometric input is each lineage’s DV position over time (absolute distance to the notochord, µm). For the Notch/lateral-inhibition extension, we additionally used each cell’s full 3D coordinates $(x, y, z)$ to recompute nearest-neighbour relationships at each time point, couple Notch dynamics between neighbouring cells, and use the resulting Notch state to modulate each cell’s Shh responsivity (Fig. 4g). This extension provides a separate sufficiency test for whether Notch-mediated heterogeneity can account for the lack of late accumulation of positional information (Fig. 4h).

Implementation details are available at:
https://git.ista.ac.at/jrenaud/pi_while_morphogenesis_methods.

#### a. Inputs: DV position and Shh response in absolute units

For each lineage $i$, the models take as input the measured DV-position trajectory $d_{t,i}$ and output a simulated Shh response $g_{t,i}$, intended to match the experimentally measured Shh response. Throughout this note, $g$ refers to the raw cumulative readout (no rank normalization), and $d$ refers to DV position, expressed in absolute units (µm).

#### b. Fixed Shh ligand field

Ligand availability is represented by a minimal exponential Shh profile along the DV axis,

$$\mathrm{Shh}(d) = \exp(-d/D),$$

with $D$ a decay length (µm). For each lineage, ligand exposure is evaluated along the measured DV-position trajectory as $\mathrm{Shh}\left(d_{i,t}\right)$. Because the ligand field is fixed across embryos and conditions, variability in $\mathrm{Shh}\left(d_{i,t}\right)$ arises only from differences in DV trajectories.

#### c. Benchmarking candidate response modules

We benchmarked multiple plausible response modules that convert ligand exposure into a cumulative reporter-like readout, and then selected a model using a generalization-first criterion focused on the morphogenesis-perturbed condition.

##### i. Optimization criterion (used during fitting)

All candidate models were fit on wild-type data by minimizing a trajectory-matching loss computed as root-mean-square error in log space between measured and modelled $g_{i,t}$, pooled over lineages. This log-RMS loss is the objective optimized during parameter search.

##### ii. Generality and model selection criterion (used after fitting)

To evaluate generality in a way aligned with downstream information-theoretic analyses, we assessed performance using a secondary loss metric alongside the fitting loss, comparing modelled versus measured trajectories after converting each embryo's values to within-embryo percentiles. This emphasizes within-embryo structure (the scale used for mutual-information estimation) rather than absolute fluorescence scaling.

For the choice of the final model, we prioritized the model with the smallest within-embryo percentile loss in *MZitga5*$^{-/-}$ embryos, the condition where generalization is most challenging (Extended Data Fig. 7a).

#### d. Selected response model: Hill sensing with a two-layer accumulation cascade

For each lineage $i$, the model evaluates ligand exposure along the measured DV trajectory as $\mathrm{Shh}(d_{i,t})$ and converts it into a bounded activation drive $u_{i,t}$ using Hill sensing,

$$u_{i,t} = \frac{\mathrm{Shh}(d_{i,t})}{K + \mathrm{Shh}(d_{i,t})},$$

where K is a half-activation constant. The Hill coefficient is degenerate with respect to the decay length of the gradient, so we fix it to 1. The predicted cumulative Shh-response readout is generated by a two-layer linear cascade that implements temporal filtering and:

$$\frac{dv_{i,t}}{dt} = u_{i,t} - \beta_1\, v_{i,t},$$

$$\frac{dg_{i,t}}{dt} = \left(\alpha\, v_{i,t} - \beta_2\, g_{i,t}\right)(1 + \xi),$$

where $v_{i,t}$ is an intermediate response state and $g_{i,t}$ is the predicted cumulative output. Parameter $\alpha$ is an amplification constant, $\beta_1$ and $\beta_2$ are decay timescales with $\beta_2 \ll \beta_1$ as a result of the fitting process, and $\xi$ a multiplicative Gaussian noise term. Simulated $g_{i,t}$ was processed with the same downstream steps used for mutual-information estimation as the ones measured *in vivo*.

#### e. Fit-once strategy and hybrid predictions across conditions

All free parameters of the selected model, including the Shh decay length D and the sensing/cascade parameters, were fit once using wild-type lineage data. The deterministic parameters $\alpha$, $\beta_1$, and $\beta_2$ were fit jointly to minimize the trajectory-matching loss defined above. The amplitude of the noise term $\xi$ was then chosen to match the mutual-information scale between model and experiment, because the deterministic model alone overestimates information values (compare Extended Data Fig. 7i,j with Fig. 2f). All fitted parameters were subsequently held fixed when generating predictions for morphogenesis-perturbed mutants and other conditions. This single-fit design ensures that genotype-specific changes in predicted Shh-response dynamics arise from substituting the measured DV-position trajectories $d_{i,t}$, rather than from re-optimizing either the Shh ligand field or the response rule (Fig. 3f–g; Extended Data Fig. 7a,g).

#### f. Notch-inspired lateral inhibition simulation

To test whether Delta–Notch coupling can generate cell-to-cell heterogeneity in Shh sensitivity and thereby prevent late-time accumulation of positional information, we implemented a Notch-inspired lateral inhibition module in which cells interact locally with their neighbours. The resulting Notch state was then used to modulate the Shh-response module in each cell by scaling the activation gain α (Extended Data Fig. 7f,h,k).

##### i. Notch state update rule and neighbourhood coupling

We assigned each cell an initial Notch value close to Notch=1, with small random variation. Notch states were then iteratively updated using a minimal model of lateral inhibition[18,19], which has two stable equilibrium states at Notch=0 and Notch=1 and drives segregation into a salt-and-pepper pattern. At each update, each cell interacts with its nearest neighbours, which are re-identified at every time point to reflect cell rearrangement during tissue remodelling. The inhibition strength and the number of neighbours could be varied within a reasonable range without changing the overall statistics of the resulting Notch patterns. By contrast, the timescale of this Notch module, and the time at which it begins to modulate the Shh response (as described below), were tuned to reproduce the late-time saturation of the wild-type information curves that the previous models did not capture, while preserving the early-time behaviour that those models captured better (Fig. 3e,g).

##### ii. Coupling Notch state to Shh response simulation

At each time step, the Notch state modulates the amplification parameter α in the Shh-response cascade, ranging from 0 when $Notch = 0$ to α when $Notch = 1$. This procedure progressively introduces cell-to-cell variation over time while preserving the fixed Shh field and the same underlying response module, enabling the comparisons shown in Extended Data Fig. 7f,h,k.

#### g. Using model-predicted Shh response for information-theoretic analyses

After simulating $g_{i,t}$ from the measured $d_{i,t}$, we computed the same positional information and cross-temporal quantities on model outputs as on experimental data, using identical estimators and preprocessing conventions. This includes positional information about instantaneous, initial, and final DV position, enabling direct comparison of model-predicted and measured information dynamics (Fig. 3g; Extended Data Fig. 7).

### Supplementary Note 9 | Shh sensitivity score and Notch correlations

#### h. Sensitivity-score definition

To quantify cell-to-cell variation in instantaneous Shh response beyond what is explained by DV position, we defined a per-cell Shh sensitivity score using within-embryo percentile normalization (Extended Data Fig. 8b). For each embryo, we converted (1) *ptch2* mRNA intensity (used here as an instantaneous Shh response), $g^{\text{inst}}$, (2) DV distance to the notochord, $d$, and (3) a candidate gene's expression level, $x$, into within-embryo percentiles in $[0,1]$: $\text{pct}(g^{\text{inst}})$, $\text{pct}(d)$, and $\text{pct}(x)$. We then defined a ventral coordinate $v = 1 - \text{pct}(d)$, so that larger $v$ corresponds to more ventral positions. Shh sensitivity was defined as a rank-residual of *ptch2* relative to DV position,

$$s = \text{pct}(g^{\text{inst}}) - (1 - \text{pct}(d)) = \text{pct}(g^{\text{inst}}) - v.$$

Thus, $s > 0$ indicates cells with higher *ptch2* mRNA than expected for their DV position, whereas $s < 0$ indicates the opposite (Extended Data Fig. 8b).

#### i. Correlation analysis with candidate pathways

To test whether candidate pathways covary with Shh sensitivity, we computed, for each embryo and each gene $x$, the Spearman correlation between the sensitivity score and the within-embryo expression percentile of that gene:

$$\rho = \text{cor}_{\text{Spearman}}(s,\ \text{pct}(x)).$$

This yields one correlation coefficient per embryo for each gene. Positive $\rho$ indicates that higher expression of $x$ is associated with higher Shh sensitivity (higher *ptch2* than expected at a given DV position), whereas negative $\rho$ indicates association with reduced sensitivity.

We then tested whether correlations were consistently nonzero across embryos using a one-sample *t*-test on embryo-level $\rho$ values (null hypothesis: mean $\rho = 0$). Representative relationships are shown for a Notch-associated progenitor marker (*her12*) and a neurogenic marker (*elavl3*) (Extended Data Fig. 8c), alongside DV–LR expression maps for the Notch-associated targets used in this analysis (Extended Data Fig. 8d; Supplementary Note 10).

### Supplementary Note 10 | Construction of DV–LR expression profiles from pixel-resolved tissue masks

To visualize spatial gene-expression patterns across the spinal cord while minimizing embryo-to-embryo differences in tissue size and orientation, we computed two-dimensional dorsoventral (DV)–left/right (LR) profiles from pixel-resolved volumes.

#### a. Tissue masks and analysis domain

For each embryo, we obtained binary masks for spinal cord, somites and notochord from either ilastik object predictions or deep-learning segmentation (Supplementary Methods 1; Extended Data Fig. 1). All downstream analyses were performed on all voxels within the spinal cord mask; voxels outside the spinal cord were excluded.

#### b. Pixel-wise distance transforms

For each spinal-cord voxel, we computed (1) the shortest Euclidean distance to the notochord mask (DV-relevant distance) and (2) the shortest Euclidean distance to the left and right somite masks (LR-relevant distances). Somites were separated into left and right components by connected-component analysis: we retained the two largest components and assigned “left” and “right” by their centroid x-coordinates. Distances were computed using Euclidean distance transforms, yielding a per-voxel notochord distance $d_{\text{noto}}$ and left/right somite distances $d_L$ and $d_R$.

#### c. Within-embryo normalization of DV and LR coordinates

Because absolute distances vary with embryo size and staging, we normalized spatial coordinates within each embryo. DV position was defined as the within-embryo percentile of $d_{\text{noto}}$ among all spinal-cord voxels, producing a DV coordinate in [0,1] (0 = most ventral/closest to notochord; 1 = most dorsal/farthest). LR position was defined from somite distances as a signed laterality index

$$LR = \frac{d_L - d_R}{d_L + d_R},$$

and then scaled to place it in approximately [-0.5,0.5] by dividing by 2. This LR coordinate is 0 at the midline, negative on the left and positive on the right (after the left/right assignment described above).

#### d. Binning on a common DV–LR grid and averaging

To obtain comparable spatial maps across embryos, we discretized the normalized DV and LR coordinates onto a fixed 2D grid with spacing 0.02 in both axes. For each voxel, we assigned a DV bin and LR bin by rounding to the nearest 0.02 grid point. We then averaged pixel intensities within each $(DV, LR)$ bin, first within embryos and then across embryos by pooling all spinal-cord voxels (equivalently, averaging over voxels from all embryos at matched DV–LR bins). For visualization, we restricted profiles to the interior range $DV \in (0.05, 0.95)$ and $LR \in (-0.45, 0.45)$ to avoid edge bins with sparse occupancy. For each marker channel (for example, endogenous *ptch2* mRNA or Notch-associated targets), we constructed DV–LR heatmaps from the binned mean intensity values.

### Supplementary Note 11 | Quantifications for fate configuration, entropy and fate information panels

#### a. Overview and datasets

This note describes the quantifications used for (1) fate configuration (cell-type fractions and mean dorsoventral positions), (2) fate-distribution entropy, and (3) fate information carried by dorsoventral position in fixed-embryo datasets and in timed Shh-inhibition experiments (Fig. 6a–d; Extended Data Fig. 9). Analyses were performed per embryo ($N$ embryos per group). Embryo-level summaries are shown as points and group summaries as mean ± s.d. unless noted.

#### b. Fate-marker staining and definition of ventral progenitor classes

Embryos were fixed and stained by multiplex RNA HCR for ventral progenitor markers to assign ventral classes at the analysis stage (10 ss for timed inhibition; 12 ss for fate mapping after live imaging). We used marker sets for p3 (*nkx2.2a*, *nkx2.2b*, *nkx2.9*), pMN (*olig2*), p2 (*lbx2*), and p1 (*prdm12b*), and in some datasets additionally p0 (*dbx1a*, *dbx1b*) to quantify broader ventral composition (Fig. 6a–c; Extended Data Fig. 9c–d). Nuclei were detected within the spinal-cord volume and each nucleus was assigned a fate label based on the multi-channel marker pattern (Fig. 5a; Extended Data Fig. 9h). For fixed specimens, DV position for each nucleus was quantified as the shortest 3D spot-to-surface distance to the notochord surface (Fig. 6b; Extended Data Fig. 9i–j).

For fate-information and entropy calculations, we defined the p3–p1 population as nuclei assigned to p3, pMN, p2, or p1. We excluded p0 and dorsal/unspecified nuclei to match the cell types covered by the lineage-tracing dataset.

#### c. Cell-type fractions, fate-distribution entropy, and fate information

For each embryo, we restricted analyses to the p3–p1 population with $F \in \{p3, pMN, p2, p1\}$. Fate fractions were computed as

$$p_F = \frac{n_F}{\sum_{f \in \{p3, pMN, p2, p1\}} n_f},$$

where $n_F$ is the number of nuclei assigned to fate $F$ in that embryo. We quantified fate-distribution entropy as

$$S(F) = -\sum_F p_F \log_2 p_F,$$

which satisfies $S(F) \leq 2$ bits for four classes, with equality only for equal fractions (Fig. 6c; Extended Data Fig. 9f,g).

For each embryo, we then computed fate information carried by DV position, $FI(d)$, using the same mutual-information framework as in Supplementary Notes 1–2 with the binning estimator and Q=10 for all embryo-level comparisons (Fig. 6d; Extended Data Fig. 9g).

#### d. Fate information carried by DV position

For each embryo, we computed fate information, $FI(d)$, restricting to the $F \in \{p3, pMN, p2, p1\}$. Fate information was estimated exactly as in Supplementary Notes 1–2 with Q=10 (Fig. 6d; Extended Data Fig. 9g).

#### e. Composition-matched Control baseline by resampling

##### i. Necessity for a Control baseline

Fate information can be written as

$$FI(d) = I(F; d) = S(F) - \langle S(F \mid d) \rangle_d,$$

so it depends both on the overall fate composition $S(F)$ and on how sharply fate is specified at a given DV position through $\langle S(F \mid d) \rangle_d$. To separate changes in $FI(d)$ driven purely by changes in fate composition from changes in the DV position–fate relationship, we defined, for each embryo, a **composition-matched Control baseline** that preserves the Control position–fate mapping while matching the embryo's observed fate proportions.

##### ii. Evaluation procedure

For each embryo $emb$ in an inhibition condition, let $n_{emb}$ be the number of p3–p1 cells and let $\{p_{F,emb}\}$ be its observed fate fractions over $F \in \{p3, pMN, p2, p1\}$. We generated synthetic datasets of size $n_{emb}$ by resampling cells from the pooled Control dataset while enforcing the same fate fractions $\{p_{F,emb}\}$; DV positions were taken from the corresponding resampled Control cells. For each resample, we computed $FI(d)$ using the same estimator settings as for real embryos. We summarized the baseline for embryo $emb$ as the mean $FI(d)$ across resamples and plotted it as the black-square reference in Fig. 6d.

##### iii. Results interpretation

This composition-matched Control baseline is the fate information expected from changes in fate composition alone, assuming the conditional entropy $\langle S(F \mid d) \rangle_d$ is unchanged. Values below this baseline would indicate a degraded DV position–fate mapping beyond composition effects. In our experiments, inhibition-condition embryos match the composition-matched baseline, indicating that changes in fate composition account for most of the observed differences in $FI(d)$ (Fig. 6d). Consistently, the ratio $FI(d)/S(F)$ remains approximately constant across inhibition conditions (Extended Data Fig. 9g). Expected and measured Control means are not identical because $FI(d)$ estimates show modest dependence on pooled sample size and estimator settings (Extended Data Fig. 4j,l).

#### f. Within-domain DV-position spread

To test whether timed Shh inhibition broadened DV positioning within each ventral fate domain, we quantified within-domain spread as the standard deviation of DV position across cells of a given fate within each embryo (Extended Data Fig. 9i–j). We then tested for a condition effect on within-domain spread using an embryo-level ordinary least squares (OLS) regression that controls for fate identity and covariates expected to influence spread:

$$SD(d) \sim C(\text{Condition}) + C(F) + \text{mean}(d) + p_F.$$

Here, $\text{mean}(d)$ is the embryo-by-fate mean DV position and $p_F$ is the embryo-by-fate fraction (included because spread can vary with domain location and effective sample size).

After adjusting for domain mean position, domain size (fraction), and fate identity, inhibition window showed no evidence of increased within-domain DV-position spread (Type II ANOVA for Condition, $P = 0.081$). The largest deviation was a trend toward reduced spread in the 5–10 hpf window ($\beta = -0.289$, $P = 0.055$), whereas the 10–12 hpf and 12–14 hpf windows were indistinguishable from control ($P = 0.741$ and 0.904).

### Supplementary Note 12 | scRNA-seq analysis

We analysed ZMAP, an integrated zebrafish single-cell RNA-seq resource assembled from multiple studies. To unbiasedly analyse the spinal cord population without biased stage distribution in limited stages and cells, our strategy is to analyse the spinal cord population (ZMAP_Tissue = spinal_cord) from a broad developmental stage (3-24 hpf) and keep the desired stage (8-16 hpf) maintaining the same neighbours, classification, UMAP etc after analysis.

In detail, we subset spinal cord cells from 3–24 hpf, then re-clustered the subset using the precomputed neighbour graph provided in the ZMAP AnnData object and Leiden clustering (resolution = 3).

We assigned clustered with ventral identities using established marker combinations[20]:

- Floor plate (FP): *shha*, *shhb*, *foxa1*
- p3: *nkx2.2a*, *nkx2.2b*, *nkx2.9*
- pMN: *olig2*
- p2: *nkx6.1*/*nkx6.2*/*irx3a* positive and *olig2* negative
- p1: *prdm12b*
- p0: *dbx1a*, *dbx1b*

Notably, there are no established p2 markers in zebrafish spinal cord yet. For example, the mammalian p2 marker *Foxn4* is not expressed in zebrafish spinal cord at the defined stages. From sc-RNA seq analysis, within the p2-like population (defined by the transcription-factor combination above and its position between pMN and p1 in UMAP space), we identified *lbx2* and *si:ch211-152f23.5* as novel markers enriched in p2 cells.

After classification, we removed floor plate cells and restricted the analysis to 8–16 hpf, leaving us with 66 libraries from 6 independent studies[21-26]. For each stage, we then computed the fraction of each ventral progenitor identity relative to total spinal cord cells (floor plate excluded). Because ZMAP pools studies with heterogeneous sampling (including potential differences in anterior–posterior representation and dissection depth across timepoints), we interpreted the timing of ventral-state emergence as the primary readout, rather than treating absolute cell-type fractions as directly comparable across stages.

### Supplementary Note 13 | Amplitude analysis and signal post-processing

We measured several dorsoventrally patterned signals over time and required a simple, robust summary of how their overall magnitude changes during development. For each embryo and time point, we therefore fit a low-parameter DV-gradient model to single-cell measurements as a function of DV position and extracted the fitted amplitude parameter $A(t)$. This analysis is used to visualize and compare signal magnitudes across time and embryos (including for hybrid-model analyses and plotting). It is not used as an input to positional-information calculations (see below).

#### a. Measurement model and background assumption

Let d denote DV position and let $x(d)$ denote the measured fluorescence or expression value in a cell at position $d$. We assume the measurement can be decomposed into (1) a biological signal that varies with DV position, $f(d)$, and (2) a uniform background $B$ that is spatially constant across the imaged volume for a given embryo and time point. Thus, for each embryo and time point,

$$x(d) \approx f(d) + B.$$

Here, $B$ captures camera offset, diffuse background, and autofluorescence. Because autofluorescence can bleach, $B$ may vary over time; however, within any single image acquisition $B$ is assumed uniform across DV position. In practice, we estimate B jointly with the DV-gradient parameters so that the baseline signal is explicitly separated from the patterned component.

#### b. DV-gradient functions

We used two simple families of DV-gradient functions $f(d)$, chosen to match the qualitative profiles observed in the data.

##### i. Gaussian profile

For markers whose expression is broadly distributed and centred around a preferred DV position, we used a Gaussian:

$$f(d) = A \cdot exp\left(-\frac{(d-\mu)^2}{2\sigma^2}\right),$$

where $A$ is the amplitude, $\mu$ is the centre, and $\sigma$ is the width.

**ii. Exponential profile**

For *ptch2* and other Shh-response readouts, we used an exponential decay from ventral to dorsal:

$$f(d) = A \cdot e^{-d/\lambda},$$

where $A$ is the amplitude and $\lambda$ is the decay length.

#### c. Fitting in log space

Our measurements span orders of magnitude, and both high- and low-expression regions are informative. On the raw scale, measurement noise is typically heteroscedastic: high-expression regions show larger absolute variance than low-expression regions. Least-squares fitting on the raw scale therefore overweights high-expression regions and can degrade fits in the low-expression tail. To reduce this imbalance, we fit in log space by minimizing squared residuals between log-transformed measurements and a log-transformed model:

$$\log x\left(d_{i,t}\right) \approx \log\left(f\left(d_{i,t}\right) + B_{t,emb}\right),$$

jointly across all cells $i$ in a given embryo $emb$ at that time point $t$.

#### d. Intensity rescaling for single-cell visualization and hybrid models

Imaging conditions can vary across experiments, introducing embryo-to-embryo differences in measured fluorescence intensity on top of biological variation. To reduce this nuisance variation and place measurements on a common scale for visualization and for hybrid-model fitting, we applied two steps, separately for each embryo and time point.

First, we background-corrected each cell's measurement using the fitted background term:

$$x_{corr,i,t} = x_{i,t} - B_{t,emb},$$

Second, we rescaled corrected measurements by an embryo-specific constant $C_{emb}$ (defined in the amplitude-normalization procedure below):

$$x_{\mathrm{corr},i,t} = x_{\mathrm{corr},i,t}/C_{emb}.$$

These adjusted values were used for plotting and for training/running the hybrid models, treating embryos on a common intensity scale.

#### e. Amplitude normalization for visualization

To visualize amplitude dynamics on a common 0–1 scale across different plots, we also applied two optional rescaling methods to the fitted amplitude $A(t)$:

**i. Within-embryo normalization:**

For each embryo $emb$, define $C_{emb} = \max_t A(t)$ and plot

$A_{\mathrm{corr}}(t) = \frac{A(t)}{C_{emb}},$

so that $\max_t A_{\mathrm{corr}}(t) = 1.$

**ii. Across-embryo reference normalization**

Define a single shared factor

$C_{\mathrm{ref}} = \max_t \langle A(t) \rangle_{\mathrm{emb}},\ \langle A(t) \rangle_{\mathrm{emb}} = \frac{1}{N} \sum_{emb=1}^{N} A_{emb}(t).$

and plot

$A_{\mathrm{corr}}(t) = \frac{A(t)}{C_{\mathrm{ref}}},$

so that the maximum of the normalized embryo-mean amplitude equals 1 (Extended Data Fig. 1j–m; Extended Data Fig. 10d).

These normalizations are used for visualization only and preserve relative changes within each curve.

#### f. Relation to the values used for mutual information estimates

Mutual-information estimates were computed after within-embryo rank normalization performed separately at each stage (Supplementary Notes 1–2). The background subtraction and positive rescaling described here were not used for mutual-information estimation. Hybrid-model outputs were analysed using the same downstream preprocessing and estimators as the experimental measurements, so any global rescaling in the modelled signals is handled consistently.
